# Monocyte control of organismal energy homeostasis

**DOI:** 10.1101/2025.02.20.639373

**Authors:** Rui Martins, Birte Blankehaus, Faouzi Braza, Pedro Ventura, Sumnima Singh, Sebastian Weis, Miguel Mesquita, Maria Pires, Sara Pagnotta, Qian Wu, Sílvia Cardoso, Elisa Jentho, Ana Figueiredo, Pedro Faísca, Ana Nóvoa, Vanessa Alexandra Morais, Stefanie K. Wculek, David Sancho, Moises Mallo, Miguel P. Soares

## Abstract

Multicellular organisms rely on inter-organ communication networks to maintain vital parameters within a dynamic physiological range. Macrophages are central to this homeostatic control system, sensing deviations of those parameters and responding accordingly to support tissue function and organismal homeostasis. Here we demonstrate that dysregulation of iron metabolism in parenchyma cells, imposed by the deletion of ferritin H chain, is sensed by monocyte-derived macrophages. In response, macrophages derived from circulating monocytes support tissue function, energy metabolism and thermoregulation, as demonstrated in bone marrow chimeric and parabiotic mice. This salutary effect is contingent on a transcriptional program, controlled in macrophages by the transcription factor A mitochondria. This transcriptional response acts in a non-cell autonomous manner to support the mitochondria of parenchyma cells, irrespectively of mitochondrial transfer. In conclusion, monocyte-derived macrophages cross-regulate Fe and energy metabolism to support tissue function and organismal homeostasis.

## INTRODUCTION

Iron (Fe) was co-opted to exchange electrons with acceptor (electrophile) and donor (nucleophile) molecules in vital biochemical processes, such as those supporting mitochondrial function and energy metabolism (Muchowska *et al*, 2019; Teh *et al*, 2024; Wade *et al*, 2021). Presumably as an evolutionary trade-off (Stearns & Medzhitov, 2015), Fe can catalyze in a unfettered manner the production of hydroxyl radicals (HO^-^) and other reactive oxygen species (ROS), when exchanging electrons with superoxide (O^•-2^) or hydrogen peroxide (H^2^ O^2^) (Winterbourn, 1995).

Cellular ROS accumulation can catalyze lipid peroxidation and trigger programmed cell death via ferroptosis (Dixon *et al*, 2012). The fitness costs associated with uncontrolled Fe redox activity are restrained by a number of evolutionarily conserved Fe-regulatory genes (Galy *et al*, 2023). These include ferritin h chain (FTH), a master regulator of Fe redox activity and bioavailability (Blankenhaus *et al*, 2019; Galy *et al*., 2023; Gozzelino & Soares, 2014; Harrison & Arosio, 1996).

Organismal Fe homeostasis relies on an inter-organ communication network encompassing Fe-recycling macrophages that express high levels of FTH (Galy *et al*., 2023). Similar to other tissue-resident macrophages lineages (Ginhoux *et al*, 2010), Fe-recycling macrophages develop from yolk sac progenitors (Kohyama *et al*, 2009) but can be replaced throughout adult life by monocyte-derived macrophages (van Furth & Diesselhoff-den Dulk, 1984) that develop from bone marrow (BM) hematopoietic progenitors (van Furth & Cohn 1968).

Monocytes differentiate into tissue resident macrophages (Guilliams *et al*, 2018) via specific transcriptional and epigenetic programs (Gosselin *et al*, 2014; Lavin *et al*, 2014), activated in response to tissue-specific cues (Amit *et al*, 2016; Okabe & Medzhitov, 2015). Heme, a Fe-containing protoporphyrin used as prosthetic group of hemoglobin, induces the genetic program supporting the development of Fe-recycling macrophages (Haldar *et al*, 2014).

Macrophages establish functional interactions with parenchyma cells in all tissues, early through embryonic development (Lazarov *et al*, 2023) and throughout post-natal life (Nobs & Kopf, 2021; Okabe & Medzhitov, 2015; Zhou *et al*, 2018a). This intimate bond “supports” the core effector functions of “primary” parenchyma cells, underlying tissue function (Adler *et al*, 2023; Meizlish *et al*, 2021; Zhou *et al*, 2018b). Under this conceptual framework, tissues can be perceived as an emerging property arising from the interactions of “supportive” and “primary” cells, where macrophages act as universal “supportive” cells (Adler *et al*., 2023; Meizlish *et al*., 2021; Zhou *et al*., 2018b).

Regulation of Fe metabolism by FTH exerts control over energy metabolism in adult mice (Blankenhaus *et al*., 2019). Considering the impact of Fe metabolism on macrophage function (Soares & Hamza, 2016) and energy homeostasis (Joffin *et al*, 2022), we asked whether the expression of FTH in macrophages impacts on organismal energy homeostasis. We found that monocyte-derived macrophages respond to dysregulation of Fe metabolism in parenchyma cells via an FTH-dependent mechanism that controls the activation of a transcriptional program supporting the mitochondria of parenchyma cells. This transcriptional program regulated in macrophage by the mitochondrial transcription factor A (TFAM) is essential to restore organismal energy balance in mice lacking FTH in parenchyma cells. We infer that monocyte-derived macrophages operate as a central component of an inter-organ surveillance system that cross-regulates Fe and energy metabolism to support organismal homeostasis.

## RESULTS

### *Fth*-competent hematopoietic cells rescue chimeric *Fth*-deleted mice

Having established that regulation of Fe metabolism by FTH exerts a critical role in the control energy metabolism in adult mice (Blankenhaus *et al*., 2019), we sought to determine the relative contribution of FTH expression in hematopoietic *vs.* parenchyma (*i.e.,* non-hematopoietic) to energy homeostasis. To this end we used *Fth^R26fl/fl^* mice allowing for global *Fth*-deletion (*i.e., Fth^R26^*^Δ*/*Δ^) in response to tamoxifen (TAM) administration and *Fth^wt/wt^* or *Fth^fl/fl^* mice as controls (Blankenhaus *et al*., 2019). *Fth^R26fl/fl^* and control *Fth^wt/wt^* or *Fth^fl/fl^* mice were lethally irradiated and transplanted with bone marrow (BM) cells from *Fth^R26fl/fl^ vs.* control *Fth^wt/wt^* or *Fth^fl/fl^* mice, to generate chimeric mice carrying an *Fth* deletion in hematopoietic *vs.* parenchyma cells, in response to TAM administration. We confirmed that BM engraftment was > 95%, as assessed by the relative proportion of donor *vs.* recipient CD45.1 *vs.* CD45.2 cells, respectively, 4 weeks after BM transplantation (*Fig. S1A-C*).

Systemic *Fth* deletion in hematopoietic and parenchyma cells from *Fth^R26^*^Δ*/*Δ^⇨*Fth^R26^*^Δ*/*Δ^ chimeras (*i.e., Fth^R26fl/fl^* mice reconstituted with *Fth^R26fl/fl^* BM) led to wasting (*Fig. 1A,B, S1D,E*), hypothermia (*Fig. 1A,C, S1D,F*) and death (*Fig. 1A,D, S1D,G*), within 10-40 days after TAM administration, consistent with global *Fth* deletion in adult *Fth^R26^*^Δ*/*Δ^ mice (Blankenhaus *et al*., 2019). Control *Fth*-competent *Fth^wt/wt^*⇨*Fth^fl/fl^* chimeric mice (*i.e., Fth^fl/fl^* mice reconstituted with *Fth^wt/wt^* BM) did not develop this lethal wasting syndrome, in response to TAM administration at the same dosage and schedule (*Fig. 1B-D, S1D-G*).

**Figure 1:**
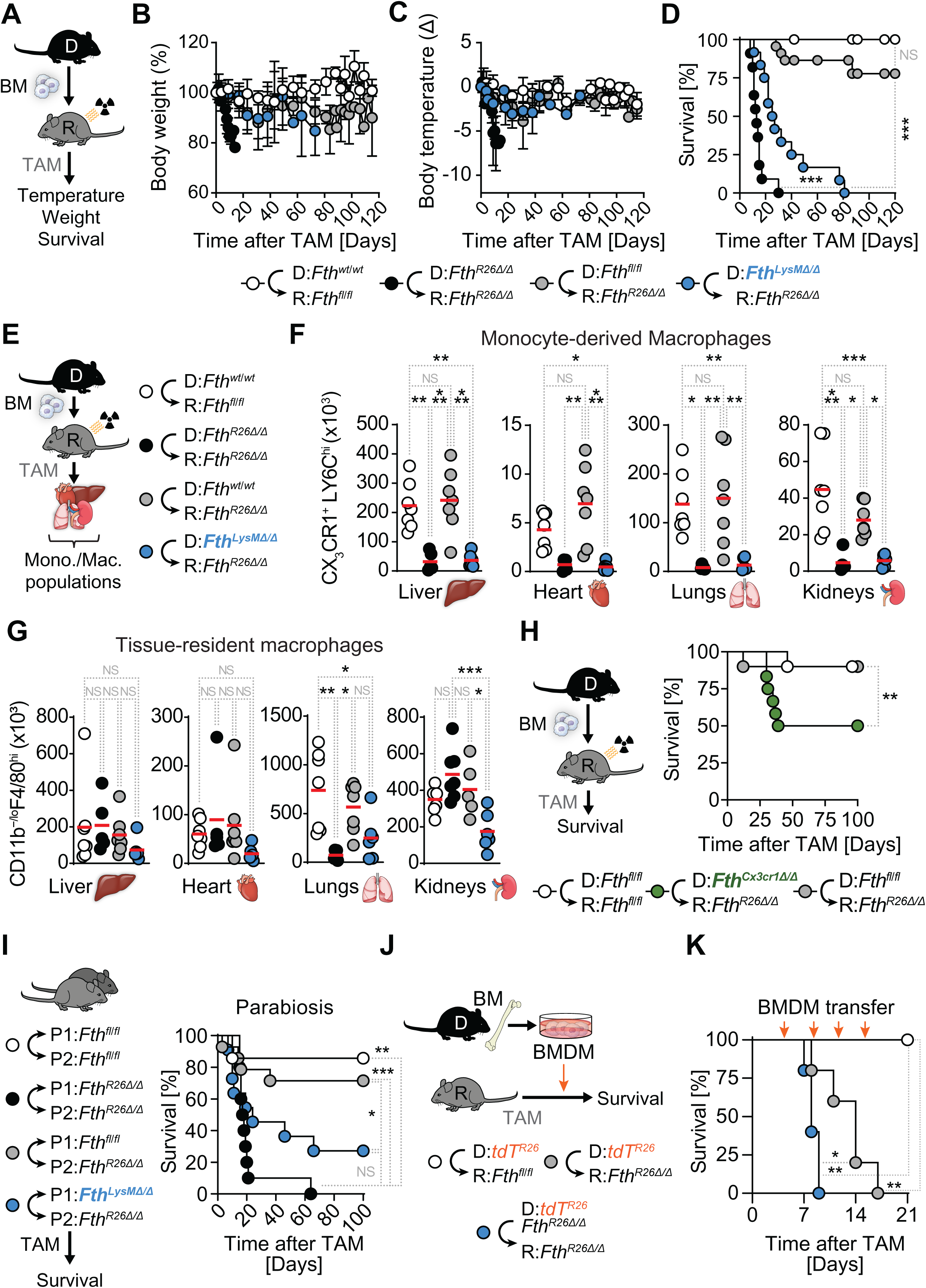
*Fth*-competent monocyte-derived macrophages rescue *Fth*-deleted mice. (**A**) Schematic representation of TAM-induced *Fth* deletion in chimeric mice (day 0) and vital parameters monitored. Relative (**B**) Body weight, (**C**) Temperature and (**D**) survival of *Fth^wt/wt^*⇨*Fth^fl/fl^*(n=7-8), *Fth^R26^*^Δ*/*Δ^⇨*Fth^R26^*^Δ*/*Δ^ (n=6), *Fth^wt/wt^*⇨*Fth^R26^*^Δ*/*Δ^ (n=10-21) and *Fth^LysM^*^Δ*/*Δ^⇨*Fth^R26^*^Δ*/*Δ^ (n=9-12) chimeric mice. Data in (B-D) is pooled from 6 independent experiments with similar trends. Data in (B, C) represented as mean ± SD. (**E**) Schematic representation of TAM-induced *Fth* deletion in chimeric mice (day 0) and flow cytometry analysis of monocyte/macrophage populations. (**F**) Absolute number of CX_3_CR1^+^LY6C^high^ monocyte-derived macrophages (backgated as Ly6G^−^CD11b^+^F4/80^low^) and (**G**) tissue-resident macrophages (Ly6G^−^CD11b^−/low^F4/80^high^) in the liver, heart, lungs and kidneys of *Fth^fl/fl^*⇨*Fth^fl/fl^* (n=7), *Fth^R26^*^Δ*/*Δ^⇨*Fth^R26^*^Δ*/*Δ^ (n=5), *Fth^fl/fl^*⇨*Fth^R26^*^Δ*/*Δ^ (n=7) and *Fth^LysM^*^Δ*/*Δ^ *Fth^R26^*^Δ*/*Δ^ (n=6) chimeric mice, 7 to 19 days post-TAM administration. Data in (F) presented as individual values (circles) and mean (red bars), pooled from 3 independent experiments with similar trends. (**H**) Schematic representation of TAM-induced *Fth* deletion (day 0) in chimeric mice, and survival of *Fth^fl/fl^*⇨*Fth^fl/fl^* (n=10), *Fth^Cx3cr1^*^Δ*/*Δ^⇨*Fth^R26^*^Δ*/*Δ^ (n=12) and *Fth^fl/fl^*⇨*Fth^R26^*^Δ*/*Δ^ (n=10) chimeric mice. Data in (H) pooled from 3 independent experiments with similar trends. (**I**) Schematic representation of parabiosis, TAM administration (day 0) and survival of parabiotic *Fth^fl/fl^*⇔*Fth^fl/fl^* (n=7 pairs), *Fth^R26^*^Δ*/*Δ^⇔*Fth^R26^*^Δ*/*Δ^ (n=10 pairs), *Fth^fl/fl^*⇔*Fth^R26^*^Δ*/*Δ^ (n=14 pairs) and *Fth^LysM^*^Δ*/*Δ^⇔*Fth^R26^*^Δ*/*Δ^ (n=11 pairs). Data in (I) pooled from 4 independent experiments with similar trends. (**J**) Schematic representation of adoptive transfer of *in vitro*-differentiated BMDMo into recipient mice, following TAM administration (day 0). (**K**) Survival of *Fth^fl/fl^* or *Fth^R26^*^Δ*/*Δ^ mice adoptively transferred with *Fth*-competent *tdT^R26^* or *Fth*-deleted *tdT^R26^Fth^R26^*^Δ*/*Δ^ BMDMo. Times of adoptive transfer indicated by orange arrows. Survival analysis was performed using Log-rank (Mantel-Cox) test. One-Way ANOVA with Tukey’s range test for multiple comparison correction was used for comparison between multiple groups. NS: non-significant, * P<0.05, ** P<0.01, *** P<0.001.

*Fth* deletion specifically in parenchyma cells (*i.e., Fth^wt/wt^*⇨*Fth^R26^*^Δ*/*Δ^ chimeras; *Fth^R26fl/fl^* mice reconstituted *Fth^wt/wt^* BM) did not cause the development of wasting (*Fig. 1B*), hypothermia (*Fig. 1C*) or lethality (*Fig. 1D*). Moreover, *Fth* deletion specifically in hematopoietic cells (*i.e., Fth^R26^*^Δ*/*Δ^⇨*Fth^fl/fl^*chimeras; *Fth^fl/fl^* mice reconstituted with *Fth^R26fl/fl^* BM) also did not cause wasting (*Fig. S1D,E*), hypothermia (*Fig. S1D,F*) nor mortality (*Fig. S1D,G*). These observations suggest that *Fth*-competent hematopoietic cells can rescue the lethal outcome of systemic *Fth* deletion in chimeric mice.

### *Fth*-competent myeloid cells rescue chimeric *Fth*-deleted mice

To establish whether myeloid cells are necessary to rescue the lethal outcome of *Fth* deletion in adult mice, we generated chimeric mice with BM cells from *Fth^LysM^*^Δ*/*Δ^ mice, carrying an *Fth* deletion specifically in myeloid cells (Ramos *et al*, 2019; Wu *et al*, 2023). *Fth*-deleted myeloid failed to prevent the development of wasting (*Fig. 1B*), hypothermia (*Fig. 1C*) and death in *Fth^LysM^*^Δ*/*Δ^⇨*Fth^R26^*^Δ*/*Δ^ chimeras (*i.e., Fth^R26fl/fl^* mice reconstituted with *Fth^LysM^*^Δ*/*Δ^ BM), with all chimeric mice succumbing within 80 days of TAM administration (*Fig. 1D*). This suggests that FTH expression in myeloid cells is essential to rescue the lethal outcome of global *Fth* deletion in chimeric mice.

Although *Fth^R26^*^Δ*/*Δ^⇨*Fth^R26^*^Δ*/*Δ^ and *Fth^LysM^*^Δ*/*Δ^⇨*Fth^R26^*^Δ*/*Δ^ chimeras succumbed to *Fth* deletion, the time dynamics differ slightly, with lethality of *Fth^R26^*^Δ*/*Δ^⇨*Fth^R26^*^Δ*/*Δ^ chimeras preceding that of *Fth^LysM^*^Δ*/*Δ^⇨*Fth^R26^*^Δ*/*Δ^ chimeras. As such, subsequent analyses were conducted at “early onset” and “late onset”, respectively, with corresponding control chimeras.

To establish whether *Fth*-competent lymphocytes are necessary to rescue the lethal metabolic collapse caused by systemic *Fth* deletion, we generated *Fth^Cd2^*^Δ*/*Δ^ mice, carrying an *Fth* deletion specifically in lymphocytes. *Fth* deletion in lymphocytes did not compromise the capacity of hematopoietic cells to prevent the development of wasting (*Fig. S1H,I*), hypothermia (*Fig. S1H,J*) nor lethality (*Fig. S1H,K*) in *Fth^Cd2^*^Δ*/*Δ^⇨*Fth^R26^*^Δ*/*Δ^ chimeras (*i.e., Fth^R26fl/fl^*mice reconstituted with *Fth^Cd2^*^Δ*/*Δ^ BM). This suggests that FTH expression in lymphocytes, the most abundant population of circulating leukocytes in adult mice, is not required to rescue the lethal outcome of *Fth* deletion in parenchyma cells.

### Tissue monocyte-derived macrophages depletion in chimeric *Fth*-deleted mice

Hematopoietic *Fth* deletion was associated with the depletion of monocyte-derived macrophages (Ly6G^−^CD11b^+^F4/80^low^) expressing C-X3-C Motif Chemokine Receptor 1 (CX_3_CR1) and high levels of Lymphocyte Antigen 6 Complex, Locus C (LY6C) (CX_3_CR1^+^LY6C^high^) in *Fth^R26^*^Δ*/*Δ^⇨*Fth^R26^*^Δ*/*Δ^ chimeras, as illustrated in the liver, heart, lungs and kidneys (*Fig. 1E,F; S2A*). In contrast, tissue-resident CD11b^−/low^F4/80^high^ macrophages were not affected, with the exception of the lungs (*Fig. 1E,G; S2B*). Moreover, monocyte-derived (*Fig. 1E,F; S2A*) and tissue-resident macrophages (*Fig. 1E,G; S2B*) were not depleted when *Fth* was deleted specifically in parenchyma cells from *Fth^wt/wt^*⇨*Fth^R26^*^Δ*/*Δ^ chimeras.

There was a marked reduction of the number of monocyte-derived macrophages when *Fth* was deleted in myeloid cells from *Fth^LysM^*^Δ*/*Δ^⇨*Fth^R26^*^Δ*/*Δ^ chimeras (*Fig. 1E,F; S2A*). This was not associated however with a reduction of tissue-resident macrophages, with the exception of the kidneys (*Fig. 1E,G; S2B*).

This suggests that when *Fth* is deleted in parenchyma cells, FTH expression becomes essential to support the tissue-dependent differentiation of CX_3_CR1^+^LY6C^high^ monocyte-derived macrophages (Guilliams *et al*., 2018; Trzebanski *et al*, 2024). Of note this is not observed when FTH expression is expressed in parenchyma cells from mice in which *Fth* is deleted specifically in myeloid cells (Bolisetty *et al*, 2015; Ikeda *et al*, 2020),

### *Fth*-competent monocyte-derived macrophages rescue chimeric *Fth*-deleted mice

Monocyte-derived macrophages support tissue function (Swirski *et al*, 2009), suggesting that classical monocytes (CX_3_CR1^+^LY6C^high^F4/80^-^) prevent the lethal outcome of global *Fth* deletion in chimeric mice. To test this hypothesis, we generated *Fth^Cx3cr1^*^Δ*/*Δ^ mice, carrying a *Fth* deletion under the control of the *Cx3cr1* promoter (Jung *et al*, 2000). *Fth*-deleted classical monocytes failed to rescue the lethal outcome of *Fth* deletion in 50% of *Fth^Cx3cr1^*^Δ*/*Δ^⇨*Fth^R26^*^Δ*/*Δ^ chimeras (*i.e., Fth^R26fl/fl^* mice reconstituted with *Fth^Cx3cr1^*^Δ*/*Δ^ BM) (*Fig. 1H*). This suggests that *Fth*-competent classical monocytes contribute are essential to support organismal homeostasis, when *Fth* is deleted in parenchyma cells.

Monocyte-derived macrophages migrate to non-lymphoid tissues via a mechanism that relies on the chemokine/chemokine receptor monocyte chemotactic protein 1 (MCP-1; CCL2) and C-C chemokine receptor type 2 (CCR2; CD192), respectively (Boring *et al*, 1997). To test whether the CCL2-CCR2 chemotactic axis is required to support the salutary effects of monocyte-derived macrophages, we reconstituted *Fth^R26fl/fl^* mice with BM from *Ccr2* deficient *Ccr2*^−*/*−^ mice (Boring *et al*., 1997). *Fth* deletion in *Ccr2*^−*/*−^⇨*Fth^R26^*^Δ*/*Δ^ chimeras was only partially lethal, compared to control *Ccr2^+/+^*⇨*Fth^fl/fl^*chimeras (*Fig. S2C*). This suggests that monocyte-derived macrophages support organismal homeostasis via a mechanism that is only partially CCL2/CCR2-dependent.

### *Fth*-competent circulating monocytes rescue chimeric *Fth*-deleted mice

To test whether circulating monocytes carry the capacity to rescue the lethal outcome of systemic *Fth* deletion we used parabiosis, an experimental approach whereby a common circulatory system is established surgically to enable the transit of circulation cells (and soluble factors) between two mice (Harris, 2013). Global *Fth* deletion caused severe wasting (*Fig. S2D*) and was lethal (*Fig. 1I*) in parabiotic *Fth^R26^*^Δ*/*Δ^↔*Fth^R26^*^Δ*/*Δ^ pairs. Expression of FTH in one parabiotic mouse restored the survival of the *Fth*-deleted parabiotic mouse (*Fig. 1I*), similar to control *Fth*-competent parabiotic pairs, receiving TAM at the same dosage and schedule (*Fig. 1I, S2D*). This suggests that *Fth*-competent hematopoietic-derived circulating cells and/or soluble factors generated in a mouse that expresses FTH are sufficient to rescue the lethal outcome of systemic *Fth* deletion.

The protective effect of *Fth*-competent circulating cells was contingent on the expression of FTH in myeloid cells, as revealed by the lethal outcome of *Fth* deletion in *Fth^LysM^*^Δ*/*Δ^↔*Fth^R26^*^Δ*/*Δ^ parabiotic pairs (*Fig. 1I, S2D*). This suggests that circulating *Fth*-competent myeloid cells, which include circulating monocytes, are required to prevent the lethal outcome of systemic *Fth* deletion, in the absence of lethal irradiation and/or hematopoietic cell reconstitution.

### *Fth*-competent BM-derived monocytes (BMDMo) rescue chimeric *Fth*-deleted mice

To probe unequivocally that *Fth*-competent monocytes carry the capacity to rescue the lethal outcome of global *Fth* deletion we generated BM-derived monocytes (BMDMo) *in vitro* and tested their protective effect upon adoptive transfer into adult *Fth*-deleted *Fth^R26^*^Δ*/*Δ^ mice. To trace BMDMo, we added a *Rosa26*-tandem dimer (td) Tomato-Flox-stop-Flox allele driving the expression of td Tomato (tdT) by Cre-driven excision, under the control of *Rosa26* promoter (*tdT^R26^* mice). Adoptive transfer of *Fth*-competent tdT^+^ BMDMo (generated from *tdT^R26^* mice) was sufficient to extend the lifespan of *Fth*-deleted *Fth^R26^*^Δ*/*Δ^ mice (*Fig. 1J*). In contrast, Adoptive transfer of *Fth*-deleted tdT^+^ BMDMo (generated from *tdT^R26^Fth^R26^*^Δ*/*Δ^ *^l^* mice) failed to extend the lifespan of *Fth*-deleted *Fth^R26^*^Δ*/*Δ^ mice (*Fig. 1J*). All recipient *Fth*-deleted *Fth^R26^*^Δ*/*Δ^ mice succumbed within 1-2 days after the last BMDMo adoptive transfer, suggesting that long-term survival of *Fth^R26^*^Δ*/*Δ^ mice is dependent on continuous supply of circulating monocytes, presumably via myelopoiesis.

### *Fth*-competent myeloid cells restore Fe homeostasis in chimeric *Fth*-deleted mice

The lethal outcome of global *Fth* deletion was associated with systemic dysregulation of Fe metabolism, as illustrated by reduced circulating transferrin and higher transferrin saturation in *Fth*-deleted *Fth^R26^*^Δ*/*Δ^⇨*Fth^R26^*^Δ*/*Δ^ *vs. Fth*-competent *Fth^wt/wt^*⇨*Fth^fl/fl^*chimeras (*Fig. 2A-C*). *Fth*-competent hematopoietic cells restored the levels of circulating transferrin and normalized transferrin saturation (*Fig. 2A-C*) in plasma from *Fth^fl/fl^*⇨*Fth^R26^*^Δ*/*Δ^ chimeras to levels in the range of those in control *Fth* competent *Fth^fl/fl^*⇨*Fth^fl/fl^*chimeras (*Fig. 2A-C*). This was not the case however, when *Fth* was deleted in myeloid cells from *Fth^LysM^*^Δ*/*Δ^⇨*Fth^R26^*^Δ*/*Δ^ chimeras, which presented a decrease in the levels of circulating transferrin and an increase in transferrin saturation (*Fig. 2A-C*), as compared to *Fth* competent *Fth^fl/fl^*⇨*Fth^fl/fl^*chimeras (*Fig. 2C*). This was associated, over time, with an increase in the concentration of circulating Fe in both *Fth^LysM^*^Δ*/*Δ^⇨*Fth^R26^*^Δ*/*Δ^ and *Fth^fl/fl^*⇨*Fth^R26^*^Δ*/*Δ^ chimeras (*Fig. 2D,E*), suggesting that, over time, *Fth*-competent monocyte-derived macrophages do not prevent the accumulation of circulating redox-active Fe imposed by global *Fth* deletion, and that this increase in circulatory Fe is independent from *Fth* deletion-induced lethality.

**Figure 2:**
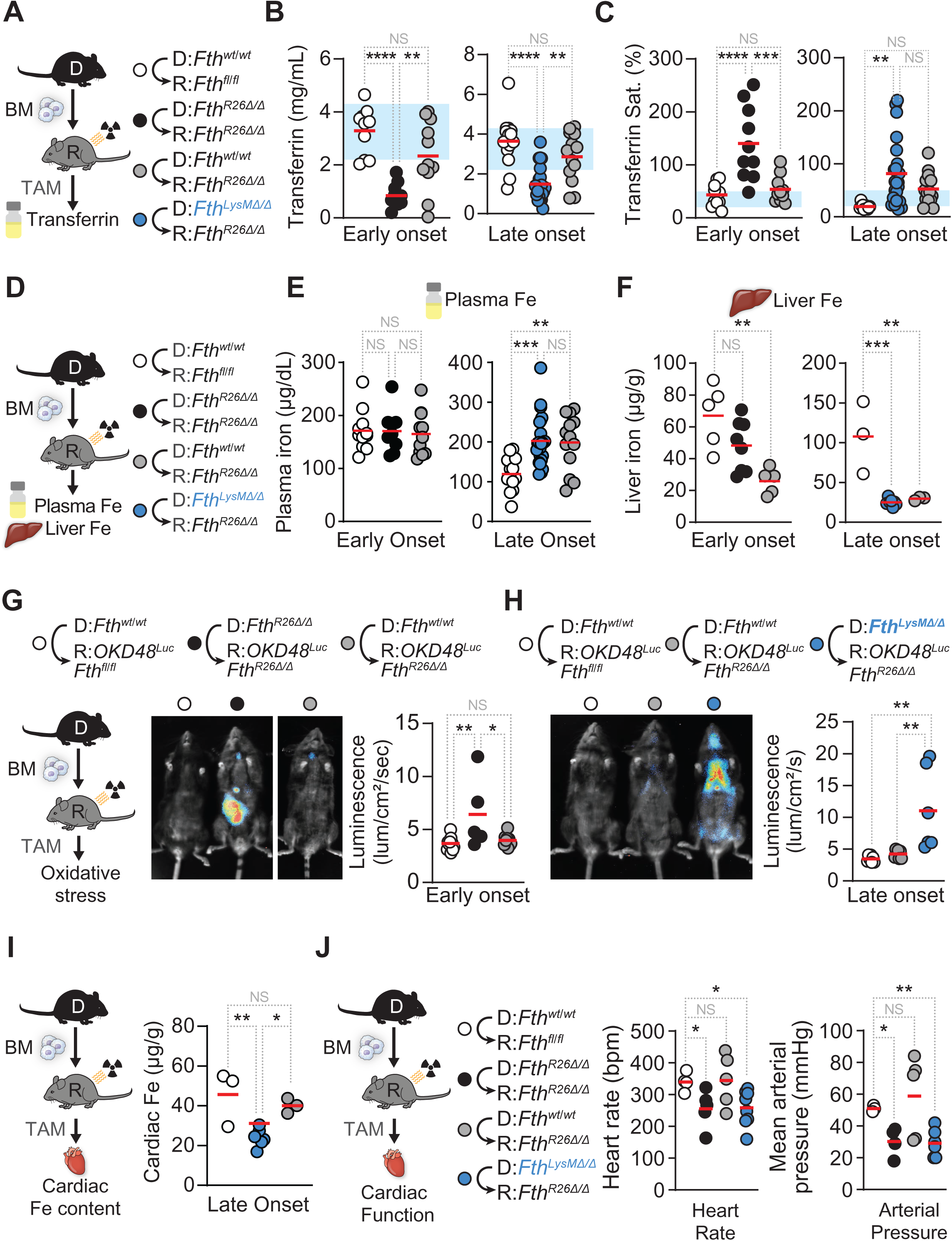
F*t*h-competent myeloid cells restore Fe homeostasis, prevent oxidative stress and restore cardiac function in chimeric *Fth*-deleted mice. (**A**) Schematic representation of chimeric mice and TAM-induced *Fth* deletion (day 0). (**B**) Plasma transferrin concentration and (**C**) transferrin saturation in *Fth^fl/fl^*⇨*Fth^fl/fl^* (n=10-14), *Fth^R26^*^Δ*/*Δ^⇨*Fth^R26^*^Δ*/*Δ^ (n=10-11), *Fth^fl/fl^*⇨*Fth^R26^*^Δ*/*Δ^ (n=11-13) and ⇨*Fth* (n=23) chimeric mice on days 7-15 (early onset), or 19-35 (late onset) following TAM administration. Data in (B, C) pooled from 6 independent experiments with similar trends. (**D**) Schematic representation of chimeric mice and TAM-induced *Fth* deletion. Iron levels in (**E**) plasma measured in *Fth^fl/fl^*⇨*Fth^fl/fl^*(n=11-13), *Fth^R26^*^Δ*/*Δ^⇨*Fth^R26^*^Δ*/*Δ^ (n=9), *Fth^fl/fl^*⇨*Fth^R26^*^Δ*/*Δ^ (n=11-13) and *Fth^LysM^*^Δ*/*Δ^⇨*Fth^R26^*^Δ*/*Δ^ (n=23) chimeric mice on days 7-15 (early onset), or 19-35 (late onset) following TAM administration and (**F**) liver measured in *Fth^fl/fl^*⇨*Fth^fl/fl^* (n=3-5), *Fth^R26^*^Δ*/*Δ^⇨*Fth^R26^*^Δ*/*Δ^ (n=8), *Fth^fl/fl^*⇨*Fth^R26^*^Δ*/*Δ^ (n=3-5) and *Fth^LysM^*^Δ*/*Δ^⇨*Fth^R26^*^Δ*/*Δ^ (n=7) chimeric mice on day 7 (early onset), or 22 (late onset) following TAM administration. Data in (E, F) pooled from 3 independent experiments with similar trends. (**G**) Schematic representation of TAM-induced *Fth* deletion in chimeric mice (day 0). Representative luminescence images depicting oxidative stress, as monitored by OKD48*^Luc^* reporter luciferase luminescence, in *Fth^fl/fl^*⇨*OKD48^Luc^Fth^fl/fl^*(n=7-11), *Fth^R26^*^Δ*/*Δ^⇨*OKD48^Luc^Fth^R26^*^Δ*/*Δ^ (n=5), *Fth^fl/fl^*⇨*OKD48^Luc^Fth^R26^*^Δ*/*Δ^ (n=7-9) and *Fth^LysM^*^Δ*/*Δ^⇨*OKD48^Luc^Fth^R26^*^Δ*/*Δ^ (n=6) chimeric mice on days 7 (**G**; early onset) or 19 (**H**; late onset) following TAM administration. Data in (G, H) presented as individual values (circles) and mean (red bars), pooled from 6 independent experiments with similar trends. (**I**) Schematic representation of chimeric mice and TAM-induced *Fth* deletion (day 0) and cardiac iron content (right) in *Fth^fl/fl^*⇨*Fth^fl/fl^*(n=3-5), *Fth^fl/fl^*⇨*Fth^R26^*^Δ*/*Δ^ (n=3-5) and *Fth^LysM^*^Δ*/*Δ^⇨*Fth^R26^*^Δ*/*Δ^ (n=6-7). (**J**) Schematic representation of chimeric mice and TAM-induced *Fth* deletion (day 0) and cardiac function parameters (heart rate; mean arterial pressure) in *Fth^fl/fl^*⇨*Fth^fl/fl^* (n=4), *Fth^R26^*^Δ*/*Δ^⇨*Fth^R26^*^Δ*/*Δ^ (n=5), *Fth^fl/fl^*⇨*Fth^R26^*^Δ*/*Δ^ (n=5) and *Fth^LysM^*^Δ*/*Δ^⇨*Fth^R26^*^Δ*/*Δ^ (n=8) chimeric mice between days 10-51 following TAM administration. Data in (J) presented as individual values (circles) and mean (red bars), pooled from 2 independent experiments with similar trends. Data in (B, C, E, F-J) represented as individual values (circles) and mean (red bars), One-Way ANOVA with Tukey’s range test for multiple comparison correction was used for comparison between multiple groups. NS: non-significant, * P<0.05, ** P<0.01, *** P<0.001, **** P<0.0001.

Systemic *Fth* deletion was associated with a sharp decrease of hepatic Fe content, compared to control *Fth*-competent chimeras (*Fig. 2F*). This was not restored however, in *Fth^fl/fl^*⇨*Fth^R26^*^Δ*/*Δ^ nor *Fth^LysM^*^Δ*/*Δ^⇨*Fth^R26^*^Δ*/*Δ^ chimeras (*Fig. 2F*). Moreover, *Fth*-deleted *Fth^R26^*^Δ*/*Δ^⇨*Fth^R26^*^Δ*/*Δ^ and *Fth^LysM^*^Δ*/*Δ^⇨*Fth^R26^*^Δ*/*Δ^ chimeras presented electron-dense crystalline inclusions in both the liver (*Fig. S3A*) and WAT (*Fig. S3B*), consistent with the formation of hemosiderin Fe deposits, likely as a result of loss of iron storage capacity upon *Fth* deletion (*Fig. 2F*). These siderosome-like structures, formed by a single membrane-bound lysosomal body, were not observed in either the liver (*Fig. S3A*) or the WAT (*Fig. S3B*) of *Fth^fl/fl^*⇨*Fth^R26^*^Δ*/*Δ^ nor control *Fth^fl/fl^*⇨*Fth^fl/fl^* chimeras. These observations suggest that *Fth*-competent monocyte-derived macrophages cannot restore the loss of hepatic Fe storage capacity.

### *Fth*-competent myeloid cells restore redox homeostasis in chimeric *Fth*-deleted mice

Using *OKD48^Luc^* reporter mice, expressing a luciferase reporter ubiquitously to monitor oxidative stress *in vivo* (Oikawa *et al*, 2012), we confirmed that global *Fth* deletion led to systemic oxidative stress in *Fth^R26^*^Δ*/*Δ^⇨*OKD48^Luc^Fth^R26^*^Δ*/*Δ^ chimeras (*i.e., OKD48^Luc^Fth^R26fl/fl^* mice reconstituted with BM from *Fth^R26fl/fl^* mice)(*Fig. 2G*), consistent with described in adult *Fth^R26^*^Δ*/*Δ^ mice (Blankenhaus *et al*., 2019). *Fth*-competent hematopoietic cells restored systemic redox balance in *Fth^fl/fl^*⇨*OKD48^Luc^Fth^R26^*^Δ*/*Δ^ chimeras (*Fig. 2G,H*). This was no longer the case however, when *Fth* was deleted in myeloid cells (*Fig. 2H*), as assessed at a later point (*i.e.*, late onset) in *Fth^LysM^*^Δ*/*Δ^⇨*OKD48^Luc^Fth^R26^*^Δ*/*Δ^ chimeras (*Fig. 2H*). This suggests that *Fth*-competent monocyte-derived macrophages are essential to prevent the development of tissue oxidative stress imposed by systemic *Fth* deletion.

### *Fth*-competent myeloid cells support tissue function in chimeric *Fth*-deleted mice

Global *Fth* deletion was associated with the development of multiorgan damage, as illustrated serologically by the accumulation of alanine aminotransferase (ALT; liver damage) (*Fig. S4A,B*), aspartate aminotransferase (AST; liver damage) (*Fig. S4A,C*), lactate dehydrogenase (LDH) (*Fig. S4A,D*), creatinine phosphokinase (CPK; muscle damage) (*Fig. S4A,E*) and urea (kidney damage) (*Fig. S4A,F*) in the plasma of *Fth^R26^*^Δ*/*Δ^⇨*Fth^R26^*^Δ*/*Δ^ chimeras *vs.* control *Fth*-competent *Fth^fl/fl^*⇨*Fth^fl/fl^*chimeras. Multiorgan damage was confirmed histologically, as illustrated in the liver (*i.e.,* hepatocellular vacuolar degeneration) and in the kidneys (*i.e.,* acute tubular cell necrosis).

*Fth*-competent hematopoietic cells prevented the multiorgan damage in *Fth^fl/fl^*⇨*Fth^R26^*^Δ^ chimeras (*Fig. S4A-G*). This was no longer observed however, when *Fth* was deleted in myeloid cells from *Fth^LysM^*^Δ*/*Δ^⇨*Fth^R26^*^Δ*/*Δ^ chimeras, which showed hepatocellular hypertrophic lesions with granular acidophilic cytoplasm, hinting at possible liver mitochondrial dysfunction (*Fig. S4A-H*). This suggests that *Fth*-competent monocyte-derived macrophages are essential to prevent multiorgan damage imposed by systemic *Fth* deletion.

### *Fth*-deleted mice succumb to life-threatening cardiac dysfunction

FTH is highly expressed in the heart (Munro & Linder, 1978) and *Fth* deletion in cardiac myocytes leads to cardiac dysfunction (Fang *et al*, 2020), suggesting that *Fth^R26^*^Δ*/*Δ^ mice develop cardiac dysfunction. However, global *Fth* deletion was not associated with the development of acute cardiac damage, as illustrated by troponin I accumulation in plasma from *Fth^R26^*^Δ*/*Δ^⇨*Fth^R26^*^Δ*/*Δ^ *vs.* control *Fth^fl/fl^*⇨*Fth^fl/fl^*chimeras (*Fig. S4I*) and confirmed histologically (*Fig. S4J*). Nevertheless, global *Fth* deletion in *Fth^R26^*^Δ*/*Δ^ mice led to a profound reduction of heart rate, end-systolic pressure (ESP) and preload recruitable stroke work (PRSW), compared to control *R26^Cre^*or *Fth^fl/fl^* mice (*Fig. S5A-C*). A number of other cardiac function parameters, including cardiac mechanical power (MaxPwr) with ensuing reduction of mean arterial pressure, cardiac output, stroke work, pressure-volume area and Pmax were also severely impaired in *Fth^R26^*^Δ*/*Δ^ *vs.* control *R26^Cre^* or *Fth^fl/fl^* mice (*Fig. S5A-C*). This suggests that FTH is essential to sustain heart function and blood circulation.

### *Fth*-competent myeloid cells support cardiac function in chimeric *Fth*-deleted mice

Systemic *Fth* deletion did not impact on cardiac Fe content in *Fth^R26^*^Δ*/*Δ^⇨*Fth^R26^*^Δ*/*Δ^ *vs.* control *Fth^fl/fl^*⇨*Fth^fl/fl^*chimeras (*Fig. S5D*). In contrast however, myeloid *Fth* deletion led to a marked decrease in cardiac Fe content in *Fth^LysM^*^Δ*/*Δ^⇨*Fth^R26^*^Δ*/*Δ^ chimeras, at a later time point, compared to control *Fth^fl/fl^*⇨*Fth^fl/fl^*chimeras (*Fig. 2I*). This was prevented by *Fth* competent hematopoietic cells in *Fth^fl/fl^*⇨*Fth^R26^*^Δ*/*Δ^ chimeras (*Fig. 2I*), suggesting that over time, *Fth*-competent myeloid cells become essential to control the cardiac Fe content of chimeric mice in which *Fth* is deleted globally.

We then asked whether macrophages prevent the development of life-threatening cardiac dysfunction imposed by systemic *Fth* deletion. Similar to observed upon *Fth* deletion in adult *Fth^R26^*^Δ*/*Δ^ mice (*Fig. S5A-C*), global *Fth* deletion in *Fth^R26^*^Δ*/*Δ ⇨^*Fth^R26Δ/Δ^* chimeric mice led to the development of life-threatening cardiac dysfunction, revealed by a profound reduction of heart rate and mean arterial pressure (*Fig. 2J*) as well as ESP, PRSW, and MaxPwr (*Fig. S5E*), compared to control *Fth^fl/fl^*⇨*Fth^fl/fl^*chimeras (*Fig. 2J, Fig. S5E*). *Fth*-competent hematopoietic cells prevented the development of life-threatening cardiac dysfunction in *Fth*-deleted *Fth^fl/fl^*⇨*Fth^R26^*^Δ^ chimeras (*Fig. 2J, Fig. S5E*). This suggests that *Fth*-competent hematopoietic cells carry the capacity to prevent the development of life-threatening cardiac dysfunction imposed by systemic *Fth* deletion.

*Fth*-deletion in the myeloid compartment of *Fth^LysM^*^Δ*/*Δ^⇨*Fth^R26^*^Δ*/*Δ^ chimeras was associated with the development of life-threatening cardiac dysfunction (*Fig. 2J, Fig. S5E*). This suggests that *Fth*-competent myeloid cells, including monocyte-derived macrophages, are essential to prevent the development of life-threatening cardiac dysfunction imposed by systemic *Fth* deletion. This is consistent with monocyte-derived macrophages promoting cardiac function under steady state conditions (Hulsmans *et al*, 2017) or in response to ischemic damage (Swirski *et al*., 2009).

### *Fth*-competent myeloid cells support energy metabolism in chimeric *Fth*-deleted mice

Cardiac function is a highly energy demanding process, entertaining the hypothesis that *Fth*-competent monocyte-derived macrophages support cardiac function, indirectly, via a mechanism that sustains organismal energy expenditure. In support of this hypothesis, global *Fth* deletion led to a collapse of energy expenditure (EE) in *Fth^R26^*^Δ*/*Δ^⇨*Fth^R26^*^Δ*/*Δ^ chimeras (*Fig. 3A,B*), similar to observed *Fth^R26^*^Δ*/*Δ^ mice (Blankenhaus *et al*., 2019). *Fth*-competent hematopoietic cells rescued EE in *Fth^fl/fl^*⇨*Fth^R26^*^Δ*/*Δ^ chimeras (*Fig. 3A,B*) while *Fth*-deletion in myeloid cells compromised the capacity of hematopoietic cells to rescue EE in *Fth^LysM^*^Δ*/*Δ^⇨*Fth^R26^*^Δ*/*Δ^ chimeras (*Fig. 3A,C*).

**Figure 3:**
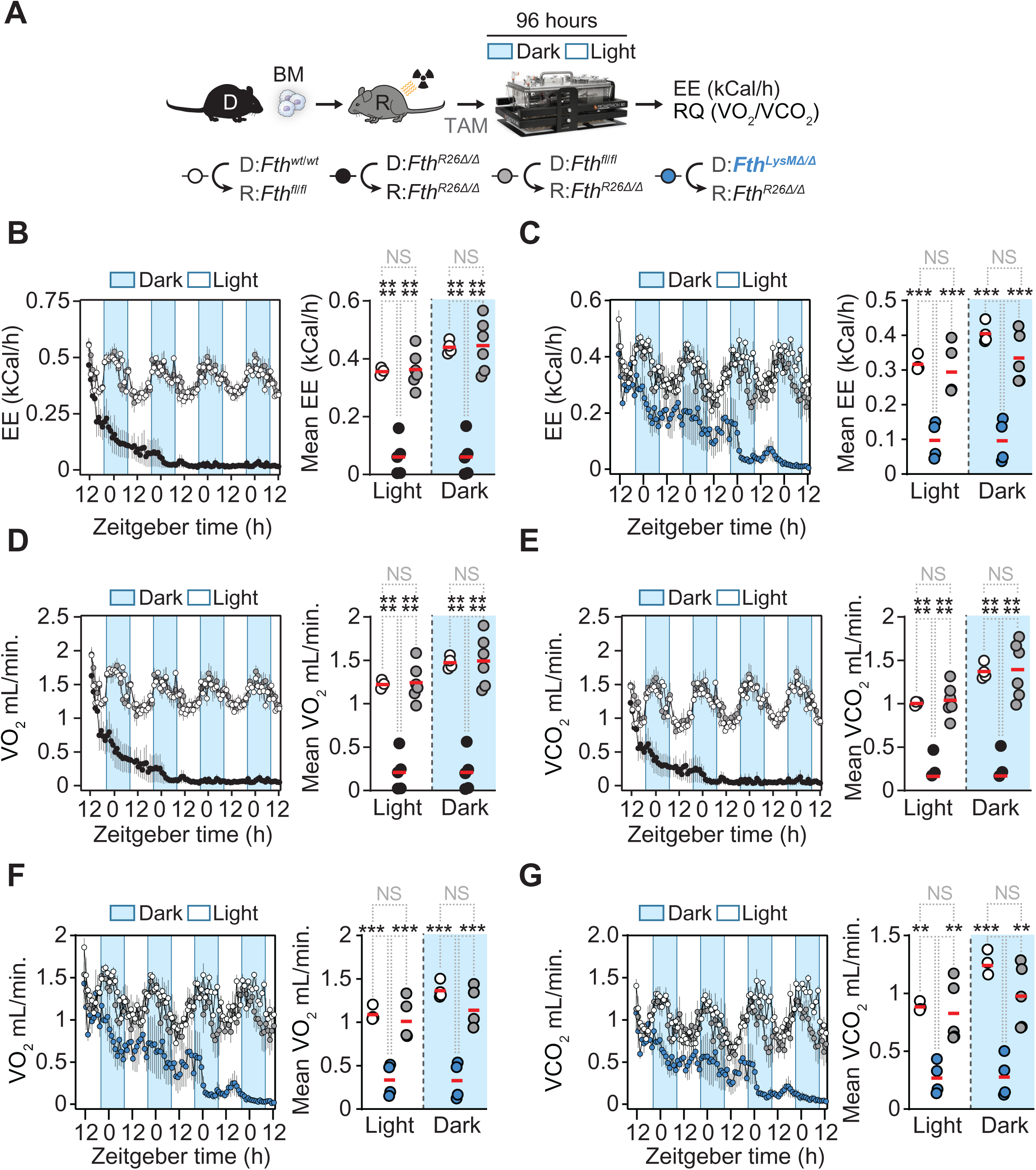
*Fth*-competent myeloid cells support energy metabolism in chimeric *Fth*-deleted mice. (**A**) Schematic representation of TAM-induced *Fth* deletion in chimeric mice (day 0). (**B**, **C**) Time course and mean of energy expenditure (EE) during day/night time in *Fth^fl/fl^*⇨*Fth^fl/fl^*(n=4), *Fth^R26^*^Δ*/*Δ^⇨*Fth^R26^*^Δ*/*Δ^ (n=5), *Fth^fl/fl^*⇨*Fth^R26^*^Δ*/*Δ^ (n=5-6) and *Fth^LysM^*^Δ*/*Δ^⇨*Fth^R26^*^Δ*/*Δ^ (n=4) chimeric mice, assessed from day 7 (**B**; early onset), or day 20 (**C**; late onset) post-TAM administration. (**D**, **E**) Time course and mean of (**D**) O_2_ consumption rate (VO_2_) and (**E**) CO_2_ production rate (VCO_2_) during day/nighttime in *Fth^fl/fl^*⇨*Fth^fl/fl^* (n=4), *Fth^R26^*^Δ*/*Δ^ *Fth^R26^*^Δ*/*Δ^ (n=5) and *Fth^fl/fl^*⇨*Fth^R26^*^Δ*/*Δ^ (n=5-6) chimeric mice, assessed from day 7 (early onset). Time course and mean of (**F**) O_2_ consumption rate (VO_2_) and (**G**) CO_2_ production rate (VCO_2_) during day/nighttime in *Fth^fl/fl^*⇨*Fth^fl/fl^* (n=4), *Fth^LysM^*^Δ*/*Δ^⇨*Fth^R26^*^Δ*/*Δ^ (n=4) and *Fth^fl/fl^*⇨*Fth^R26^*^Δ*/*Δ^ (n=5-6) chimeric mice, assessed from day 20 (late onset). Data in (B-G) is displayed as mean ± SD (time course), or as individual values (circles) and mean (red bars) (dot plots). Data in (B-G) is pooled from 2 independent experiments with similar trends. One-Way ANOVA with Tukey’s range test for multiple comparison correction was used for comparison between multiple groups. NS: non-significant, ** P<0.01, *** P<0.001, **** P<0.0001.

Systemic *Fth* deletion reduced consumed O_2_ (VO_2_) (*Fig. 3D*) and exhaled CO_2_ (VCO_2_) to the same extent (*Fig. 3E*) and therefore did not reflect on respiratory quotient (RQ), (*Fig. S3A-C*). *Fth*-competent hematopoietic cells restored VO_2_ and VCO_2_ in *Fth^fl/fl^*⇨*Fth^R26^*^Δ*/*Δ^ chimeras (*Fig. 3D,E*) while *Fth*-deleted myeloid cells failed to do so in Fth^LysMΔ/Δ^⇨Fth^R26Δ/Δ^ Global *Fth* deletion in chimeric mice was associated with the suppression of locomotor activity (*Fig. S6D*) and food intake (*Fig. S6E*). *Fth*-competent hematopoietic cells restored locomotor activity (*Fig. S6D*) and food intake (*Fig. S6E*) in *Fth^fl/fl^*⇨*Fth^R26^*^Δ*/*Δ^ chimeras (*Fig. S6D,E*). This was no longer observed upon *Fth*-deletion in myeloid cells (*Fig. S6F;G*).

These observations suggest that *Fth*-competent monocyte-derived macrophages are essential to restore energy metabolism under *Fth* deletion in parenchyma cells. This salutary effect is associated a regain of locomotor activity and food intake.

### *Fth*-competent myeloid cells support BAT thermogenesis in chimeric *Fth*-deleted mice

Mice allocate up one third of their EE to support core body temperature, under standard husbandry conditions (∼22°C) (Ganeshan & Chawla, 2017; Ganeshan *et al*, 2019; Reitman, 2018). This is achieved, to a large extent, via brown adipose tissue (BAT) thermogenesis (Cannon & Nedergaard, 2004; Lowell *et al*, 1993), entertaining the hypothesis that *Fth*-competent monocyte-derived macrophages sustain BAT thermogenesis in chimeric mice. In support of this hypothesis, global *Fth* deletion compromised BAT thermogenesis (*Fig. 4A,B*) and caused BAT wasting (*Fig. S7A,B*) in *Fth^R26^*^Δ*/*Δ^⇨*Fth^R26^*^Δ*/*Δ^ chimeras. *Fth*-competent hematopoietic cells restored BAT thermogenesis in *Fth^fl/fl^*⇨*Fth^R26^*^Δ*/*Δ^ chimeras (*Fig. 4A,B*): This was no longer the case when *Fth* was deleted in myeloid cells (*Fig. 4C,D*), with a tendency for BAT wasting and reduction in average lipid droplet at a later time point (*Fig. S7C,D*), albeit not significant. This suggests that *Fth*-competent myeloid cells, including monocyte-derived macrophages, prevent the collapse of BAT thermogenesis and thermoregulation imposed by systemic *Fth* deletion.

**Figure 4:**
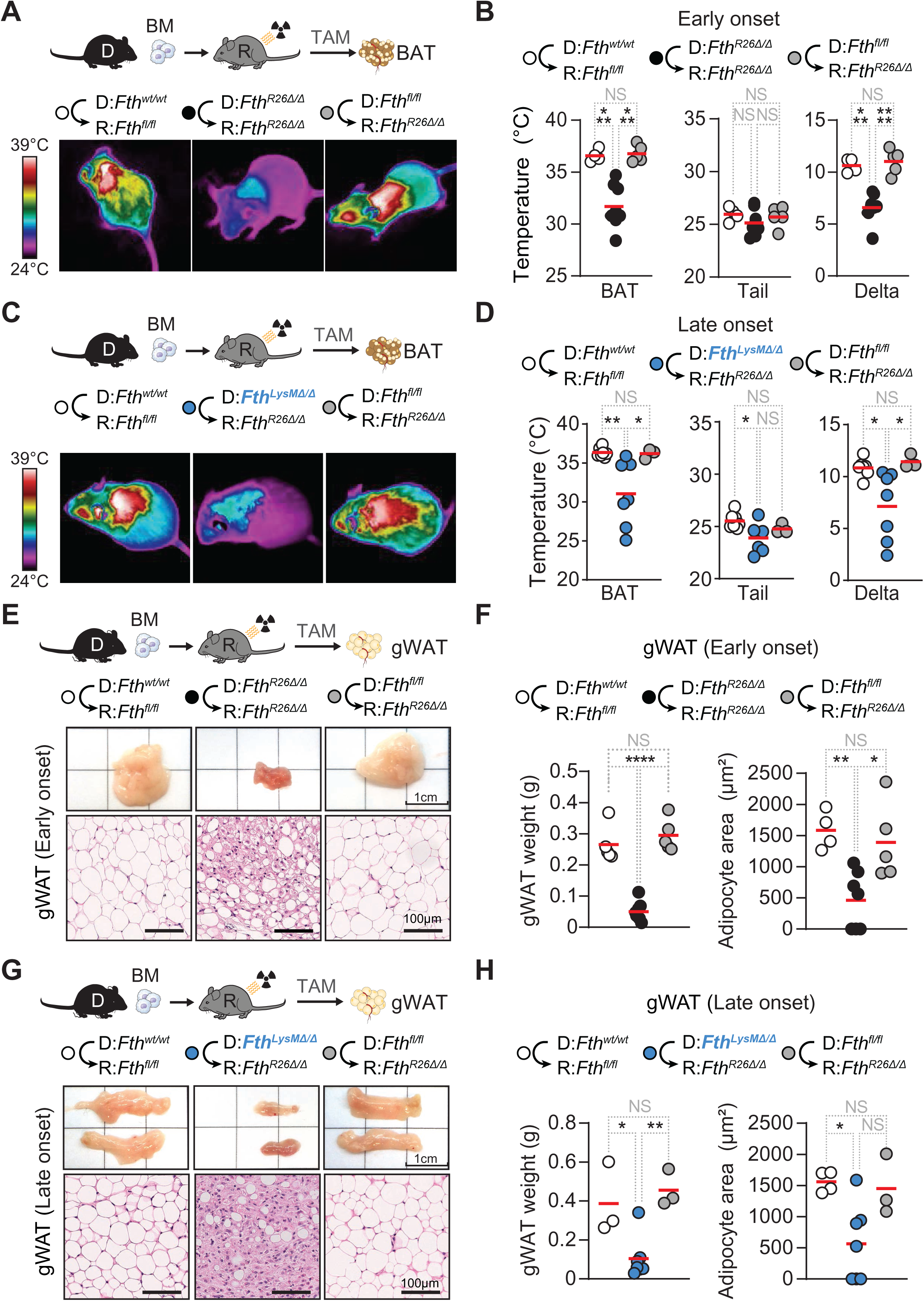
*Fth*-competent myeloid cells support adipose tissue function in chimeric *Fth*-deleted mice. (**A**) Schematic representation of chimeric mice and TAM-induced *Fth* deletion (day 0) with corresponding representative infrared thermal images (FLIR) of *Fth^fl/fl^*⇨*Fth^fl/fl^*, *Fth^R26^*^Δ*/*Δ^⇨*Fth^R26^*^Δ*/*Δ^ and *Fth^fl/fl^*⇨*Fth^R26^*^Δ*/*Δ^ chimeric mice, on day 7 (early onset) post-TAM administration. (**B**) BAT (left) and tail (center) temperatures extracted from thermal images, and temperature delta (right; core body temperature – tail temperature) of *Fth^fl/fl^*⇨*Fth^fl/fl^*(n=4), *Fth^R26^*^Δ*/*Δ^⇨*Fth^R26^*^Δ*/*Δ^ (n=8) and *Fth^fl/fl^*⇨*Fth^R26^*^Δ*/*Δ^ (n=5) chimeric mice, collected on day 7 (early onset) post-TAM administration. (**C**) Schematic representation of chimeric mice and TAM-induced *Fth* deletion (day 0) with corresponding representative (FLIR) of *Fth^fl/fl^*⇨*Fth^fl/fl^*, *Fth^LysM^*^Δ*/*Δ^⇨*Fth^R26^*^Δ*/*Δ^ and *Fth^fl/fl^*⇨*Fth^R26^*^Δ*/*Δ^ chimeric mice, collected between days 16-39 (late onset) post TAM administration using an infrared thermal imaging camera (FLIR). (**D**) BAT (left) and tail (center) temperatures extracted from thermal images, and temperature delta (right; core body temperature – tail temperature) of *Fth^fl/fl^*⇨*Fth^fl/fl^* (n=7), *Fth^LysM^*^Δ*/*Δ^⇨*Fth^R26^*^Δ*/*Δ^ (n=7) and *Fth^fl/fl^*⇨*Fth^R26^*^Δ*/*Δ^ (n=3) chimeric mice, collected between days 16-39 (late onset) post TAM administration. Data in (B, D) represented as individual values (circles) and mean (red bars). Data in (A-D) is pooled from 3 independent experiments with similar trends. (**E**) Schematic representation of chimeric mice and TAM-induced *Fth* deletion, with corresponding representative macroscopic and histological images of gonodal white adipose tissue pads (gWAT) in *Fth^fl/fl^*⇨*Fth^fl/fl^*, *Fth^R26^*^Δ*/*Δ^⇨*Fth^R26^*^Δ*/*Δ^ and *Fth^fl/fl^*⇨*Fth^R26^*^Δ*/*Δ^ chimeric mice, collected on day 7 (early onset) post TAM administration. (**F**) gWAT weight (left) and mean (red bars) adipocyte area (right) in *Fth^fl/fl^*⇨*Fth^fl/fl^*(n=4-5), *Fth^R26^*^Δ*/*Δ^⇨*Fth^R26^*^Δ*/*Δ^ (n=7-8) and *Fth^fl/fl^*⇨*Fth^R26^*^Δ*/*Δ^ (n=5) chimeric mice, collected on day 7 (early onset) post TAM administration. (**G**) Schematic representation of chimeric mice and TAM-induced *Fth* deletion, with corresponding representative macroscopic and histological images of gonodal white adipose tissue pads (gWAT) in *Fth^fl/fl^*⇨*Fth^fl/fl^*, *Fth^LysM^*^Δ*/*Δ^⇨*Fth^R26^*^Δ*/*Δ^ and *Fth^fl/fl^*⇨*Fth^R26^*^Δ*/*Δ^ chimeric mice, collected between days 16-39 (late onset) post-TAM administration. (**H**) gWAT pad weight (left) and mean (red bars) adipocyte area (right) of *Fth^fl/fl^*⇨*Fth^fl/fl^* (n=3-4), *Fth^LysM^*^Δ*/*Δ^⇨*Fth^R26^*^Δ*/*Δ^ (n=7) and *Fth^fl/fl^*⇨*Fth^R26^*^Δ*/*Δ^ (n=3) chimeric mice, collected between days 16-39 (late onset) post-TAM administration. Data in (F, H) represented as individual values (circles) and mean (red bars). Data in (E-H) is pooled from 3 independent experiments with similar trends. One-Way ANOVA with Tukey’s range test for multiple comparison correction was used for comparison between multiple groups. NS: non-significant, * P<0.05, ** P<0.01, *** P<0.001, **** P<0.0001.

### *Fth*-competent myeloid cells support WAT function in chimeric *Fth*-deleted mice

BAT thermogenesis is fueled via white adipose tissue (WAT) lipolysis (Heine *et al*, 2018), entertaining the hypothesis that *Fth*-competent monocyte-derived macrophages regulate WAT lipolysis. In keeping with this hypothesis, global *Fth* deletion led to visceral (*i.e.,* gonadal) (*Fig. 4E,F*) and subcutaneous (*i.e.,* inguinal) WAT wasting (*Fig. S7E,F*), with reduction of lipid droplet (*i.e.,* adipocyte) size in *Fth^R26^*^Δ*/*Δ^⇨*Fth^R26^*^Δ*/*Δ^ chimeras (*Fig. 4E,F; S7E,F*). While *Fth*-competent hematopoietic cells prevented WAT wasting in *Fth^fl/fl^*⇨*Fth^R26^*^Δ*/*Δ^ chimeras (*Fig. 4E,F; S7E,F*), WAT wasting was prominent when *Fth* was deleted in myeloid cells from *Fth^LysM^*^Δ*/*Δ^⇨*Fth^R26^*^Δ*/*Δ^ chimeras (*Fig. 4G,H; S7G,H*). This suggests that *Fth*-competent myeloid cells that give rise to monocyte-derived macrophages, carry the capacity to normalize WAT function and BAT thermogenesis in chimeric mice in which *Fth* is deleted globally.

### *Fth*-competent myeloid cells partially restore ferritin expression in tissues from chimeric *Fth*-deleted mice

Macrophages secrete and transfer ferritin (Cohen et al., 2010; Meyron-Holtz et al., 2011), suggesting that FTH might be transferred from *Fth*-competent monocyte-derived macrophages to *Fth*-deficient bystander parenchyma cells. In keeping with this hypothesis, *Fth*-competent hematopoietic cells restored 30-40% of the relative level of FTH protein expression in heart, liver, lung and kidneys of *Fth^fl/fl^*⇨*Fth^R26^*^Δ*/*Δ^ chimeras, compared to control *Fth^fl/fl^*⇨*Fth^fl/fl^* chimeras (*Fig. 5A,B*). In contrast, FTH protein expression was barely detectable when *Fth* was deleted in myeloid cells from *Fth^LysM^*^Δ*/*Δ^⇨*Fth^R26^*^Δ*/*Δ^ chimeras (*Fig. 5A,B*). This suggests that under global *Fth* deletion, *Fth*-competent monocyte-derived macrophages are essential to partially restore the level of FTH protein expression in different organs.

**Figure 5:**
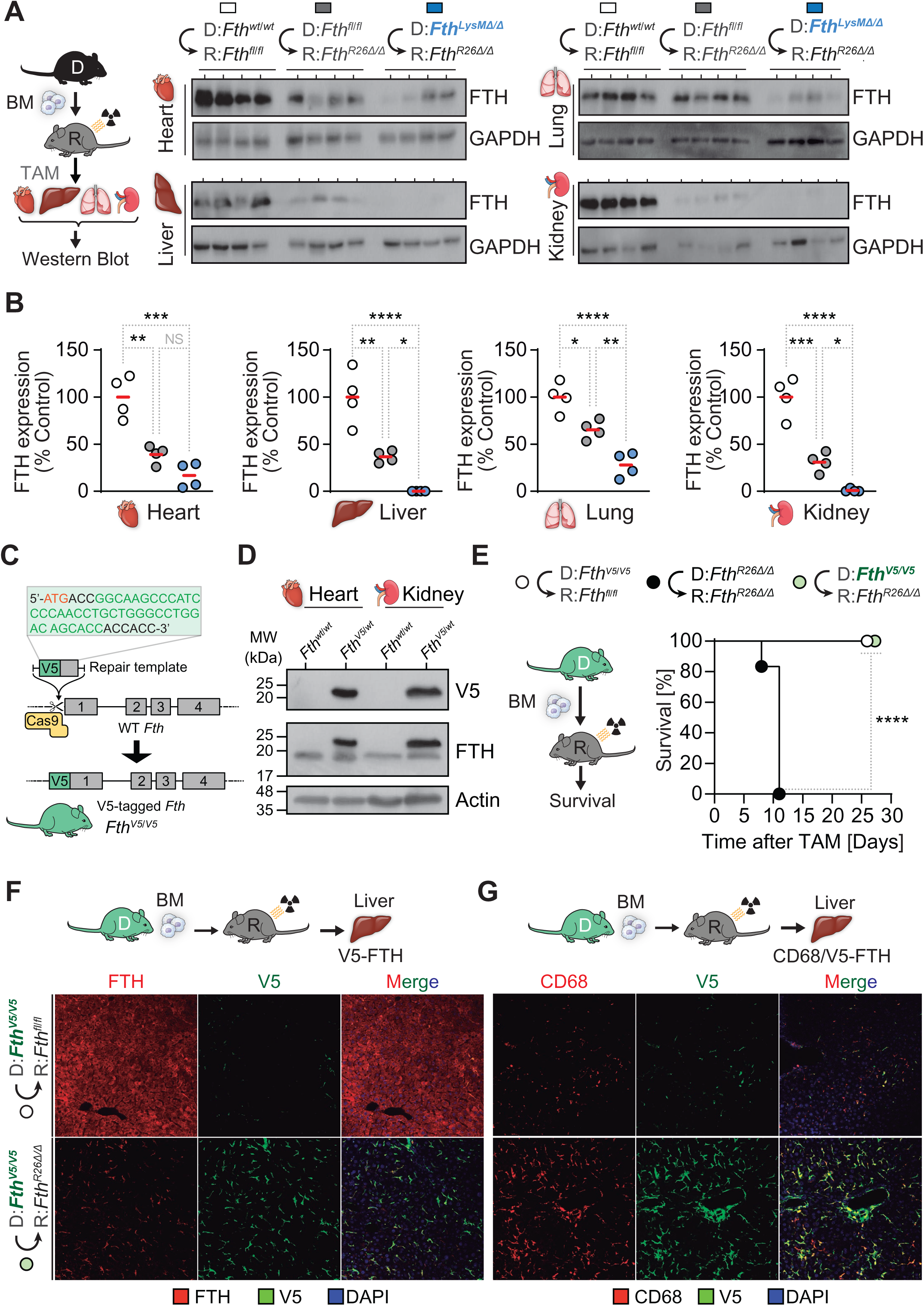
F*t*h-competent myeloid cells partially restore tissue ferritin expression but do not deliver ferritin to parenchyma cells in chimeric *Fth*-deleted mice. (**A**) Schematic representation of TAM-induced *Fth* deletion in chimeric mice (day 0) and analysis of FTH protein expression (Western blot). (**B**) Relative quantification of FTH protein expression in heart, liver, lungs and kidneys from *Fth^fl/fl^*⇨*Fth^fl/fl^* (n=4), *Fth^LysM^*^Δ*/*Δ^⇨*Fth^R26^*^Δ*/*Δ^ (n=4) and *Fth^fl/fl^*⇨*Fth^R26^*^Δ*/*Δ^ (n=4) chimeric mice, between days 7-23 post-TAM administration. Data in (B) represented as the % of FTH expression relative to *Fth^fl/fl^*⇨*Fth^fl/fl^*control chimeras and is pooled from 3 independent experiments. (**C**) Schematic representation of the CRISPR/CaS9 editing strategy for V5-tagged FTH. (**D**) Detection of V5-tagged and endogenous FTH protein expression (Western blot) in control *Fth^wt/wt^* and edited *Fth^V5/wt^* mice. (**E**) Schematic representation of TAM-induced *Fth* deletion (day 0) and survival of *Fth^V5/V5^*⇨*Fth^fl/fl^* (n=13), *Fth^R26^*^Δ*/*Δ^⇨*Fth^R26^*^Δ*/*Δ^ (n=6) and *Fth^V5/V5^*⇨*Fth^R26^*^Δ*/*Δ^ (n=11) chimeric mice. Data in (E) is pooled from 4 independent experiments with similar trends. (**F**) Schematic representation of TAM-induced *Fth* deletion in chimeric mice, and representative images of FTH and V5 immunofluorescence detection in livers from *Fth^V5/V5^*⇨*Fth^fl/fl^*and *Fth^V5/V5^*⇨*Fth^R26^*^Δ*/*Δ^ chimeric mice. (**G**) Schematic representation of TAM-induced *Fth* deletion and representative images of CD68 (macrophages) and V5 immunofluorescence detection in livers from *Fth^V5/V5^*⇨*Fth^fl/fl^* and *Fth^V5/V5^*⇨*Fth^R26^*^Δ*/*Δ^ chimeric mice. One-Way ANOVA with Tukey’s range test for multiple comparison correction was used for comparison between multiple groups. Survival analysis was performed using Log-rank (Mantel-Cox) test. NS: non-significant, * P<0.05, ** P<0.01, *** P<0.001, **** P<0.0001.

### *Fth*-competent myeloid cells do not deliver ferritin to parenchyma cells from chimeric *Fth*-deleted mice

To determine whether or not monocyte-derived macrophages transfer FTH to parenchyma cells we used a CRISPR/CaS9 based approach (Jinek et al., 2012) to tag the N-terminus of FTH with a 14 amino-acid V5 peptide (*Fth^V5/WT^*mouse)(*Fig. 5C*). Expression of V5-tagged FTH protein was validated in whole heart and kidney extracts from hemizygous *Fth^V5/WT^* mice, by western blot (*Fig. 5D*).

Hematopoietic cells expressing V5-tagged FTH retained the capacity to rescue the lethal outcome of systemic *Fth* deletion in *Fth^V5/V5^*⇨*Fth^R26^*^Δ*/*Δ^ chimeras (*i.e., Fth^R26fl/fl^* mice reconstituted with *Fth^V5/V5^* BM) (*Fig. 5E*). However, the expression of the V5-tagged FTH protein was restricted to CD68^+^ macrophages, without detectable transfer to parenchyma cells, as assessed by confocal microscopy in the liver of *Fth^V5/V5^*⇨*Fth^R26^*^Δ*/*Δ^ chimeras (*Fig. 5F,G*). This suggests that *Fth*-competent monocyte-derived macrophages rescue the lethal outcome of systemic *Fth* deletion, irrespectively of ferritin transfer to *Fth*-deleted parenchyma cells.

### *Fth*-competent myeloid cells rescue chimeric *Fth*-deleted mice irrespectively of cellular Fe import/export

We hypothesized that *Fth*-competent macrophages rescue the lethal outcome of global *Fth* deletion via a mechanism that involves cellular Fe import and/or Fe export to or from *Fth*-deleted parenchyma cells, respectively. To test these hypothesis, we reconstituted *Fth^R26fl/fl^*mice with BM cells from mice harboring a myeloid-specific deletion of the main cellular Fe importer transferrin receptor 1 (TfR1; encoded by *Tfrc*) (*Fig. S8A*) or the cellular Fe exporter Ferroportin (FPN1; encoded by *Slc40a1*) (*Fig. S8B*) (Wu *et al*., 2023). *Slc40a1*-deleted myeloid cells (*Fig. S8A*) or *Tfrc-*deleted myeloid cells (*Fig. S8B*) retained the capacity to rescue the lethal outcome of systemic *Fth* deletion in *Slc40a1^LysM^*^Δ*/*Δ^⇨*Fth^R26^*^Δ*/*Δ^ or *Tfrc^LysM^*^Δ*/*Δ^⇨*Fth^R26^*^Δ*/*Δ^ chimeras, respectively (*Fig. S8*). This shows that monocyte-derived macrophages rescue the lethal outcome of global *Fth* deletion, via a mechanism that does not rely on Fe transit between *Fth*-competent macrophages and *Fth*-deleted parenchyma cells.

### *Fth*-competent myeloid cells support the mitochondria of *Fth*-deleted parenchyma cells from chimeric *Fth*-deleted mice

Whole-body *Fth* deletion causes a severe disruption of mitochondria structure and function in parenchyma cells (Blankenhaus *et al*., 2019). This suggests that *Fth*-competent macrophages rescue the lethal outcome of global *Fth* deletion, via a mechanism that supports the mitochondria of *Fth*-deleted parenchyma cells. In contrast to *Fth*-deletion in adult *Fth^R26^*^Δ*/*Δ^ mice (Blankenhaus *et al*., 2019), *Fth* deletion *Fth^R26^*^Δ*/*Δ^⇨*Fth^R26^*^Δ*/*Δ^ or *Fth^LysM^*^Δ*/*Δ^⇨*Fth^R26^*^Δ*/*Δ^ chimeras was not associated with a reduction of the number of mitochondria *per* cell, compared to control *Fth^fl/fl^*⇨*Fth^fl/fl^*chimeras, as assessed in the heart, WAT and liver (*Fig.6A-C*). However, the mitochondria morphology of parenchyma cells from *Fth^R26^*^Δ*/*Δ^⇨*Fth^R26^*^Δ*/*Δ^ chimeras was clearly disrupted, as reveled by swelling and irregular shape of mitochondrial *cristae* and reduced electron density of the mitochondrial matrix (*Fig.6D*). Normal mitochondrial morphological features were restored in *Fth^fl/fl^*⇨*Fth^R26^*^Δ*/*Δ^ but not in *Fth^LysM^*^Δ*/*Δ^⇨*Fth^R26^*^Δ*/*Δ^ chimeras (*Fig. 6D*), suggesting that *Fth*-competent macrophages are essential to support the mitochondrial structure under systemic *Fth* deletion.

**Figure 6:**
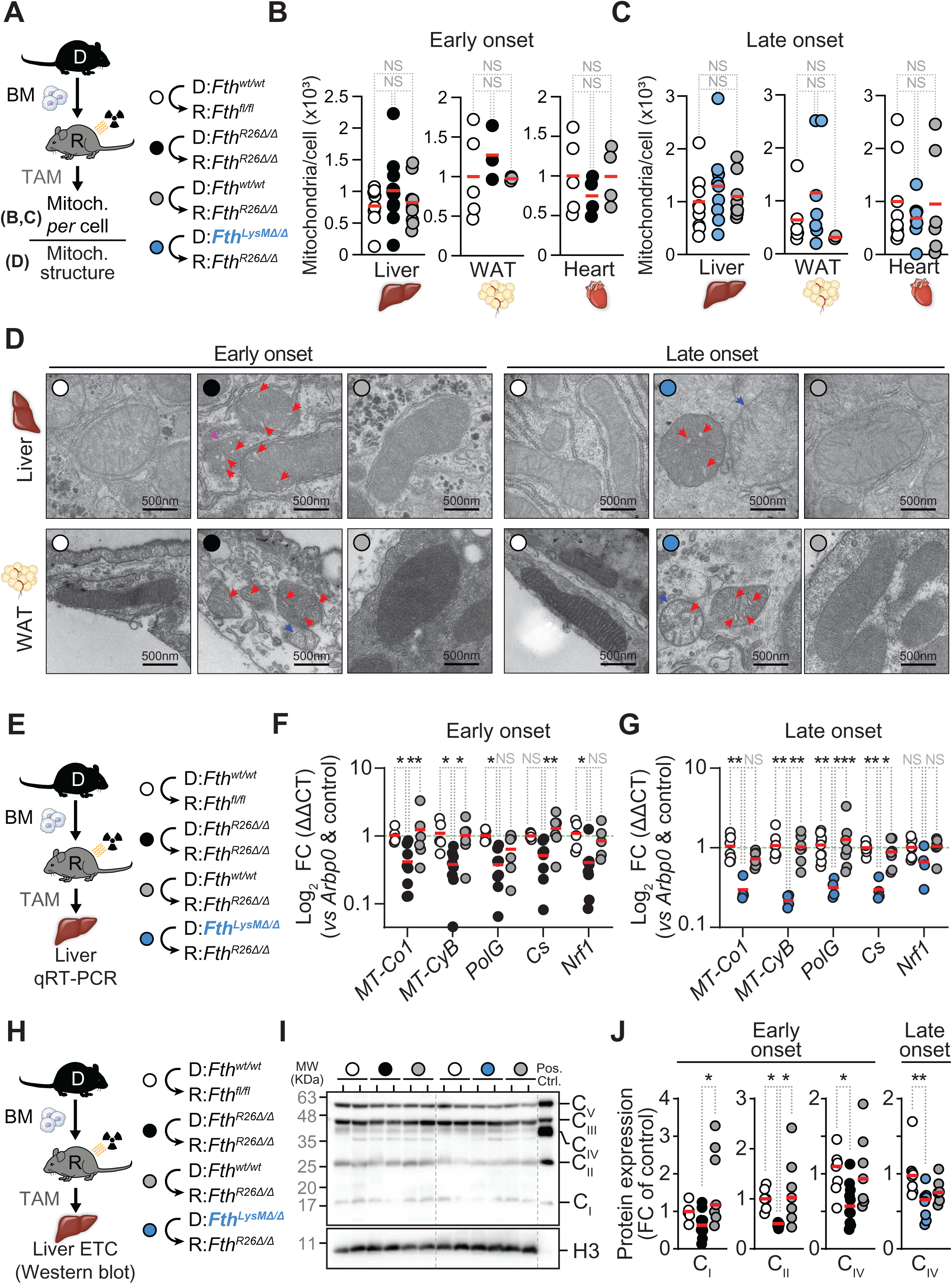
*Fth*-competent monocyte-derived macrophages support the mitochondria of parenchyma cells from *Fth*-deleted chimeric mice. (**A**) Schematic representation of chimeric mice, TAM administration (day 0) and mitochondrial analysis. (**B**, **C**) Number of mitochondria *per* cell in liver, WAT and heart of (**B**) *Fth^fl/fl^*⇨*Fth^fl/fl^*(n=5-8), *Fth^R26^*^Δ*/*Δ^⇨*Fth^R26^*^Δ*/*Δ^ (n=3-12) and *Fth^fl/fl^*⇨*Fth^R26^*^Δ*/*Δ^ (n=3-9) chimeric mice, between days 7-9 (early onset) or (**C**) *Fth^fl/fl^*⇨*Fth^fl/fl^* (n=6-10), *Fth^LysM^*^Δ*/*Δ^⇨*Fth^R26^*^Δ*/*Δ^ (n=2-8) and *Fth^fl/fl^*⇨*Fth^R26^*^Δ*/*Δ^ (n=7-8) chimeric mice, on day 22-39 (late onset). Data in (B, C) represented as individual values (circles) and mean (red bars), assessed by quantitative PCR according to the ratio of nuclear (i.e. hexokinase 2; *Hk2*) and mitochondrial (*i.e.,* NADH-ubiquinone oxidoreductase chain 1; *mt-Nd1*) DNA. (**D**) Representative transmission electron microscopy images of mitochondria structure in gWAT adipocytes and liver hepatocytes from *Fth^fl/fl^*⇨*Fth^fl/fl^*, *Fth^R26^*^Δ*/*Δ^⇨*Fth^R26^*^Δ*/*Δ^ and *Fth^fl/fl^*⇨*Fth^R26^*^Δ*/*Δ^ chimeric mice, on day 8 (early onset), or from *Fth^fl/fl^*⇨*Fth^fl/fl^*, *Fth^LysM^*^Δ*/*Δ^⇨*Fth^R26^*^Δ*/*Δ^ and *Fth^fl/fl^*⇨*Fth^R26^*^Δ*/*Δ^ chimeric mice, on day 30 (late onset). Red arrows indicate swollen cristae, purple arrows indicate disrupted mitochondrial membranes, blue arrows indicate loss of mitochondrial membrane potential, as suggested by decreased electron density. (**E**) Schematic representation of chimeric mice, TAM administration (day 0) and qRT-PCR assessment. (**F**, **G**) Log_2_ fold change (FC) in expression of mitochondrial genes or genes involved in regulation of mitochondria in livers from (**F**) *Fth^fl/fl^*⇨*Fth^fl/fl^*(n=5), *Fth^R26^*^Δ*/*Δ^⇨*Fth^R26^*^Δ*/*Δ^ (n=9) and *Fth^fl/fl^*⇨*Fth^R26^*^Δ*/*Δ^ (n=6) chimeric mice, between days 7-8 (early onset), or from (**G**) *Fth^fl/fl^*⇨*Fth^fl/fl^*(n=9), *Fth^LysM^*^Δ*/*Δ^⇨*Fth^R26^*^Δ*/*Δ^ (n=4) and *Fth^fl/fl^*⇨*Fth^R26^*^Δ*/*Δ^ (n=7) chimeric mice, between days 19 and 22 (late onset). Data in (F, G) is pooled from 3 independent experiments with similar trend and represented as individual values (circles) and mean (red bars), normalized to housekeeping gene expression (acidic ribosomal phosphoprotein P0, *Arbp0*) and to gene expression values of control *Fth^fl/fl^*⇨*Fth^fl/fl^* chimeric mice. (**H**) Schematic representation of chimeric mice, TAM administration (day 0) and western blot assessment. (**I**) Representative Western blot and (**J**) quantification of ETC subunits for complexes I-V in the livers of *Fth^fl/fl^*⇨*Fth^fl/fl^*(n=7), *Fth^R26^*^Δ*/*Δ^⇨*Fth^R26^*^Δ*/*Δ^ (n=10) and *Fth^fl/fl^*⇨*Fth^R26^*^Δ*/*Δ^ (n=7) chimeric mice, between days 7-8 (early onset), or from *Fth^fl/fl^*⇨*Fth^fl/fl^*(n=8), *Fth^LysM^*^Δ*/*Δ^⇨*Fth^R26^*^Δ*/*Δ^ (n=8) and *Fth^fl/fl^*⇨*Fth^R26^*^Δ*/*Δ^ (n=6) chimeric mice, between days 19 and 22 (late onset). Data in (J) is pooled from 4 independent experiments with similar trends. One-Way ANOVA with Tukey’s range test for multiple comparison correction was used for comparison between multiple groups. Two-Way ANOVA with Tukey’s range test for multiple comparison correction was used for comparison between multiple groups within multiple genes. NS: non-significant, * P<0.05, ** P<0.01, *** P<0.001.

### *Fth*-competent myeloid cells restore mitochondria gene expression in chimeric *Fth*-deleted mice

Global *Fth* deletion in *Fth^R26^*^Δ*/*Δ^⇨*Fth^R26^*^Δ*/*Δ^ chimeras was associated with a reduction of the relative level of expression of hepatic mitochondrial mRNA, encoded by mitochondrial (*i.e., MT-Co1*: ETC complex IV Cytochrome C oxidase I, *MT-Cyb:* ETC complex III cytochrome B; *MT-PolG*: Mitochondrial DNA polymerase) or nuclear (*Cs*: Citrate synthase; *Nrf1*: Nuclear respiratory factor 1) genes, as compared to control *Fth^fl/fl^*⇨*Fth^fl/fl^*chimeras (*Fig. 6E,F*). *Fth*-competent hematopoietic cells restored the expression of these hepatic mitochondrial mRNAs in *Fth^fl/fl^*⇨*Fth^R26^*^Δ^ chimeras (*Fig. 6E,F*). This was no longer the case when *Fth* was deleted in myeloid cells from ⇨*Fth* chimeras (*Fig. E,G*).

Systemic *Fth* deletion was also associated with reduction of the relative level of a subset of mitochondrial ETC proteins, as assessed in the liver of *Fth^R26^*^Δ*/*Δ^⇨*Fth^R26^*^Δ*/*Δ^ *vs.* control *Fth^fl/fl^*⇨*Fth^fl/fl^* chimeras (*Fig. 6H-J; S8E*). *Fth*-competent hematopoietic cells restored the expression of these mitochondrial ETC proteins (*Fig. 6H-J; S8E*), while deletion of *Fth* in myeloid cells compromised this rescuing effect (*Fig. 6H-J; S8E*). These observations suggest that *Fth*-competent monocyte-derived macrophages engage in an intercellular crosstalk that supports the mitochondria of *Fth*-deficient parenchyma cells.

### Mitochondrial biogenesis is a hallmark of rescuing macrophages

To explore the mechanism via which *Fth*-competent monocyte-derived macrophages restore the mitochondria of *Fth*-deficient chimeric mice, we generated *tdT^LysM^* mice expressing tdT under the control of the LysM promoter. These were crossed with *Fth^fl/fl^* mice, to generate *tdT^LysM^Fth^LysM^*^Δ*/*Δ^ mice, deleting *Fth* specifically in tdT^+^ myeloid cells.

There was a marked reduction in the number of viable monocytes/macrophages in the liver WAT and heart from chimeric mice reconstituted with tdT^+^ *Fth*-deleted myeloid cells (*i.e., Fth^R26fl/fl^* mice reconstituted with *tdT^LysM^Fth^LysM^*^Δ*/*Δ^ BM; *tdT^LysM^Fth^LysM^*^Δ*/*Δ^⇨*Fth^R26^*^Δ*/*Δ^ chimeras), compared to control chimeras (*i.e., Fth^fl/fl^* mice reconstituted with *tdT^LysM^* BM; *tdT^LysM^*⇨*Fth^fl/fl^*)(*Fig.S9A,C,D*). This was not observed in chimeric mice reconstituted with *Fth*-competent hematopoietic cells (*i.e., Fth^R26fl/fl^* mice reconstituted with *tdT^LysM^*BM; *tdT^LysM^*⇨*Fth^R26^*^Δ*/*Δ^).

To characterize the transcriptional response of rescuing monocyte-derived macrophages, tdT^+^ (CD45^+^CD11b^+^Ly6G^_^CD19^_^TCRβ^_^) monocytes/macrophages were FACS-sorted from the liver and WAT of chimeric mice (*Fig.S9A,C*). Analyzes of RNA sequencing (RNAseq) revealed a marked upregulation of a large swath of mitochondrial genes in *Fth*-competent *vs. Fth*-deleted macrophages from *tdT^LysM^*⇨*Fth^R26^*^Δ*/*Δ^ *vs. tdT^LysM^Fth^LysM^*^Δ*/*Δ^⇨*Fth^R26^*^Δ*/*Δ^ chimeras, respectively (*Fig.7A,B*; *Fig.S9E-F*). Gene ontology enrichment showed a wide range of terms directly related to mitochondria (*Fig.7C*), suggesting that *Fth* is strictly required to support this transcriptional response.

**Figure 7:**
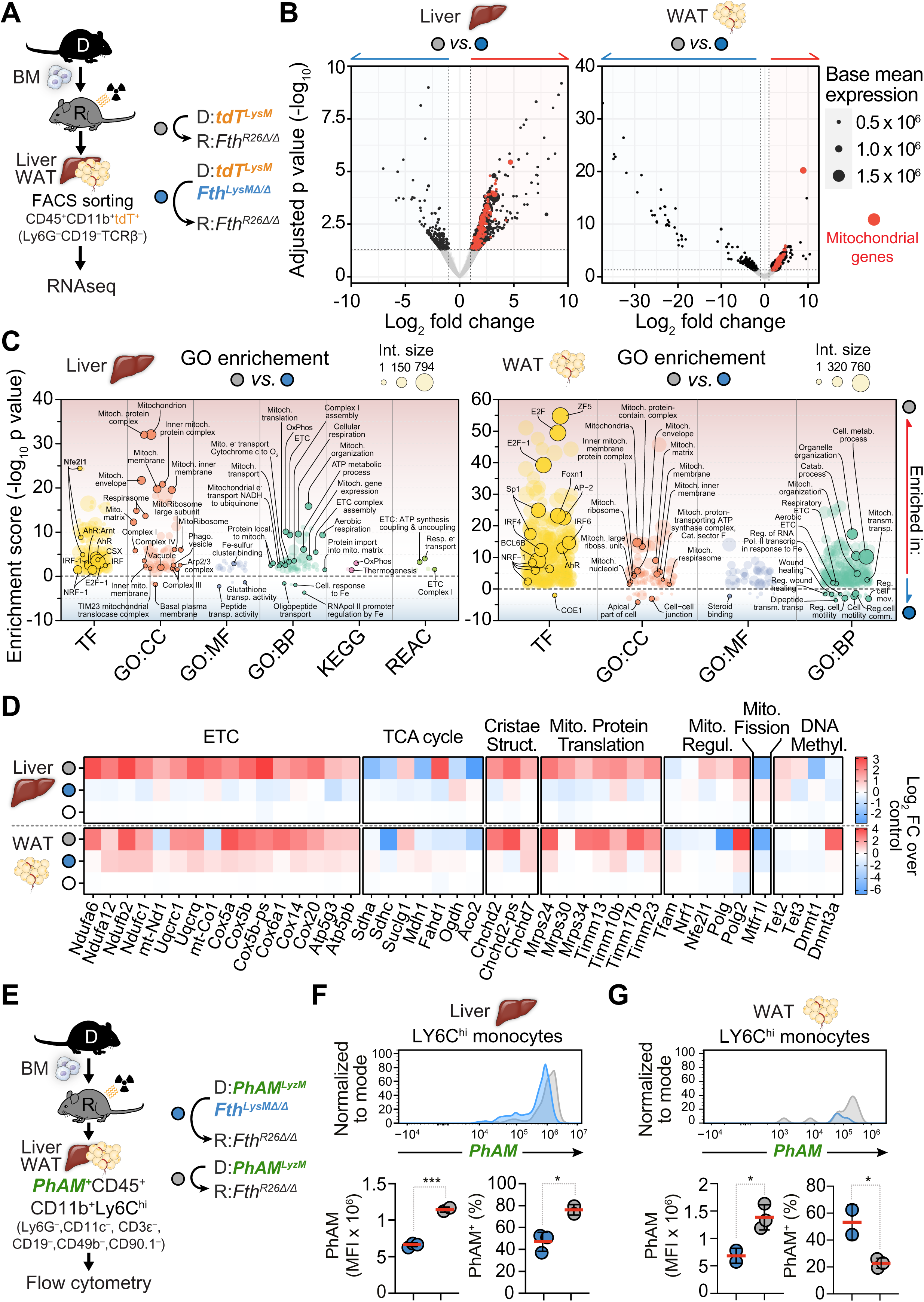
*Fth*-competent monocyte-derived macrophages deploy a mitochondrial gene transcriptional program. (**A**) Schematic representation of chimeric mice, TAM administration and fluorescence-activated cell sorting (FACS) of *LysM^+^* monocyte/macrophages (CD45^+^,CD11b^+^,Ly6G^−^,CD19^−^,TCRβ^−^) in liver and WAT. (**B**) Volcano plots of differentially regulated genes between *LysM^+^*monocyte/macrophages sorted from liver (left) or WAT (right) of *tdT^LysM^*⇨*Fth^R26^*^Δ*/*Δ^ (n=3), *tdT^LysM^Fth^LysM^*^Δ*/*Δ^⇨*Fth^R26^*^Δ*/*Δ^ (n=5) chimeric mice, on day 19 post-TAM administration. Red dots depict mitochondrial genes that are significantly differentially regulated. (**C**) Gene ontology analysis depicting ontologies that are significantly enriched comparing *LysM^+^*monocyte/macrophages sorted from liver (left) and WAT (right) from *tdT^LysM^*⇨ *Fth^R26^*^Δ*/*Δ^ (n=3) and *tdT^LysM^Fth^LysM^*^Δ*/*Δ^⇨*Fth^R26^*^Δ*/*Δ^ (n=5) chimeric mice on day 19 post-TAM administration. Ontologies significantly enriched in *LysM^+^*monocyte/macrophages from *tdT^LysM^*⇨*Fth^R26^*^Δ*/*Δ^ are depicted as: enrichment score -log_10_ p value > 1,301. Ontologies significantly enriched in *LysM^+^* monocyte/macrophages from *tdT^LysM^Fth^LysM^*^Δ*/*Δ^⇨*Fth^R26^*^Δ*/*Δ^ are depicted as: enrichment score -log_10_ p value < -1,301. Ontology classes: TF = transcription factors; GO:CC = gene ontology : cellular component; GO:MF = GO : molecular function; GO:BP = GO : biological process; KEGG = Kyoto Encyclopedia of Genes and Genomes pathway; REAC = reactome. (**D**) Heatmap of mean Log_2_ FC (over control; *tdT^LysM^*⇨ *Fth^fl/fl^*) in gene expression of genes involved in mitochondrial function and regulation, in *LysM^+^*monocyte/macrophages sorted from the liver (top) and WAT (bottom) of *tdT^LysM^*⇨*Fth^fl/fl^*(n=3), *tdT^LysM^*⇨*Fth^R26^*^Δ*/*Δ^ (n=3) and *tdT^LysM^Fth^LysM^*^Δ*/*Δ^⇨*Fth^R26^*^Δ*/*Δ^ (n=5) chimeric mice on day 19 post-TAM administration. (**E**) Schematic representation of chimeric mice, TAM administration and flow cytometry analysis of PhAM reporter-expressing mitochondria in *LysM^+^* cells. Representative histograms of PhAM fluorescence intensity, mean fluorescence intensity (MFI) of PhAM, and percentage of PhAM^+^ cells in LY6C^high^ monocytes from the liver (**F**) and WAT (**G**) of *PhAM^LysM^*⇨*Fth^fl/fl^*(n=2-3) and *PhAM^LysM^Fth^LysM^*^Δ*/*Δ^⇨*Fth^R26^*^Δ*/*Δ^ (n=2-3) chimeric mice, collected on day 100 post-TAM administration. Data in (F, G) dot plots represented as individual values (circles) and mean (red bars). Student’s t-test was used for comparison between two groups. NS: non-significant, * P<0.05, *** P<0.001.

Analysis of protein-protein interactions for induced genes from enriched mitochondria related ontologies (*Fig.7C*), revealed tight interaction networks involving structural mitochondrial, mitoribosome and respirasome/ETC (oxidative phosphorylation) proteins, in both liver (*Fig.S10A*) and WAT (*Fig.S10B*) macrophages. In addition, the intersection of upregulated genes revealed a network subset shared in both organs, suggesting that these genes are part of a core effector response (*Fig.S10C*). Specifically, the transcriptional response of *Fth*-competent macrophages included the induction of genes supporting the ETC, such as *Uqcrq* (complex III subunit), as well as genes involved in mitochondria biogenesis (e.g. *Nrf1* and *PolG2*), maintenance of mitochondrial cristae structure (e.g. *ChChd2*) and mitochondrial protein translation (*Fig.7D*). Moreover, we also observed a downregulation of genes in the mitochondria tricarboxylic acid (TCA) cycle (*Fig.7D*).

We noted that mitochondrial fission regulator 1 like (*Mtfr1l*), showed decreased expression in in *Fth*-competent *vs. Fth*-deleted macrophages (*Fig.7D*). This suggests that suppression of mitochondrial regulation through fission contributes to the rescuing capacity of *Fth*-competent macrophages (*Fig.7D*).

RNAseq analysis of cardiac *Fth*-competent *vs. Fth*-deleted macrophages from chimeric mice, revealed a distinct transcriptional response that did not include the regulation of mitochondrial genes (*Fig.S11A-B*). This suggests that *Fth*-competent monocyte-derived macrophages activate transcriptional programs that are tissue-specific and likely prevent multi-organ dysfunction imposed by global *Fth* deletion.

### Mitochondrial biogenesis is a hallmark of rescuing monocyte-derived macrophages

To monitor the mitochondria of monocyte-derived macrophages, we introduced an additional *Rosa26-mito-Dendra2-Flox-stop-Flox* allele in *LysM^Cre^*, driving the expression of a Dendra2 fluorescent protein fused to mitochondria targeting sequence of the cytochrome c oxidase subunit VIII (mito-Dendra2/PhAM) (Pham *et al*, 2012) mice. *PhAM^LysM^* mice were crossed with *Fth^fl/fl^* mice to generate *PhAM^LysM^Fth^fl/fl^*mice, deleting *Fth* specifically in PhAM^+^ myeloid cells.

The relative level of PhAM expression was higher in *Fth*-competent *vs. Fth*-deleted monocyte-derived macrophages, from the liver (*Fig.7E,F*) and WAT (*Fig.7E,G*) of *PhAM^LysM^*⇨*Fth^R26^*^Δ*/*Δ^ *vs. PhAM^LysM^Fth^R26^*^Δ*/*Δ^⇨*Fth^R26^*^Δ*/*Δ^ chimeras, respectively. This is consistent with the transcriptional program supporting mitochondrial biogenesis increasing the number of mitochondria in *Fth*-competent *vs. Fth*-deleted monocyte-derived macrophages.

Of note, the percentage of PhAM+ monocyte-derived macrophages was lower in WAT from *PhAM^LysM^*⇨*Fth^R26^*^Δ*/*Δ^ *vs. PhAM^LysM^Fth^R26^*^Δ*/*Δ^⇨*Fth^R26^*^Δ*/*Δ^ chimeras (*Fig.7G*). This suggests that there is an alteration on the relative percentage of one or several other CD45+ populations in *PhAM^LysM^Fth^R26^*^Δ*/*Δ^⇨*Fth^R26^*^Δ*/*Δ^ or *PhAM^LysM^*⇨*Fth^R26^*^Δ*/*Δ^ chimeras.

### *Fth*-competent myeloid cells rescue chimeric *Fth*-deleted mice irrespectively of mitochondrial transfer to parenchyma cells

Monocyte-derived macrophages can transfer functional mitochondria to parenchyma cells (Borcherding & Brestoff, 2023; Nakai *et al*, 2024). This suggested that *Fth*-competent macrophages prevent the lethal outcome of global *Fth* deletion via intercellular mitochondrial transfer to *Fth*-deleted parenchyma cells. To test this hypothesis, we compared the expression of mito-Dendra2 (PhAM) in parenchyma cells from chimeric mice to that of *PhAM^R26^* mice expressing mito-Dendra2 constitutively in parenchyma cells (*Fig. 8A*). Relative expression of mito-Dendra2 was evaluated according to a PhAM index, resulting from the ratio of mean mito-Dendra2 fluorescent intensity over the percentage of mito-Dendra2^+^ parenchyma cells (*Fig. 8A*).

**Figure 8:**
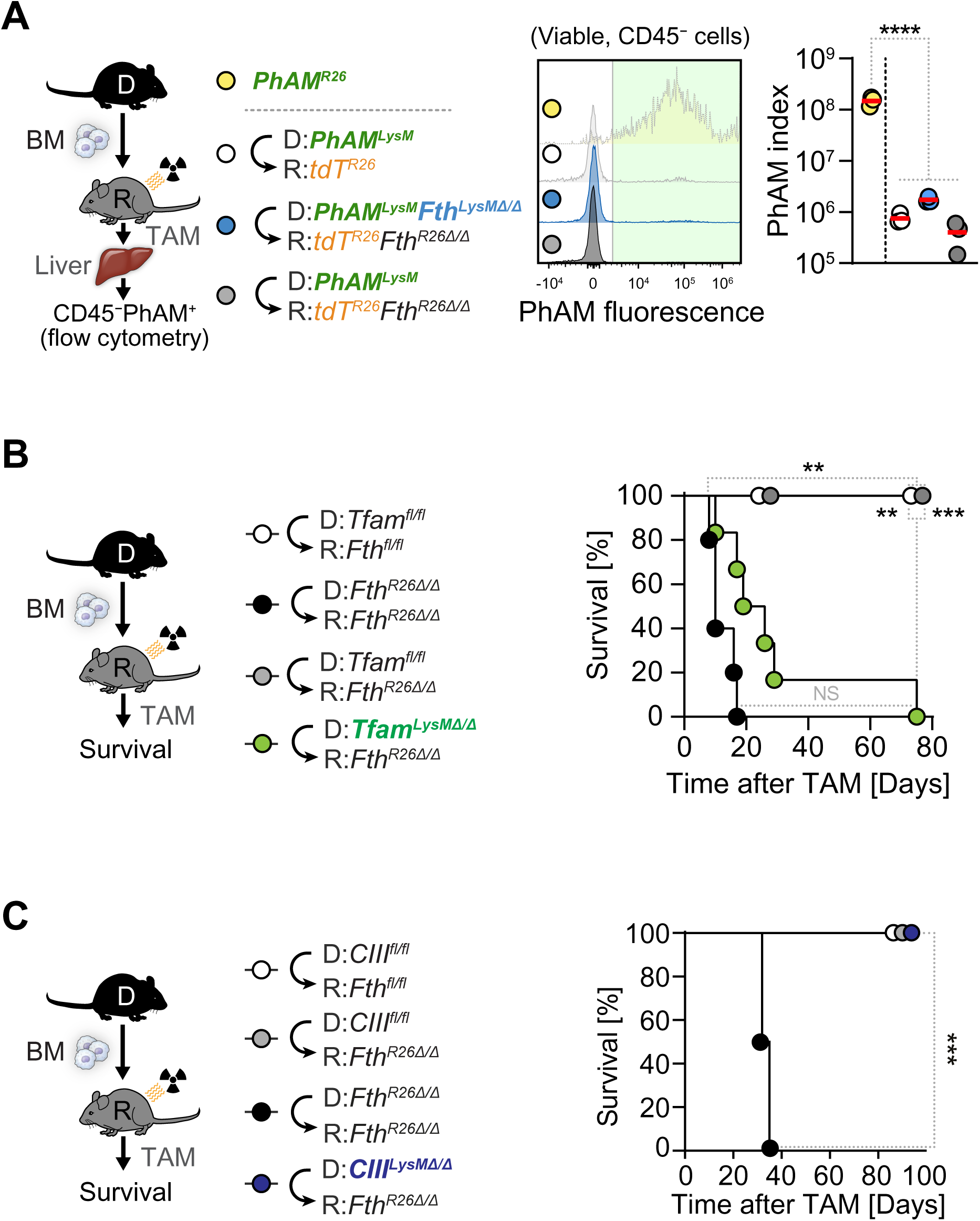
Mitochondrial biogenesis supports the rescuing capacity of *Fth*-competent monocyte-derived macrophages in chimeric *Fth*-deleted mice. (**A**) Schematic representation of chimeric mice, TAM administration (day 0), and flow cytometry analysis of livers from control *PhAM^R26^* mice (n=3), and *PhAM^LysM^*⇨*tdT^R26^*(n=3), *PhAM^LysM^Fth^LysM^*⇨*tdT^R26^Fth^R26^*^Δ*/*Δ^ (n=3), *PhAM^LysM^*⇨*tdT^R26^Fth^R26^*^Δ*/*Δ^ (n=3) chimeras with representative PhAM fluorescence histograms and PhAM expression index of CD45^−^ parenchymal cells. PhAM index is calculated as the percentage of CD45^−^PhAM^+^ cells, multiplied by CD45^−^PhAM^+^ cells’ MFI. (**B**) Schematic representation of chimeric mice, TAM administration (day 0), and survival of *Tfam^fl/fl^*⇨*Fth^fl/fl^*(n=4), *Fth^R26^*^Δ*/*Δ^⇨*Fth^R26^*^Δ*/*Δ^ (n=5), *Tfam^fl/fl^*⇨*Fth^R26^*^Δ*/*Δ^ (n=6) and *Tfam^LysM^*^Δ*/*Δ^⇨*Fth^R26^*^Δ*/*Δ^ (n=6) chimeric mice on day 0. Data in (B) is pooled from 2 independent experiments with similar trends. (**C**) Schematic representation of chimeric mice, TAM administration (day 0), and survival of *CIII^fl/fl^*⇨*Fth^fl/fl^*(n=6), *CIII^fl/fl^*⇨*Fth^R26^*^Δ*/*Δ^ (n=6), *Fth^R26^*^Δ*/*Δ^⇨*Fth^R26^*^Δ*/*Δ^ (n=2), and *CIII^LysM^*^Δ*/*Δ^⇨*Fth^R26^*^Δ*/*Δ^ (n=5) chimeric mice on day 0. Survival analysis was performed using Log-rank (Mantel-Cox) test. NS: non-significant, ** P<0.01, *** P<0.001, **** P<0.0001.

The PhAM index of parenchyma cells in the liver of different chimeric mice was detectable but residual compared to control *PhAM^R26^* (*Fig. 8A*). While consistent with macrophages transferring PhAM^+^ mitochondria to parenchyma cells in chimeric mice (Nakai *et al*., 2024), these observations suggest that *Fth*-competent prevent the lethal outcome of global *Fth* deletion irrespectively of intercellular mitochondrial transfer.

### Myeloid cells rescue chimeric *Fth*-deleted mice via a mechanism that relies on a transcriptional program supporting mitochondrial biogenesis

To test whether mitochondrial biogenesis supports the rescuing capacity of *Fth*-competent macrophages we deleted the master regulator of mitochondrial DNA (mtDNA) transcription and replication TFAM (Larsson *et al*, 1998), specifically in myeloid cells from *Tfam^LysM^*^Δ*/*Δ^ mice (Wculek *et al*, 2023). *Tfam*-deleted myeloid cells failed to prevent the lethal outcome of *Fth*-deletion in *Tfam^LysM^*^Δ*/*Δ^⇨*Fth^R26^*^Δ*/*Δ^ chimeras (*i.e., Fth^R26fl/fl^* mice reconstituted with *Tfam^LysM^*^Δ*/*Δ^ BM), as compared to chimeras reconstituted with *Tfam*-competent myeloid cells (*i.e., Fth^R26fl/fl^* mice reconstituted with *Tfam^fl/fl^*BM; *Tfam^fl/fl^*⇨*Fth^R26^*^Δ*/*Δ^) (*Fig. 8B*). This suggests the regulation of mtDNA replication and/or transcription by TFAM is essential to endow macrophages with the capacity to rescue the lethal outcome of global *Fth* deletion.

TFAM controls the transcription of several components of the mitochondrial ETC (*i.e.,* complex I, III, IV, and V) supporting oxidative phosphorylation (OXPHOS). To disentangle OXPHOS from other mitochondrial functions supported by TFAM (Wculek *et al*., 2023), we deleted *Uqcrq*, the gene encoding the Ubiquinol-cytochrome c reductase, complex III subunit VII, specifically in myeloid cells from *Uqcrq^LysM^*^Δ*/*Δ^ mice (Wculek *et al*., 2023; Weinberg *et al*, 2019). *Uqcrq*-deleted myeloid cells retained the capacity to prevent the lethal outcome of global *Fth*-deletion in *Uqcrq^LysM^*^Δ*/*Δ^⇨*Fth^R26^*^Δ*/*Δ^ chimeras (*i.e., Fth^R26fl/fl^* mice reconstituted with *Uqcrq^LysM^*^Δ*/*Δ^ BM), similar to *Uqcrq*-competent myeloid cells in *Uqcrq^fl/fl^* ⇨*Fth^R26^*^Δ*/*Δ^ chimeras (*i.e., Fth^R26fl/fl^* mice reconstituted with *Uqcrq^fl/fl^* BM)(*Fig. 8C*). This suggests OXPHOS is not essential to support the rescuing capacity of monocyte-derived macrophage.

## DISCUSSION

Our findings suggest that monocyte-derived macrophages become essential to support tissue function and organismal homeostasis in response to dysregulation of Fe homeostasis in parenchyma cells. This is consistent with tissues representing an emergent property of the interactions between “primary” and “supportive” cells (Adler *et al*., 2023; Meizlish *et al*., 2021; Zhou *et al*., 2018b), whereby macrophages are the universal supportive cells that facilitate the tissue-specific functions of parenchyma cells (Adler *et al*., 2023; Meizlish *et al*., 2021; Zhou *et al*., 2018b).

The interaction of macrophages with parenchyma cells is thought to give rise to integrated cellular modules, such as illustrated in the livers by the interaction of Kupfer cells with stellate cells, hepatocytes and endothelial cells (Bonnardel *et al*, 2019). Additional interactions with peripheral neurons provide these integrated cellular modules with the capacity to surveil systemic variations of vital parameters and provide tailored responses (Godinho-Silva *et al*, 2019; Veiga-Fernandes & Mucida, 2016). These integrated cellular modules are also apparent in the adipose tissue, where macrophages establish stable physical interactions with adipocytes, vascular endothelial cells (Moura Silva *et al*, 2021; Silva *et al*, 2019) and peripheral neurons (Pirzgalska *et al*, 2017) as well as with mesenchymal cells (Ko *et al*, 2020). These are essential to regulate WAT lipolysis (Ko *et al*., 2020; Pirzgalska *et al*., 2017) and BAT thermoregulation (Wolf *et al*, 2017; Zeng *et al*, 2015). Our finding that macrophages rescue the lethal outcome of global *Fth* deletion via a mechanism that controls WAT lipolysis and BAT thermoregulation (*Fig. 4*) is consistent with this organizing principle (Adler *et al*., 2023; Meizlish *et al*., 2021; Zhou *et al*., 2018b).

Our findings should contribute to explain how regulation of Fe metabolism modulates WAT function (Blankenhaus *et al*., 2019; Romero *et al*, 2022; Wang *et al*, 2024; Yook *et al*, 2021), BAT thermogenesis (Blankenhaus *et al*., 2019; Wang *et al*., 2024; Yook *et al*., 2021) and energy balance (Blankenhaus *et al*., 2019; Lu *et al*, 2024; Wang *et al*., 2024). Moreover, they are consistent with diet Fe-deficiency compromising thermoregulation in rodents (Dillmann *et al*, 1979) and humans (Beard *et al*, 1990; Brigham & Beard, 1996; Lukaski *et al*, 1990) as well as with diet Fe overload leading to a negative energy balance in rodents (Romero *et al*., 2022).

That circulating rather than tissue-resident macrophages take control of energy homeostasis in response to global *Fth* deletion is demonstrated using parabiosis (*Fig. 1I*) as well as by the adoptive transfer of BMDMo (*Fig. 1J*). While illustrating an extraordinary capacity of circulating monocytes to partake in the inter-organ crosstalk that regulates systemic Fe and energy metabolism, it is likely that other circulating myeloid-derived, such as myeloid-derived suppressor cells (Veglia *et al*, 2021), might cells contribute to this process.

The major impact of Fe on macrophage function (Soares & Hamza, 2016) suggested that macrophages respond directly to the Fe status of parenchyma cells to support organismal homeostasis. This hypothesis, however, is not supported by the observation that macrophages retain the capacity to rescue the lethal outcome of global *Fth* deletion irrespectively of cellular Fe import by TFR1 (*Fig. S8A,B*) or export, by SLC40a1 (*Fig. S8C,D*). Moreover, macrophages rescue the lethal outcome of global *Fth* deletion irrespectively of ferritin secretion and transfer to parenchyma cells (*Fig. 5C-G*). Overall, this suggests that *Fth*-competent macrophages do not rely on inter-cellular Fe sensing and/or transfer to support *Fth*-deleted parenchyma cells.

As an alternative hypothesis, macrophages sense the consequences of dysregulated cellular Fe metabolism in *Fth*-deleted parenchyma cells, namely, mitochondrial dysfunction (*Fig. 6D-G*) (Blankenhaus *et al*., 2019). In support of this hypothesis, macrophages respond to a number of mitochondrial cues (Murphy & O’Neill, 2024), including to the mtDNA released from mitochondria (Murphy & O’Neill, 2024). In keeping with this hypothesis *Fth*-competent macrophages that rescue the lethal outcome of global *Fth* deletion present a gene expression profile consistent with a type I interferon response (*Fig. 7A-C*), a hallmark of the macrophage response to mtDNA (Al Amir Dache & Thierry, 2023; He *et al*, 2022). Moreover, remodeling of the mitochondria *cristae* structure, such as observed in *Fth*-deleted parenchyma cells (*Fig. 6D*) (Blankenhaus *et al*., 2019), triggers the release of mtDNA and induce a type I interferon response in macrophages (He *et al*., 2022).

*Fth*-competent macrophages that rescue the lethal outcome of global *Fth* deletion presented a singular gene expression profile, associated with maintenance of mitochondrial function (*i.e.,* ETC, TCA) and structure (*i.e., cristae*) as well as mitochondrial biogenesis (*Fig. 7A-G*). This genetic program is consistent with the transcriptional response controlled by TFAM (Larsson *et al*., 1998), a mitochondrial transcriptional regulator that is required to endow macrophages with the capacity yo rescue the lethal outcome of global *Fth* deletion (*Fig. 8B*).

While TFAM plays an essential role in supporting the expression of mitochondrial ETC genes (Wculek *et al*., 2023), this is probably not essential for macrophages to rescue *Fth*-deleted chimeric mice (*Fig. 8B*). This suggests that the induction of a transcriptional program supporting mitochondria biogenesis is the functional hallmark of macrophages that rescue the lethal outcome of global *Fth* deletion.

The genetic program supporting mitochondria biogenesis (*Fig. 7A-D*) was activated only in *Fth*-competent macrophages from *Fth*-deleted chimeric recipient mice (*Fig. S9C-F*). This suggests that FTH expression in macrophages is essential to support this transcriptional program, activated by *Fth* deleted parenchyma cells.

It is possible that the transcriptional program supporting mitochondria biogenesis in macrophages involves the DNA methyl transferase 3A (DNMT3A). In support of this hypothesis, we found that DNMT3A was highly induced in *Fth*-competent macrophages that rescue the lethal outcome of global *Fth* deletion (*Fig. 7D*). This is consistent with FTH supporting DNMT3A expression (Ye *et al*, 2019) and with DNMT3A regulating the expression of TFAM in macrophages (Cobo *et al*, 2022). A non-mutually exclusive hypothesis is that FTH regulates TFAM via a mechanism involving the ten-eleven translocation (TET) dioxygenases, which use Fe as an essential co-factor and the TCA-derived α-ketoglutarate as a substrate to catalyze cytosine demethylation (Lopez-Moyado *et al*, 2024). In support of this hypothesis we found that FTH regulates cytosine demethylation by TET dioxygenases (Wu *et al*, 2024), which induce TFAM expression macrophages (Pan *et al*, 2017).

While consistent with recent findings in a similar experimental model (Nakai *et al*., 2024) our findings suggest that macrophages rescue the lethal outcome of global *Fth* deletion irrespectively of intercellular mitochondrial transfer to parenchyma cells. This is supported by the detectable but residual intercellular mitochondrial transfer from *Fth*-competent macrophages to *Fth*-deleted parenchyma cells in chimeric mice (*Fig. 8A*).

A non-mutually exclusive hypothesis is that the cytoprotective effect of FTH (Berberat *et al*, 2003; Gozzelino *et al*, 2012; Pham *et al*, 2004) is essential for macrophages to phagocytose mitochondria, extruded from *Fth*-deleted parenchyma cells in the chimeric mice. In support of this hypothesis, mitochondria phagocytosis is a major supportive function of macrophages as illustrated in BAT (Brestoff *et al*, 2021; Rosina *et al*, 2022) and the heart (Nicolás-Ávila *et al*, 2020). While consistent with the depletion of *Fth*-deleted monocyte-derived macrophages in *Fth*-deleted chimeric mice (*Fig. 1F; S9D*) this hypothesis remains however to be tested experimentally.

In conclusion, monocyte-derived macrophages take control of organismal Fe, redox and energy metabolism in response to global *Fth* deletion. This extraordinary capacity relies on a mechanism whereby FTH supports a transcriptional program that acts in a cell autonomous manner to promote mitochondrial biogenesis and in a non-cell autonomous manner to regulate the mitochondria of parenchyma cells in different tissues. These findings support the notion that macrophages play a central role in supporting the function of parenchyma cells in different tissues to sustain homeostatic control of multicellular organisms.

## ACKNOWLEDGEMENTS

The authors are indebted to all members of the Inflammation group (GIMM) for insightful technical and intellectual contributions, to Erin M. Tranfield and Ana L. Vinagre at the Electron Microscopy Unit (GIMM), to the staff at the flow cytometry facility, as well as to the staff at the GIMM animal facility.

**MPS** was supported by Gulbenkian, ‘La Caixa’ foundation (HR18-00502) and FCT (PTDC/IMI-IMU/5723/2014; FEDER/29411/2017; PTDC/MED-FSL/4681/2020, <u>10.54499/PTDC/MED-FSL/4681/2020</u>; and 2022.02426.PTDC, <u>10.54499/2022.02426.PTDC</u>), Oeiras-ERC Frontier Research Incentive Awards, SymbNET Research Grants (H2020-WIDESPREAD-2020-5-952537), DFG Cluster of Excellence ‘‘Balance of the Microverse’’ (DFG, EXC 2051; 390713860) and Congento (LISBOA-01-0145-FEDER-022170). **RM** was supported by an EMBO long-term fellowship (ALTF290-2017ARC), Marie Skłodowska-Curie Research Fellowship (MSCA-IF-EF-ST-753236) and Fundação para a Ciência e a Tecnologia (2021.03494.CEECIND; <u>10.54499/2021.03494.CEECIND/CP1674/CT0004</u>). **BB** and **SC** were supported in part by European Community 7th Framework 294709-DAMAGECONTROL ERC-2011-AdG to MPS, FB by Marie Skłodowska-Curie Research Fellowship (REGDAM 707998). **SW** was supported by the German Ministry of Education and Research (BMBF; grant 01 EO 1502) via the Jena Center of Sepsis Control and Care. **MM** was supported by Fundação para a Ciência e a Tecnologia (UI/BD/152257/2021). **QW** was supported by a Marie Skłodowska-Curie Research Fellowship (MSCA-IF2019-892773), the International Postdoctoral Exchange Fellowship Program from the Peoplés Republic of China (20190090), and the National Natural Science Foundation of China (32171166, 82030003).

## AUTHOR CONTRIBUTION

**MPS** conceived the project and wrote the manuscript with critical contribution from **RM and BB**, who designed, carried and interpreted most of the experimental work with **FB**. **SC** bred all the mice used and helped with experiments. **MMesquita** performed flow cytometry experiments and data analysis.

All authors provided critical feedback and contributed critically to shaping experimental work, analysis and manuscript writing.

## SUPPLEMENTARY FIGURE LEGENDS

**Supplementary Figure 1:**
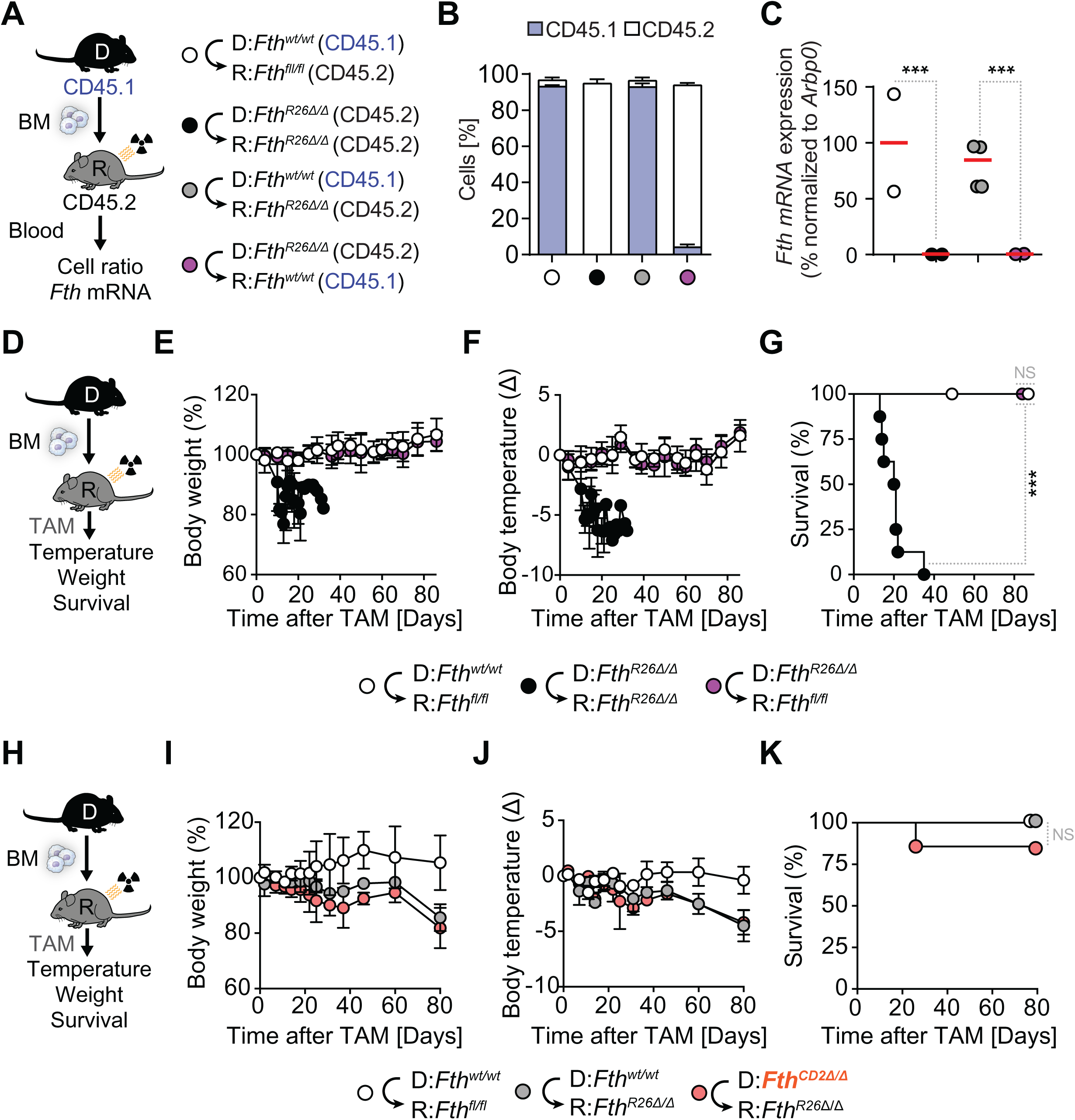
*Fth*-competent chimeric mice, rescue from whole-body *Fth* deletion-induced lethality independently of T-lymphocytes. (**A**) Schematic representation of chimeric mice and engraftment assessment. (**B**) Relative immune cell percentage of donor or recipient origin and (**C**) corresponding *Fth* gene expression in the blood of *Fth^wt/wt^*⇨*Fth^wt/wt^* (n=2), *Fth^R26^*^Δ*/*Δ^⇨*Fth^R26^*^Δ*/*Δ^ (n=2), *Fth^wt/wt^*⇨*Fth^R26^*^Δ*/*Δ^ (n=3) and *Fth^R26^*^Δ*/*Δ^⇨*Fth^wt/wt^* (n=2) chimeric mice, collected on day 10 post-TAM administration, as determined via flow cytometry analysis (B) and qRT-PCR (C). (**D**) Schematic representation of TAM-induced *Fth* deletion in chimeric mice and monitoring vital parameters. Relative body weight (**E**), temperature (**F**) and survival (**G**) of *Fth^fl/fl^*⇨*Fth^fl/fl^*(n=8), *Fth^R26^*^Δ*/*Δ^⇨*Fth^R26^*^Δ*/*Δ^ (n=8) and *Fth^R26^*^Δ*/*Δ^⇨*Fth^fl/fl^* (n=8) chimeric mice following TAM administration on day 0. Data in (E, F) is represented as mean ± SD. Data in (E-G) is pooled from 3 experiments. (**H**) Schematic representation of TAM-induced *Fth* deletion in chimeric mice and monitoring vital parameters. Relative body weight (**I**), temperature (**J**) and survival (**K**) of *Fth^wt/wt^*⇨*Fth^fl/fl^* (n=3), *Fth^wt/wt^*⇨*Fth^R26^*^Δ*/*Δ^ (n=3) and *Fth^CD2^*^Δ*/*Δ^ *Fth^R26^*^Δ*/*Δ^ (n=7) chimeric mice following TAM administration on day 0. Data in (I, J) is represented as mean ± SD. Data in (I-K) is pooled from 2 experiments. One-Way ANOVA with Tukey’s range test for multiple comparison correction was used for comparison between multiple groups. Survival analysis was performed using Log-rank (Mantel-Cox) test. NS: non-significant, *** P<0.001.

**Supplementary Figure 2:**
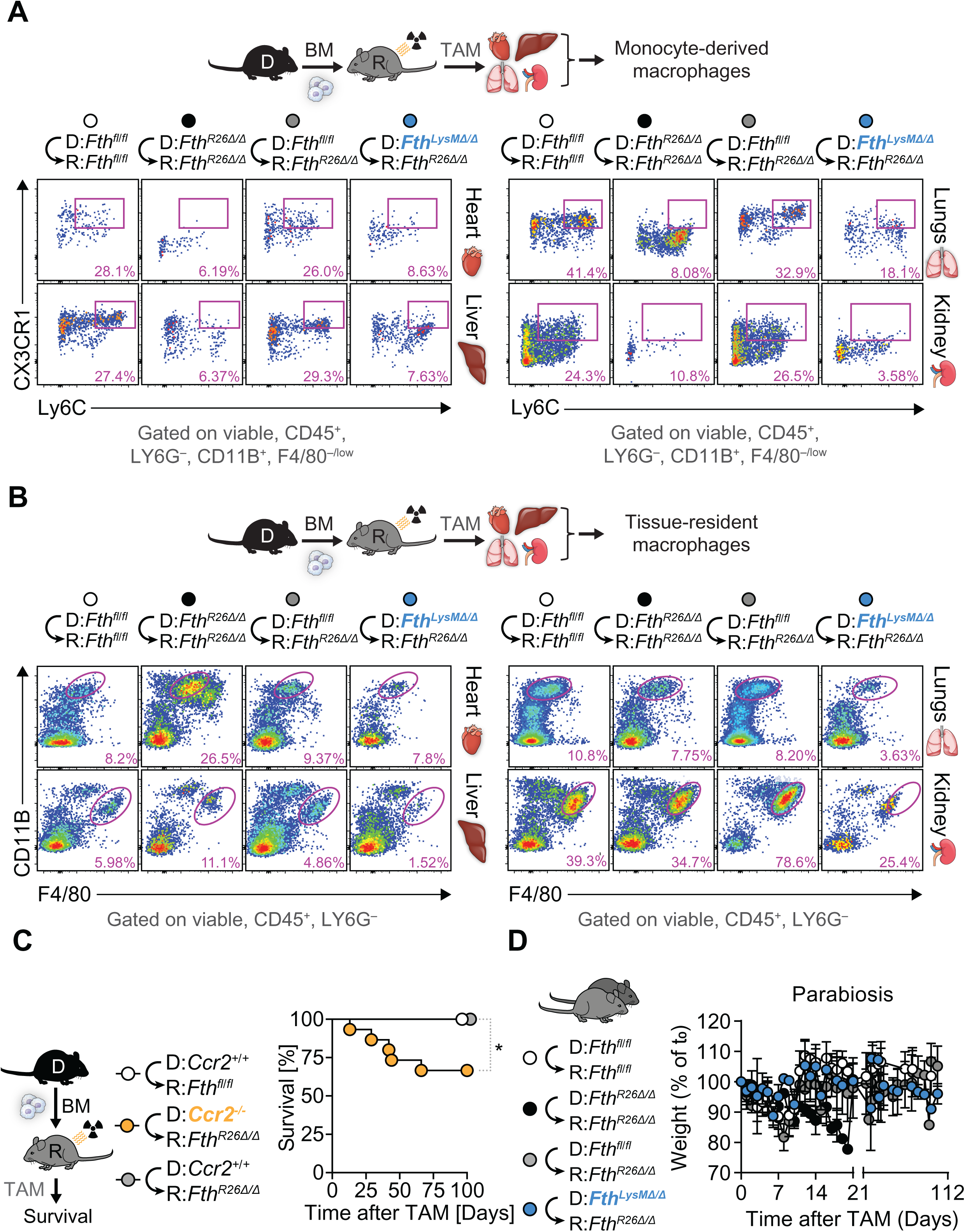
Monocyte-derived macrophages are depleted in chimeric *Fth*-deleted mice. Schematic representation of TAM-induced *Fth* deletion in chimeric mice (day 0) and representative flow cytometry dot plots for (**A**) monocyte-derived macrophage (backgated as Ly6G^−^, CD11b^+^, F4/80^low^) and (**B**) tissue-resident macrophage (Ly6G^−^, CD11b^−/low^, F4/80^high^) populations present in the liver, heart, lungs and kidneys of *Fth^fl/f^*⇨*Fth^fl/fl^*(n=7), *Fth^R26^*^Δ*/*Δ^⇨*Fth^R26^*^Δ*/*Δ^ (n=5), *Fth^fl/fl^*⇨*Fth^R26^*^Δ*/*Δ^ (n=7) and *Fth^LysM^*^Δ*/*Δ^⇨*Fth^R26^*^Δ*/*Δ^ (n=6) chimeric mice, 7 to 19 days post-TAM administration. Data in (A, B) is pooled from 3 independent experiments with similar trends. (**C**) Schematic representation of TAM-induced *Fth* deletion in chimeric mice, and survival of *Ccr2^+/+^*⇨*Fth^fl/fl^* (n=7), *Ccr2^−/−^*⇨*Fth^R26^*^Δ*/*Δ^ (n=6) and *Ccr2^+/+^*⇨⇨*Fth^R26^*^Δ*/*Δ^ (n=15) chimeric mice following TAM administration on day 0. Data in (A) is pooled from 3 independent experiments with similar trends. (**D**) Time course of relative body weight changes of *Fth^fl/fl^*⇔*Fth^fl/fl^* (n=7), *Fth^R26^*^Δ*/*Δ^⇔*Fth^R26^*^Δ*/*Δ^ (n=10), *Fth^fl/fl^*⇔*Fth^R26^*^Δ*/*Δ^ (n=14) and *Fth^LysM^*^Δ*/*Δ^⇔*Fth^R26^*^Δ*/*Δ^ (n=11) parabiotic mouse pairs, following TAM administration on day 0. Data in (D) are presented as mean ±SD, normalized to the initial body weight (t_0_) and pooled from 4 independent experiments with similar trends. Survival analysis was performed using Log-rank (Mantel-Cox) test. NS: non-significant, * P<0.05.

**Supplementary Figure 3:**
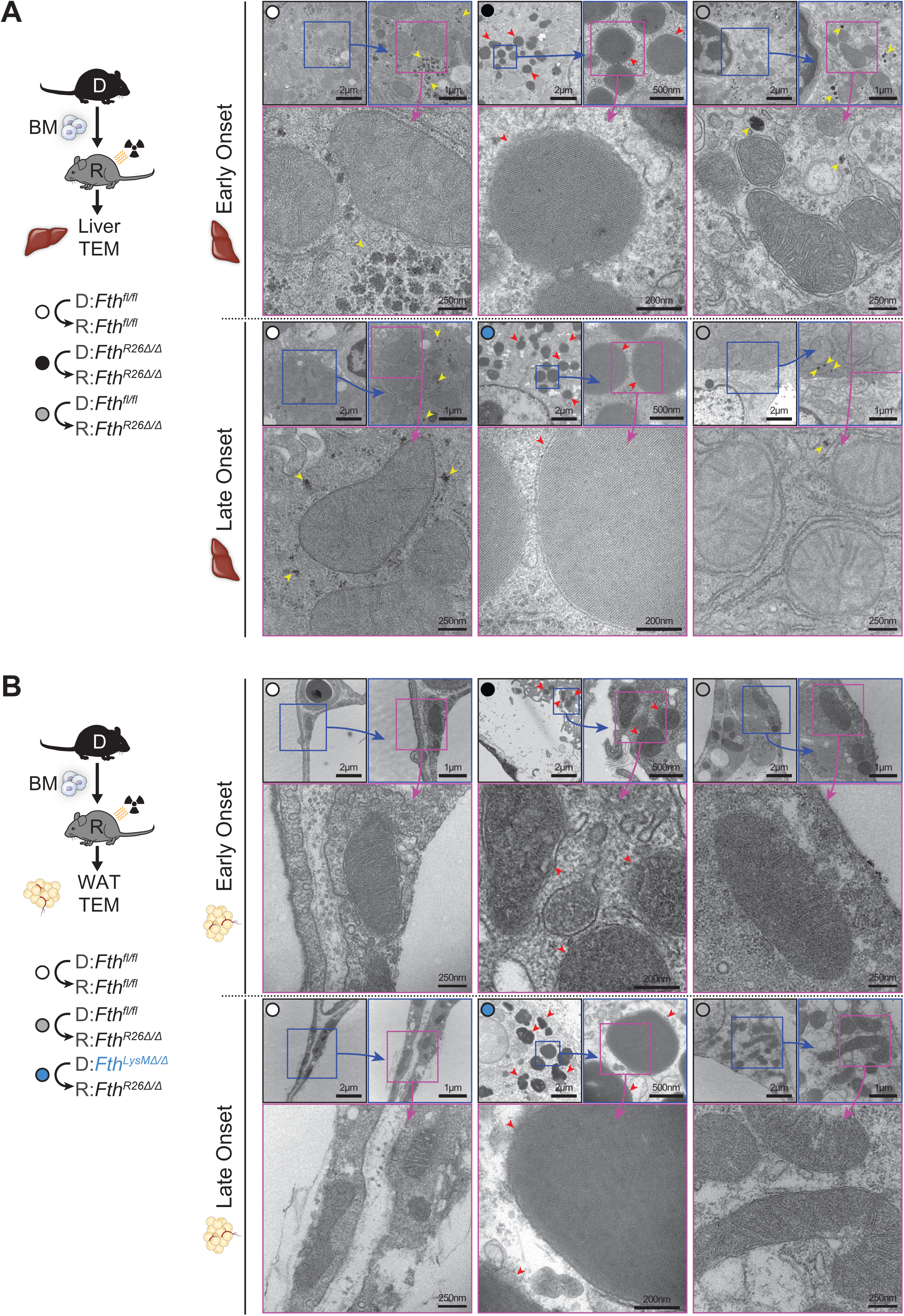
*Fth*-competent myeloid cells prevent the formation of iron-filled siderosomes in chimeric *Fth*-deleted mice. Schematic representation of chimeric mice and TAM-induced *Fth* deletion (day 0) and representative transmission electron microscopy images of siderosome-like structures present in (**A**) hepatocytes (liver) and (**B**) adipocytes (gWAT) from *Fth^R26^*^Δ*/*Δ^⇨*Fth^R26^*^Δ*/*Δ^ chimeric mice, collected on day 8 (early onset), and *Fth^LysM^*^Δ*/*Δ^⇨*Fth^R26^*^Δ*/*Δ^ chimeric mice, collected on day 30 (late onset), but absent in either *Fth^fl/fl^*⇨*Fth^fl/fl^*or *Fth^fl/fl^*⇨*Fth^R26^*^Δ*/*Δ^ chimeric mice, at either time points. Red arrows indicate membrane-bound, electron-dense, siderosome-like structures filled with apparent crystalline iron, consistent with earlier descriptions (Sato *et al*, 1978). Yellow arrows indicate hepatic glycogen granules. Data in (A, B) is representative of 2 independent experiments with similar trends.

**Supplementary Figure 4:**
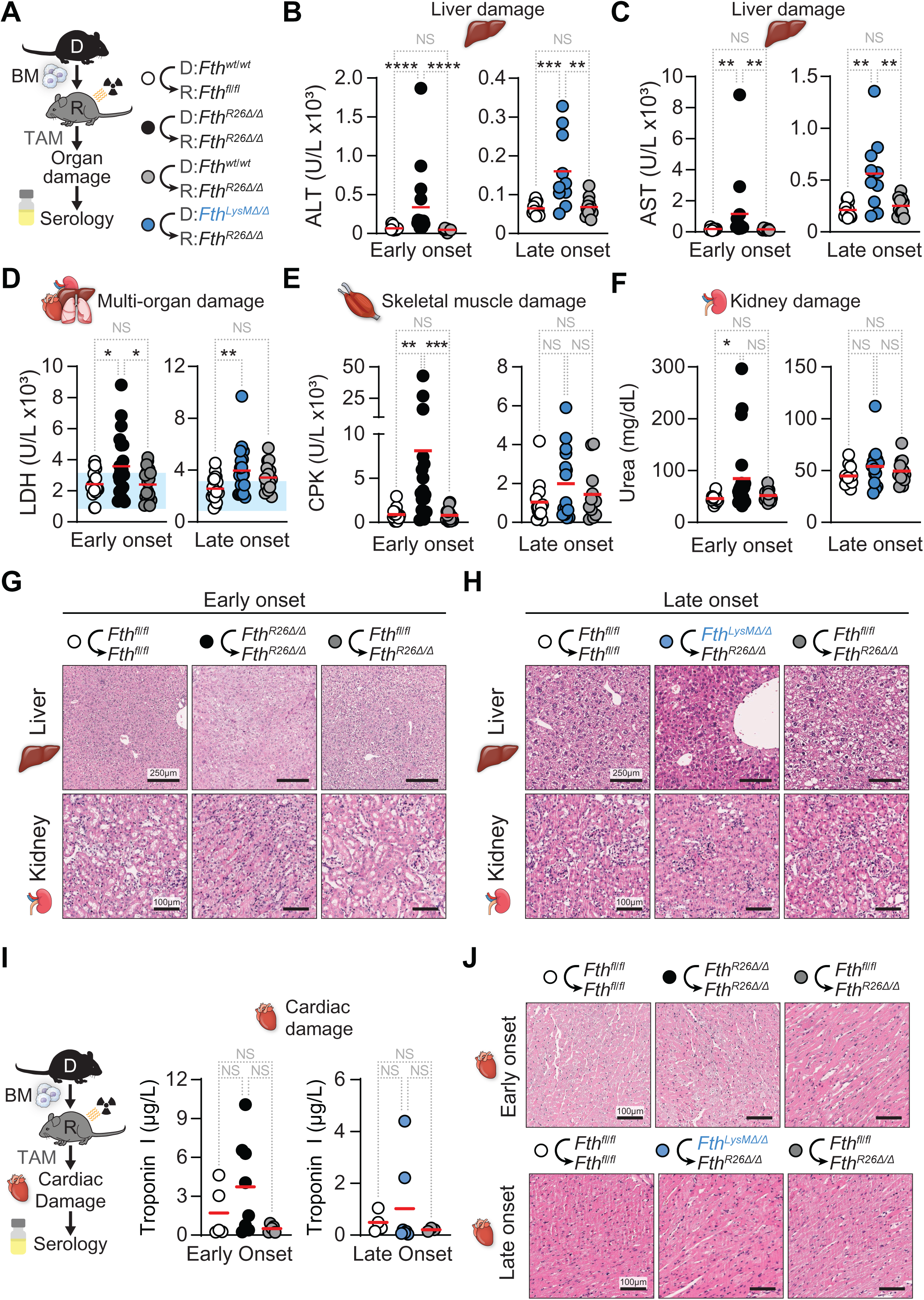
*Fth*-competent myeloid cells support tissue function in chimeric *Fth*-deleted mice. (**A**) Schematic representation of chimeric mice and TAM-induced *Fth* deletion. Plasma levels of (**B**) alanine transaminase (ALT), (**C**) aspartate transaminase (AST), (**D**) lactate dehydrogenase (LDH), (**E**) creatine phosphokinase (CPK) and (**F**) urea, measured in *Fth^fl/fl^*⇨*Fth^fl/fl^*(n=11-21), *Fth^R26^*^Δ*/*Δ^⇨*Fth^R26^*^Δ*/*Δ^ (n=15-25), *Fth^fl/fl^*⇨*Fth^R26^*^Δ*/*Δ^ (n=9-18) and *Fth^LysM^*^Δ*/*Δ^⇨*Fth^R26^*^Δ*/*Δ^ (n=11-20) chimeric mice on days 7-15 (early onset), or 19-35 (late onset) following TAM administration. Data in (B-F) represented as individual values (circles) and mean (red bars) pooled from 4-6 independent experiments with similar trends. (**G**, **H**) Representative hematoxylin and eosin (H&E) stained histology images of liver, kidney and heart from *Fth^fl/fl^*⇨*Fth^fl/fl^*, *Fth^R26^*^Δ*/*Δ^⇨*Fth^R26^*^Δ*/*Δ^, *Fth^fl/fl^*⇨*Fth^R26^*^Δ*/*Δ^ and *Fth^LysM^*^Δ*/*Δ^⇨*Fth^R26^*^Δ*/*Δ^ chimeric mice on day 7 (early onset) (**G**), or 22 (late onset) (**H**) following TAM administration. One-Way ANOVA with Tukey’s range test for multiple comparison correction was used for comparison between multiple groups. NS: non-significant, * P<0.05, ** P<0.01, *** P<0.001, **** P<0.0001.

**Supplementary Figure 5:**
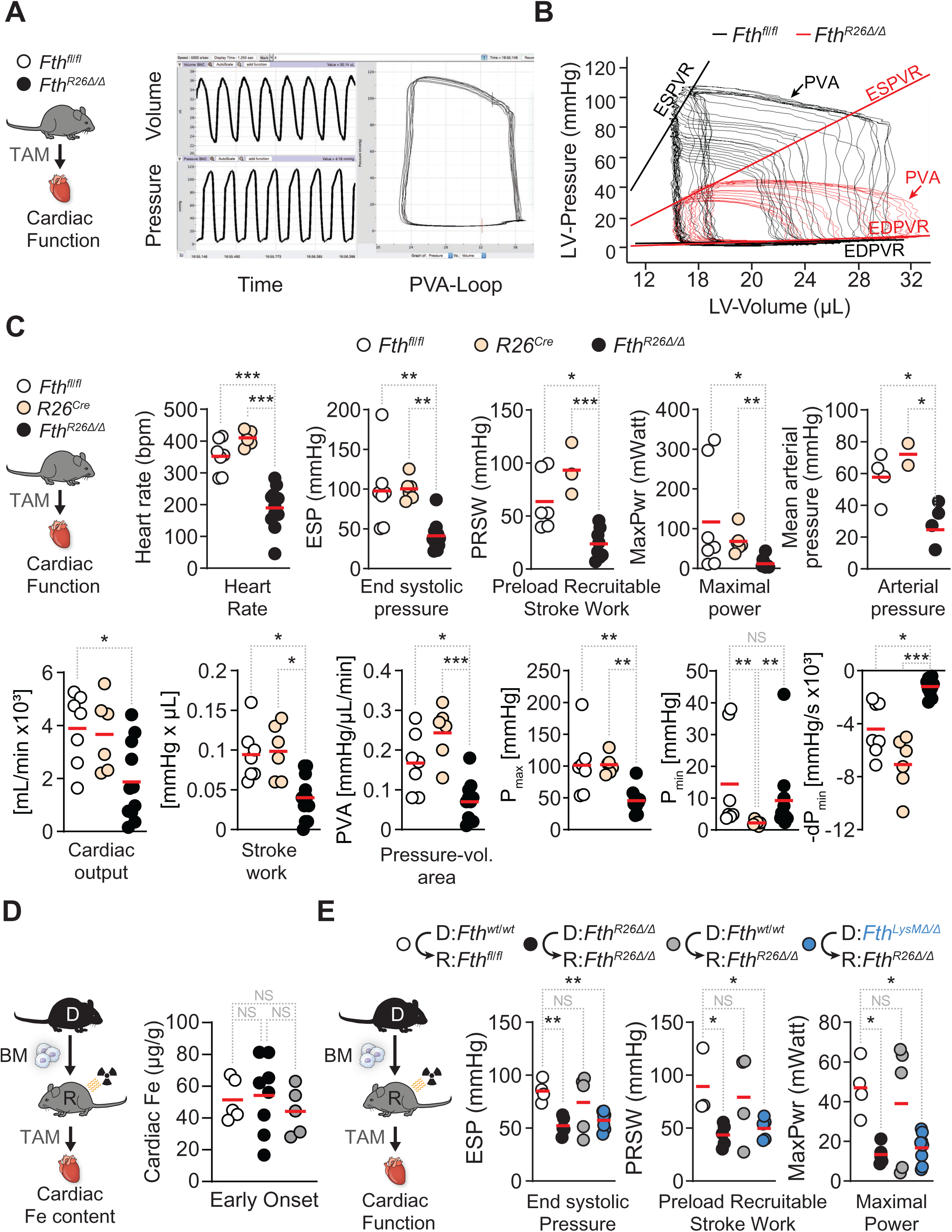
*Fth*-competent myeloid cells support tissue function in chimeric *Fth*-deleted mice. (**A**) Schematic representation of TAM-induced *Fth* deletion, cardiac function assessment and pressure volume loop analysis (PVA loop). (**B**) PVA loop analysis of *Fth^fl/fl^* (n=7) and *Fth^R26^*^Δ*/*Δ^ (n=11) mice, on day **7** post-TAM administration. ESPVR = end systolic pressure-volume relationship, EDPVR = end diastolic pressure-volume relationship. (**C**) Schematic representation of TAM-induced *Fth* deletion (day 0) and cardiac function parameter quantification of *Fth^fl/fl^*(n=7), *R26^Cre^* (n=6) and *Fth^R26^*^Δ*/*Δ^ (n=11) mice, on day 7 post-TAM administration. Data represented as individual values (circles) and mean (red bars). (**D**) Cardiac iron content in *Fth^fl/fl^*⇨*Fth^fl/fl^*(n=3-5), *Fth^R26^*^Δ*/*Δ^ *Fth^R26^*^Δ*/*Δ^ (n=8) and *Fth^fl/fl^*⇨*Fth^R26^*^Δ*/*Δ^ (n=3-5) chimeric mice, 7 days (early onset) following TAM administration. Data pooled from 3 independent experiments with similar trends. (**E**) Schematic representation of chimeric mice and TAM-induced *Fth* deletion (day 0) and cardiac function parameters (end systolic pressure; preload recruitable stroke work; maximal power) in *Fth^fl/fl^*⇨*Fth^fl/fl^* (n=4), *Fth^R26^*^Δ*/*Δ^⇨*Fth^R26^*^Δ*/*Δ^ (n=5), *Fth^fl/fl^*⇨*Fth^R26^*^Δ*/*Δ^ (n=5) and *Fth^LysMΔ/Δ^*⇨*Fth^R26Δ/Δ^* (n=8) chimeric mice between days 10-51 following TAM administration. Data pooled from 2 independent experiments with similar trends. Data in (C-E) presented as individual values (circles) and mean (red bars). One-Way ANOVA with Tukey’s range test for multiple comparison correction was used for comparison between multiple groups. NS: non-significant, * P<0.05, ** P<0.01, *** P<0.001.

**Supplementary Figure 6:**
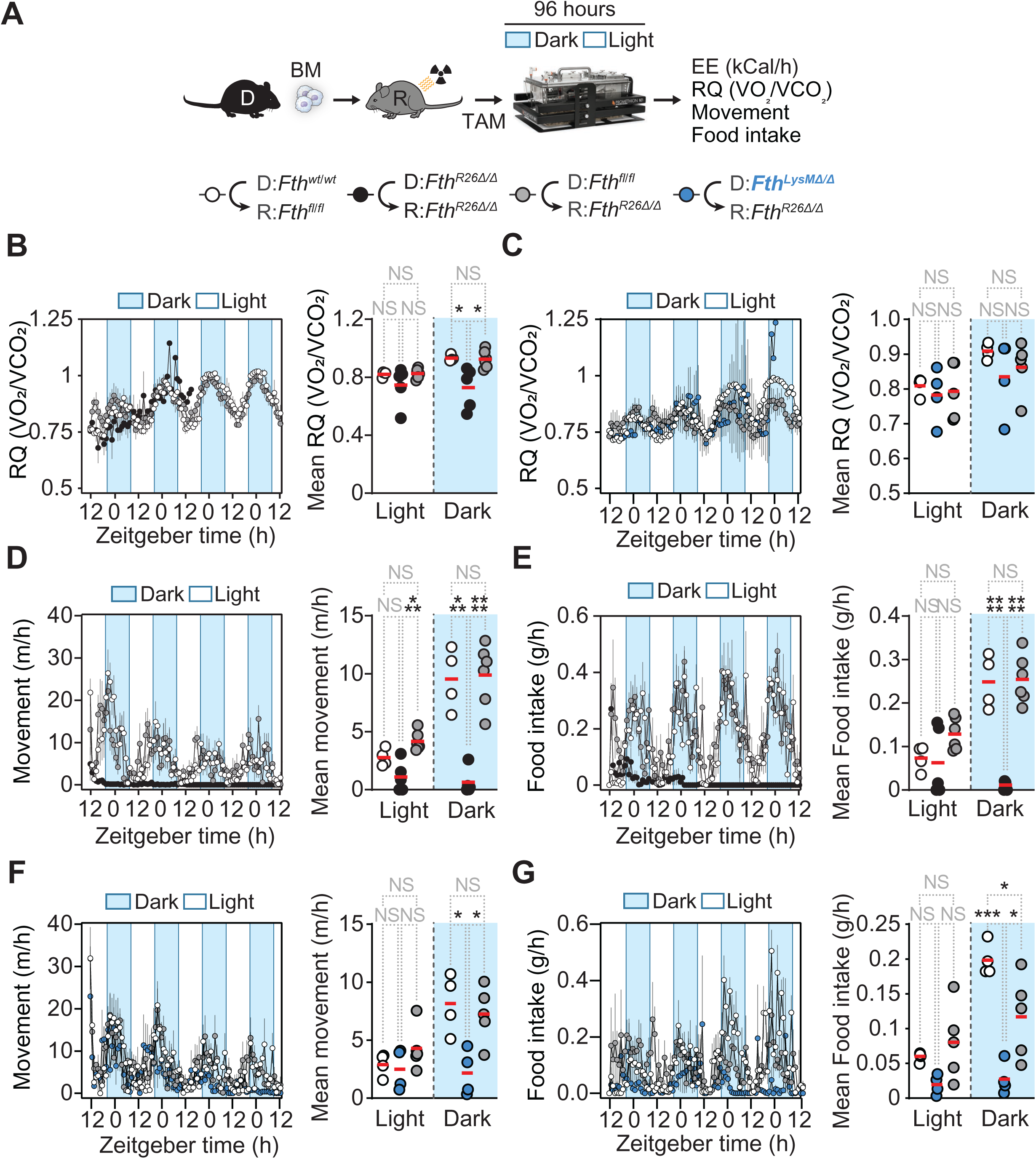
*Fth*-competent myeloid cells restore movement and food intake in chimeric *Fth*-deleted mice. (**A**) Schematic representation of TAM-induced *Fth* deletion in chimeric mice (day 0) and metabolic cage assessment of metabolic parameters. (**B**, **C**) Time course of respiratory quotient (RQ, calculated as VO_2_/VCO_2_), and mean (red bars) RQ during daytime/nighttime (dot plots) of *Fth^fl/fl^*⇨*Fth^fl/fl^*(n=4), *Fth^R26^*^Δ*/*Δ^⇨*Fth^R26^*^Δ*/*Δ^ (n=5), *Fth^fl/fl^*⇨*Fth^R26^*^Δ*/*Δ^ (n=5-6) and *Fth^LysM^*^Δ*/*Δ^⇨*Fth^R26^*^Δ*/*Δ^ (n=4) chimeric mice, assessed from day 7 (**B**; early onset), or day 20 (**C**; late onset) post TAM administration. Time course and mean (red bars) of daytime/nighttime values (dot plots) for mouse movement (m/h; **D**), and rate of food intake (g/h; **E**) of *Fth^fl/fl^*⇨*Fth^fl/fl^*(n=4), *Fth^R26^*^Δ*/*Δ^⇨*Fth^R26^*^Δ*/*Δ^ (n=5) and *Fth^fl/fl^*⇨*Fth^R26^*^Δ*/*Δ^ (n=6) chimeric mice, assessed from day 7 (early onset). Time course and mean (red bars) of daytime/nighttime values (dot plots) for mouse movement (m/h; **F**), and rate of food intake (g/h; **G**) of *Fth^fl/fl^*⇨*Fth^fl/fl^*(n=4), *Fth^LysM^*^Δ*/*Δ^⇨*Fth^R26^*^Δ*/*Δ^ (n=4) and *Fth^fl/fl^*⇨*Fth^R26^*^Δ*/*Δ^ (n=5) chimeric mice, assessed from day 20 (late onset). Data in (B-G) is displayed as mean ± SD (time course), or as individual values (circles) and mean (red bars) (dot plots). Data in (B-G) is pooled from 2 independent experiments with similar trend. One-Way ANOVA with Tukey’s range test for multiple comparison correction was used for comparison between multiple groups. NS: non-significant, * P<0.05, *** P<0.001, **** P<0.0001.

**Supplementary Figure 7:**
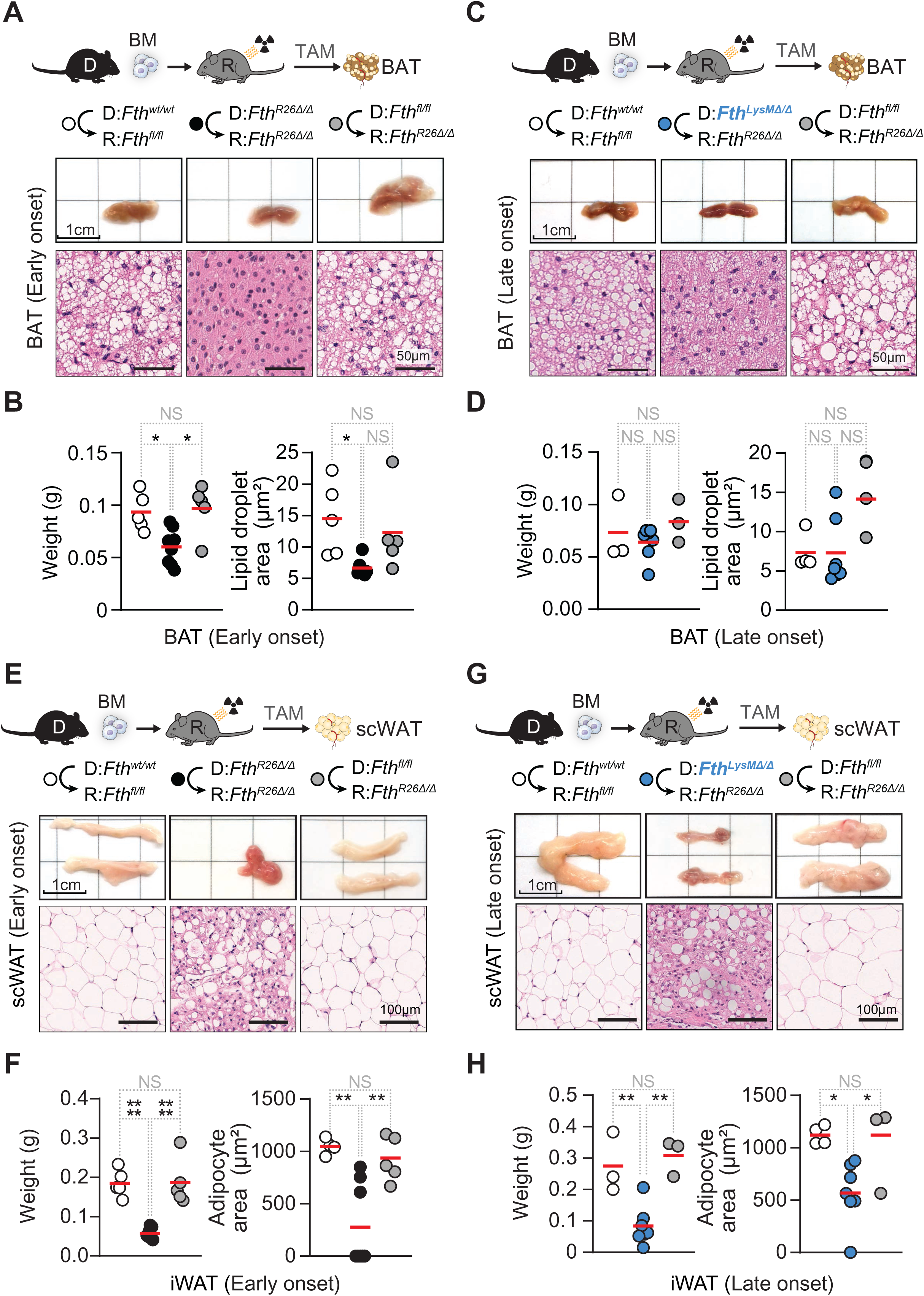
*Fth*-competent myeloid cells support BAT and WAT function in chimeric *Fth*-deleted mice. (**A**) Schematic representation of chimeric mice and TAM-induced *Fth* deletion, and representative macroscopic and histological images of brown adipose tissue pads (BAT) of *Fth^fl/fl^*⇨*Fth^fl/fl^* (n=5), *Fth^R26^*^Δ*/*Δ^⇨*Fth^R26^*^Δ*/*Δ^ (n=8) and *Fth^fl/fl^*⇨*Fth^R26^*^Δ*/*Δ^ (n=5) chimeric mice, collected on day 7 (early onset) post TAM administration. (**B**) BAT pad weight (left) and mean (red bars) BAT adipocyte lipid droplet area (right) of *Fth^fl/fl^*⇨*Fth^fl/fl^* (n=5), *Fth^R26^*^Δ*/*Δ^⇨*Fth^R26^*^Δ*/*Δ^ (n=8) and *Fth^fl/fl^*⇨*Fth^R26^*^Δ*/*Δ^ (n=5) chimeric mice, collected on day 7 (early onset) post TAM administration. (**C**) Schematic representation of chimeric mice and TAM-induced *Fth* deletion, and representative macroscopic and histological images of BAT of *Fth^fl/fl^*⇨*Fth^fl/fl^* (n=3-4), *Fth^LysM^*^Δ*/*Δ^⇨*Fth^R26^*^Δ*/*Δ^ (n=6) and *Fth^fl/fl^*⇨*Fth^R26^*^Δ*/*Δ^ (n=3-4) chimeric mice, collected between days 16 and 39 (late onset) post TAM administration. (**D**) BAT pad weight (left) and mean (red bars) BAT adipocyte lipid droplet area (right) of *Fth^fl/fl^*⇨*Fth^fl/fl^*(n=3-4), *Fth^LysM^*^Δ*/*Δ^⇨*Fth^R26^*^Δ*/*Δ^ (n=6) and *Fth^fl/fl^*⇨*Fth^R26^*^Δ*/*Δ^ (n=3-4) chimeric mice, collected between days 16 and 39 (late onset) post TAM administration. Data in (B, D) represented as individual values (circles) and mean (red bars), pooled from 3 independent experiments with similar trends. (**E**) Schematic representation of chimeric mice and TAM-induced *Fth* deletion, and representative macroscopic and histological images of inguinal white adipose tissue pads (iWAT) of *Fth^fl/fl^*⇨*Fth^fl/fl^* (n=4-5), *Fth^R26^*^Δ*/*Δ^⇨*Fth^R26^*^Δ*/*Δ^ (n=8) and *Fth^fl/fl^*⇨*Fth^R26^*^Δ*/*Δ^ (n=5) chimeric mice, collected on day 7 (early onset) post TAM administration. (**F**) iWAT pad weight (left) and mean (red bars) iWAT adipocyte area (right) of *Fth^fl/fl^*⇨*Fth^fl/fl^* (n=4-5), *Fth^R26^*^Δ*/*Δ^⇨*Fth^R26^*^Δ*/*Δ^ (n=8) and *Fth^fl/fl^*⇨*Fth^R26^*^Δ*/*Δ^ (n=5) chimeric mice, collected on day 7 (early onset) post TAM administration. (**G**) Schematic representation of chimeric mice and TAM-induced *Fth* deletion, and representative macroscopic and histological images of iWAT of *Fth^fl/fl^*⇨*Fth^fl/fl^*(n=3-4), *Fth^LysM^*^Δ*/*Δ^⇨*Fth^R26^*^Δ*/*Δ^ (n=7) and *Fth^fl/fl^*⇨*Fth^R26^*^Δ*/*Δ^ (n=3) chimeric mice, collected between days 16 and 39 (late onset) post TAM administration. (**H**) iWAT pad weight (left) and mean (red bars) iWAT adipocyte area (right) of *Fth^fl/fl^*⇨*Fth^fl/fl^*(n=3-4), *Fth^LysM^*^Δ*/*Δ^⇨*Fth^R26^*^Δ*/*Δ^ (n=7) and *Fth^fl/fl^*⇨*Fth^R26^*^Δ*/*Δ^ (n=3) chimeric mice, collected between days 16 and 39 (late onset) post TAM administration. Data in (F, H) represented as individual values (circles) and mean (red bars), pooled from 3 independent experiments with similar trends. One-Way ANOVA with Tukey’s range test for multiple comparison correction was used for comparison between multiple groups. NS: non-significant, * P<0.05, ** P<0.01, **** P<0.0001.

**Supplementary Figure 8:**
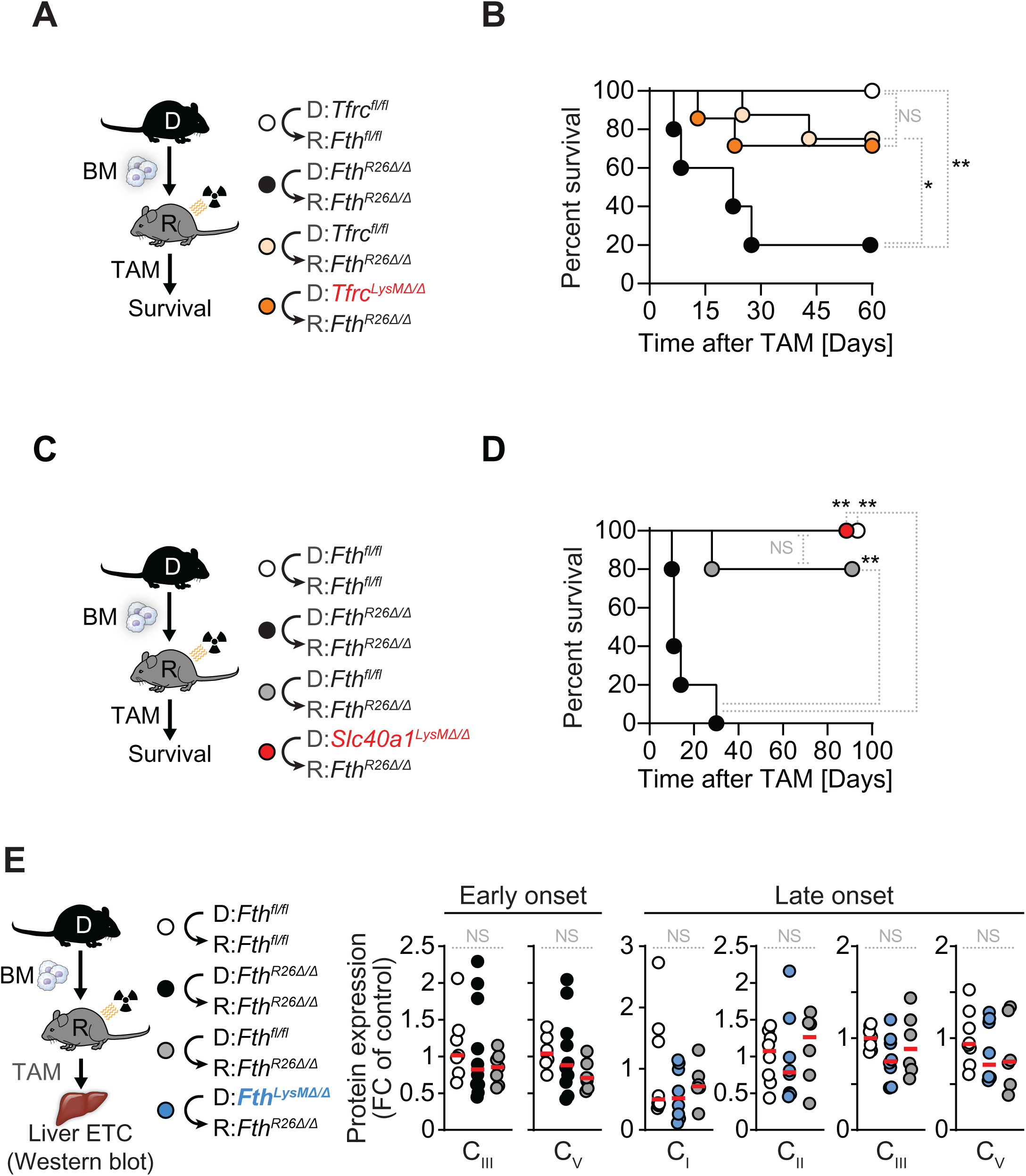
Myeloid cells rescue chimeric *Fth*-deleted mice irrespectively of cellular Fe import/export. (**A**) Schematic representation of TAM-induced *Fth* deletion in chimeric mice, and (**B**) survival of *Tfrc^fl/fl^*⇨*Fth^fl/fl^*(n=7), *Fth^R26^*^Δ*/*Δ^⇨*Fth^R26^*^Δ*/*Δ^ (n=5), *Tfrc^fl/fl^*⇨*Fth^R26^*^Δ*/*Δ^ (n=8) and *Tfrc^LysM^*^Δ*/*Δ^⇨*Fth^R26^*^Δ*/*Δ^ (n=7) chimeric mice following TAM administration. Data in (B) is pooled from 2 independent experiments with similar trends. (**C**) Schematic representation of TAM-induced *Fth* deletion in chimeric mice, and (**D**) survival of *Fth^fl/fl^*⇨*Fth^fl/fl^* (n=5), *Fth^R26^*^Δ*/*Δ^⇨*Fth^R26^*^Δ*/*Δ^ (n=5), *Fth^fl/fl^*⇨*Fth^R26^*^Δ*/*Δ^ (n=5), and *Slc40a1^LysM^*^Δ*/*Δ^⇨*Fth^R26^*^Δ*/*Δ^ (n=5) chimeric mice on day 0. (**E**) quantification of ETC subunits for complexes I-V in the livers of *Fth^fl/fl^*⇨*Fth^fl/fl^* (n=7), *Fth^R26^*^Δ*/*Δ^⇨*Fth^R26^*^Δ*/*Δ^ (n=10) and *Fth^fl/fl^*⇨*Fth^R26^*^Δ*/*Δ^ (n=7) chimeric mice, between days 7-8 (early onset), or from *Fth^fl/fl^*⇨*Fth^fl/fl^*(n=8), *Fth^LysM^*^Δ*/*Δ^⇨*Fth^R26^*^Δ*/*Δ^ (n=8) and *Fth^fl/fl^*⇨*Fth^R26^*^Δ*/*Δ^ (n=6) chimeric mice, between days 19 and 22 (late onset). Data in (E) is pooled from 4 independent experiments with similar trends. One-Way ANOVA with Tukey’s range test for multiple comparison correction was used for comparison between multiple groups. Survival analysis was performed using Log-rank (Mantel-Cox) test. NS: non-significant, * P<0.05, ** P<0.01.

**Supplementary Figure 9:**
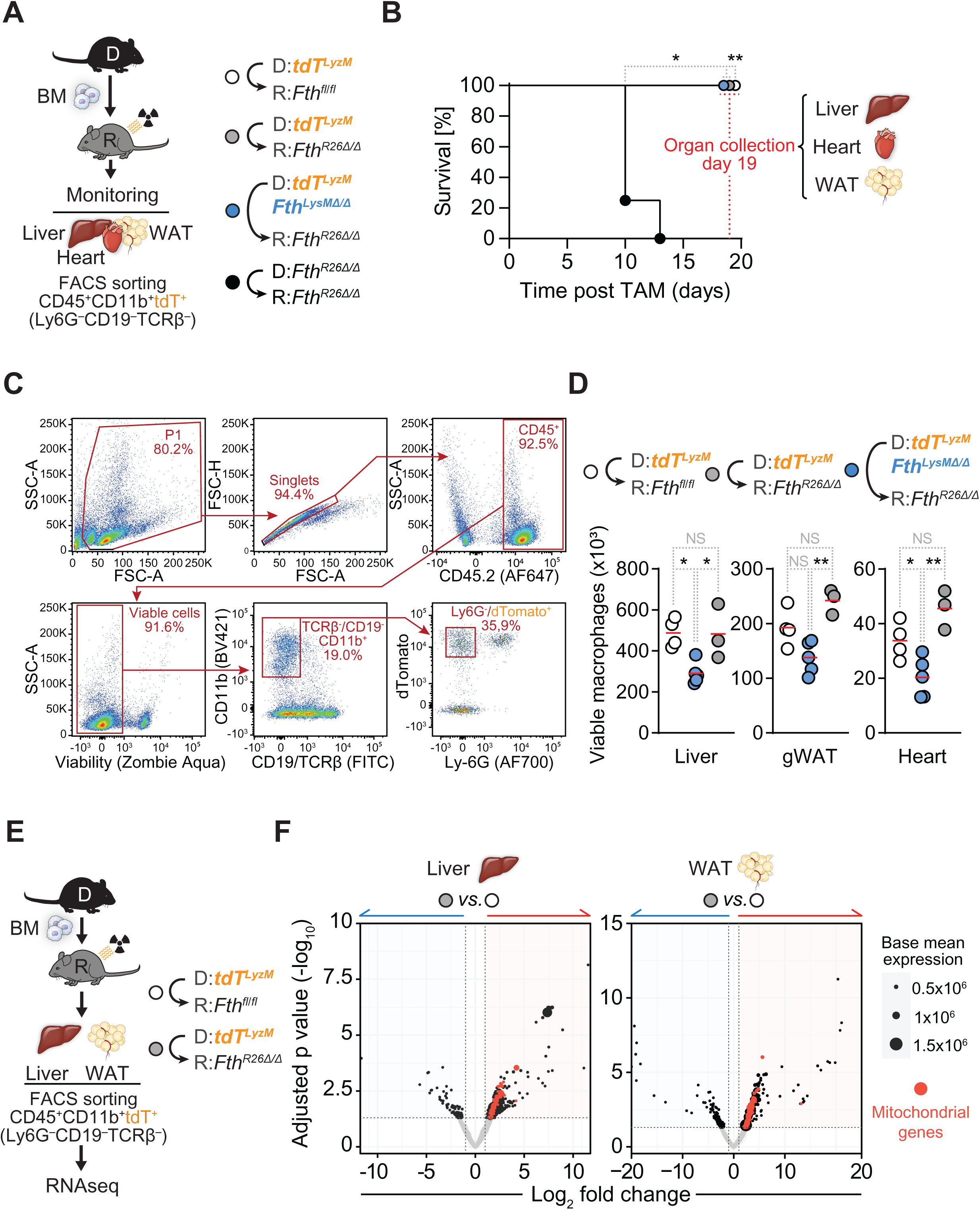
RNA sequencing analysis of monocyte/macrophages in chimeric *Fth*-deleted mice. (**A**) Schematic representation of chimeric mice, TAM administration and fluorescence-activated cell sorting (FACS) of *LysM^+^*monocyte/macrophages (CD45^+^,CD11b^+^,Ly6G^−^,CD19^−^,TCRβ^−^) in liver, heart and WAT. (**B**) Survival of *tdT^LysM^*⇨*Fth^fl/fl^*(n=4), *tdT^LysM^*⇨*Fth^R26^*^Δ*/*Δ^ (n=3), *tdT^LysM^Fth^LysM^*^Δ*/*Δ^⇨*Fth^R26^*^Δ*/*Δ^ (n=5) and *Fth^R26^*^Δ*/*Δ^⇨*Fth^R26^*^Δ*/*Δ^ (n=4) chimeric mice until organ collection on day 19 post-TAM administration. (**C**) Gating strategy for sorting *LysM^+^* monocyte/macrophages (CD45^+^, CD11b^+^, Ly6G^−^, CD19^−^, TCRβ^−^) in liver, WAT and Heart. (**D**) Number of viable *LysM^+^* monocyte/macrophages (CD45^+^, CD11b^+^, Ly6G^−^, CD19^−^, TCRβ^−^) in the liver, gWAT and heart of *tdT^LysM^*⇨*Fth^fl/fl^* (n=4), *tdT^LysM^*⇨*Fth^R26^*^Δ*/*Δ^ (n=3) and *tdT^LysM^Fth^LysM^*^Δ*/*Δ^⇨*Fth^R26^*^Δ*/*Δ^ (n=5) chimeric mice on day 19 post-TAM administration. (**E**) Schematic representation of chimeric mice, TAM administration and fluorescence-activated cell sorting (FACS) of *LysM^+^* monocyte/macrophages (CD45^+^, CD11b^+^, Ly6G^−^, CD19^−^, TCRβ^−^) in liver and WAT. (**F**) Volcano plots of differentially regulated genes between *LysM^+^* monocyte/macrophages sorted from liver (left) or WAT (right) of *tdT^LysM^*⇨*Fth^R26^*^Δ*/*Δ^ (n=3), *tdT^LysM^*⇨*Fth^fl/fl^*(n=4) chimeric mice, on day 19 post TAM administration. Red dots depict mitochondrial genes that are significantly differentially regulated. One-Way ANOVA with Tukey’s range test for multiple comparison correction was used for comparison between multiple groups. Survival analysis was performed using Log-rank (Mantel-Cox) test. NS: non-significant, * P<0.05, ** P<0.01.

**Supplementary Figure 10:**
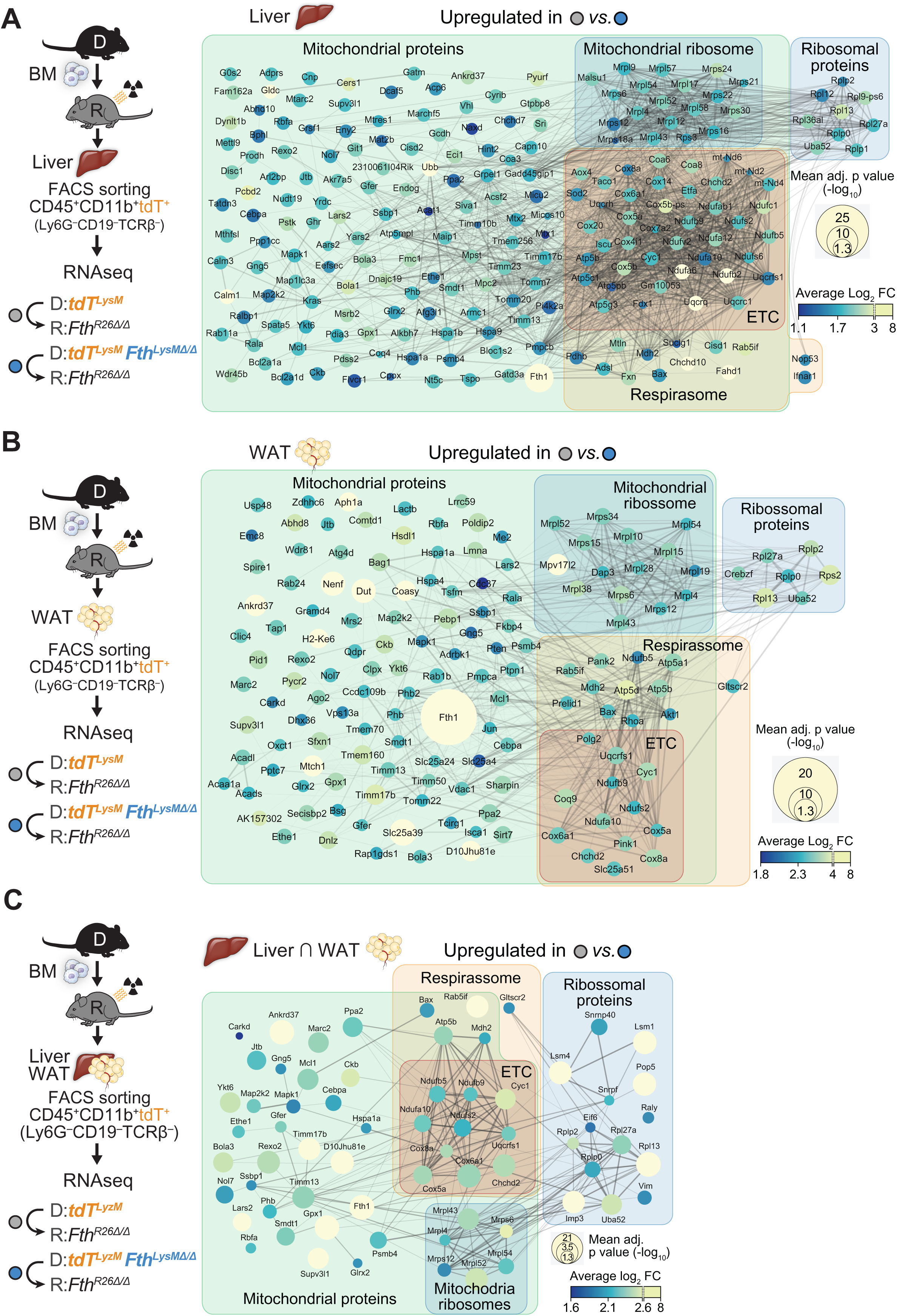
Mitochondrial gene expression program upregulated by *Fth*-competent monocyte-derived macrophages in *Fth*-deleted chimeras. (**A-C**) Schematic representations of chimeric mice, TAM administration and fluorescence-activated cell sorting (FACS) of *LysM^+^* monocyte/macrophages (CD45^+^,CD11b^+^,Ly6G^−^,CD19^−^,TCRβ^−^) and STRING database (STRING-DB) interaction networks of mitochondrial, and mitochondria-related genes (as per gene ontology analysis) significantly upregulated in *LysM^+^* monocyte-derived macrophages from *tdT^LysM^*⇨*Fth^R26^*^Δ*/*Δ^ chimeric mice, as compared to *tdT^LysM^Fth^LysM^*^Δ*/*Δ^⇨*Fth^R26^*^Δ*/*Δ^ chimeric mice in either (**A**) liver or (**B**) WAT. (**C**) Mitochondrial and mitochondria-related genes that are significantly upregulated in *LysM^+^* monocyte-derived macrophages from both the liver and WAT (intersection of significantly upregulated genes) of *tdT^LysM^*⇨*Fth^R26^*^Δ*/*Δ^ chimeric mice, as compared to *tdT^LysM^Fth^LysM^*^Δ*/*Δ^ *Fth^R26^*^Δ*/*Δ^ chimeric mice. In (A-C), known interactions annotated in the STRING-DB are depicted by edges connecting gene dots.

**Supplementary Figure 11:**
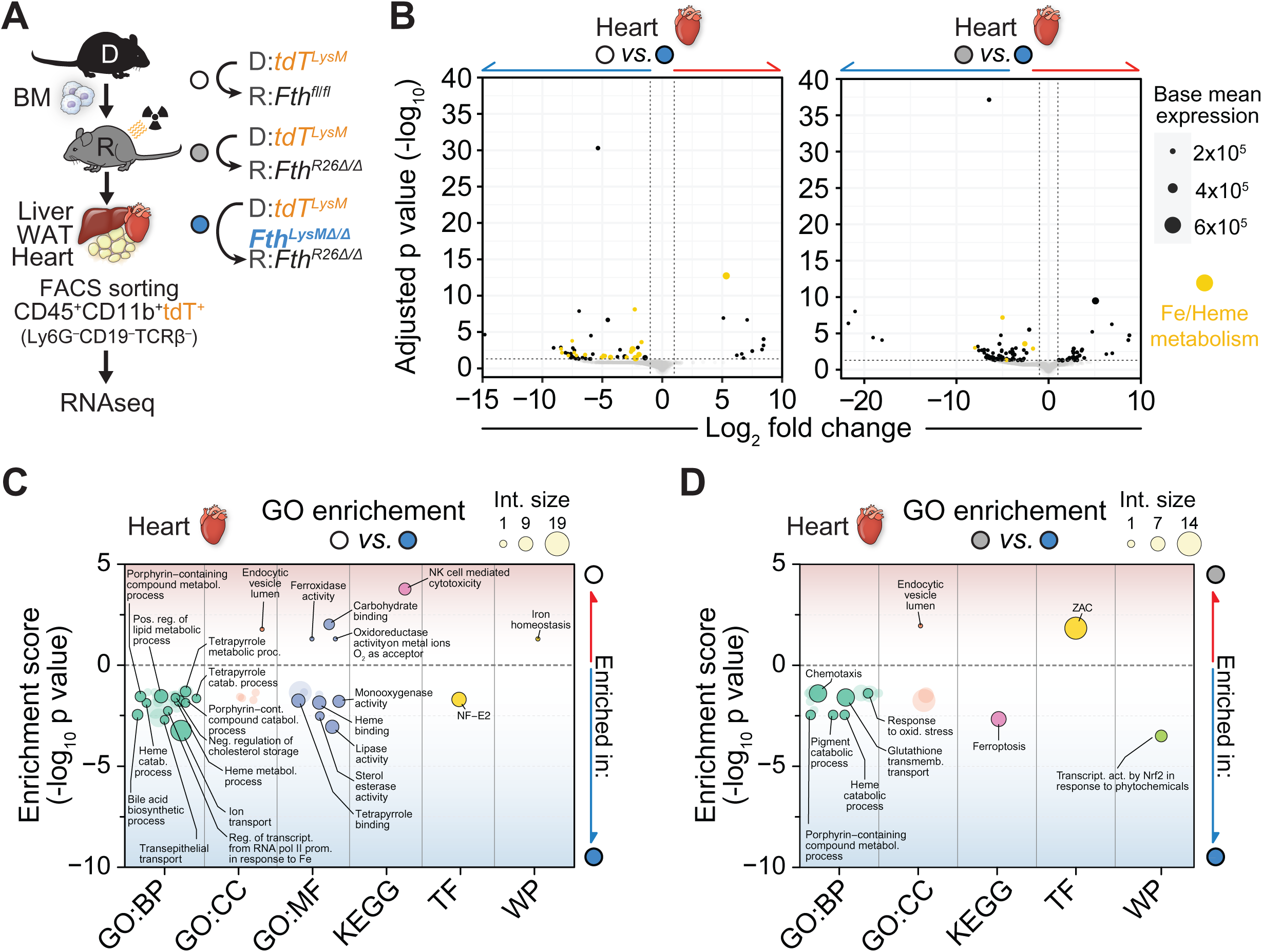
*Fth*-competent monocyte-derived macrophages in the heart of *Fth*-deleted chimeras do not employ a mitochondrial gene transcriptional program. (**A**) Schematic representation of chimeric mice, TAM administration and fluorescence-activated cell sorting (FACS) of *LysM^+^* monocyte/macrophages (CD45^+^, CD11b^+^, Ly6G^−^, CD19^−^, TCRβ^−^) in liver, WAT and heart. (**B**) Volcano plots of differentially regulated genes between *LysM^+^* monocyte/macrophages sorted from the heart of *tdT^LysM^Fth^LysM^*^Δ*/*Δ^,:=;*Fth^R26^*^Δ*/*Δ^ (n=5) vs. *tdT^LysM^*,:=;*Fth^fl/fl^*(n=4; left) or *tdT^LysM^*,:=;*Fth^R26^*^Δ*/*Δ^ (n=3; right) chimeric mice, on day 19 post TAM administration. Yellow dots depict genes involved in iron/heme metabolism that are significantly differentially regulated. (**C**) Gene ontology analysis depicting ontologies that are significantly enriched comparing *LysM^+^* monocyte/macrophages sorted from heart of *tdT^LysM^Fth^LysM^*^Δ*/*Δ^,:=;*Fth^R26^*^Δ*/*Δ^ (n=5) chimeras, vs. *tdT^LysM^*,:=;*Fth^fl/fl^*(n=4; left) or *tdT^LysM^*,:=;*Fth^R26^*^Δ*/*Δ^ (n=3; right) chimeric mice on day 19 post-TAM administration. Ontologies significantly enriched in *LysM^+^* monocyte/macrophages from *tdT^LysM^*,:=;*Fth^fl/fl^* are depicted as: enrichment score -log_10_ p value > 1,301. Ontologies significantly enriched in *LysM^+^* monocyte/macrophages from *tdT^LysM^Fth^LysM^*^Δ*/*Δ^,:=;*Fth^R26^*^Δ*/*Δ^ or *tdT^LysM^*,:=;*Fth^R26^*^Δ*/*Δ^ chimeras are depicted as: enrichment score -log_10_ p value < -1,301. Ontology classes: TF = transcription factors; GO:CC = gene ontology : cellular component; GO:MF = GO : molecular function; GO:BP = GO: biological process; KEGG = Kyoto Encyclopedia of Genes and Genomes pathway; WP = Wiki Pathways.

## MATERIALS AND METHODS

### Animals

Mice were bred and maintained under specific pathogen-free (SPF) conditions at the Gulbenkian Institute for Molecular Medicine (GIMM). All experimental protocols were approved by the Ethics Committee of the IGC, the “Órgão Responsável pelo Bem-estar dos Animais” (ORBEA) (license A009/2011 and A006-2022) and the Portuguese National Entity (Direcção Geral de Alimentação e Veterinária) . Experimental procedures were performed according to the Portuguese (Portaria no. 1005/92, Decreto-Lei no. 113/2013 and Decreto-lei no.1/2019) and European (Directive 2010/63/EU) legislations, concerning housing, husbandry, and animal welfare. *R26^CreERT2^Fth*^Δ*/*Δ^; *LysM^Cre^Fth*^Δ*/*Δ^; *CD2^Cre^Fth*^Δ*/*Δ^ and *Cx3CR1^Cre^Fth*^Δ*/*Δ^ mice were generated by crossing C57BL/6 *Fth^fl/fl^* mice obtained from Prof. Lukas Kuhn (ETH, Switzerland) with C57BL/6 *R26^CreERT2^*, *LysM^Cre^*, *CD2^Cre^* and *Cx3cr1^Cre^* mice, respectively. *LysM^Cre^Tfrc*^Δ*/*Δ^ and *LysM^Cre^Slc40a1^fl/fl^* mice were generated by crossing B6.129S *Tfrc^fl/fl^* and C57BL/6 *Slc40a1^fl/fl^* mice with *LysM^Cre^* mice, respectively. *R26^tdTomato/CreERT2^* and *LysM^Cre^ R26^tdTomato^* mice were generated by crossing *R26^tdTomato^* mice with *R26^CreERT2^*and *LysM^Cre^* mice, respectively. *R26^tdTomato/CreERT2^Fth*^Δ*/*Δ^ and *LysM^Cre^ R26^tdTomato^Fth*^Δ*/*Δ^ mice were generated by further crossing *R26^tdTomato/CreERT2^* and *LysM^Cre^ R26^tdTomato^* with C57BL/6 *Fth^fl/fl^*mice. *Fth^V5^* mice contain an allele encoding the FTH protein fused to a V5 epitope at its N-terminal end, just downstream of its natural ATG. This allele was generated through CRISPR/CaS9-mediated homologous recombination. The sgRNA was generated by *in vitro* transcription from the “gRNA-V5-*Fth*” plasmid. This plasmid was built by hybridizing oligonucleotides “V5-*Fth*-gRNA-up” and “V5-*Fth*-gRNA-down” (see *Table 2)* and introduced into the *BbsI* sites of the gRNA basic plasmid {Casaca, 2016 #903}. The gRNA-V5-*Fth* plasmid was linearized and transcribed with T7 RNA polymerase using the MEGAshortscript^TM^ T7 Transcription kit (Thermo Fisher cat. #AM1354) and purified with the MEGAclear^TM^ Transcription Clean-up kit (Thermo Fisher cat. #AM1908). The replacement single stranded DNA oligonucleotide “V5-*Fth* replace” (see ***Table 2***) containing the V5 coding region flanked by genomic 60 nucleotide homology sequences was obtained from IDT. To generate the *Fth^V5^* mice, a mix containing V5-*Fth*-sgRNA (10 ng/μl), *CaS9* mRNA (10 ng/μl), and the V5-*Fth* replace oligo (10 ng/μl) was introduced into fertilized C57BL/6 mouse oocytes by pronuclear microinjection. Identification of the recombinant allele was done by PCR on genomic DNA purified from tail biopsies using the oligonucleotide pair “*Fth*-Chk-Fwd” and “*Fth*-Chk-Rev” (see ***Table 2***) that amplifies the relevant genomic area from both the wild type and the V5-tagged alleles (160 and 205 bps, respectively). The PCR fragments were separated by electrophoresis in a 15% polyacrylamide/TBE gel, the band corresponding to the V5-containing allele recovered, and its sequence confirmed. *R26^PhAM/CreERT2^* and *LysM^Cre^R26^PhAM^*mice were generated by crossing *R26^PhAM^* mice with *R26^CreERT2^* and *LysM^Cre^* mice, respectively. *R26^PhAM/CreERT2^Fth*^Δ*/*Δ^ and *LysM^Cre^R26^PhAM^Fth*^Δ*/*Δ^ mice were generated by further crossing *R26^PhAM/CreERT2^* and *LysM^Cre^R26^PhAM^*mice with C57BL/6 *Fth^fl/fl^* mice.

### Bone Marrow Chimeras

All bone marrow chimera combinations were generated by lethally irradiating (8.5 Gy) recipient mice and reconstituted 4h post-irradiation with freshly isolated or cryo-preserved bone marrow cells isolated from donor mice (retroorbital injection of 2∼3×10^6^ cells in 100 μL RPMI). Successful hematopoietic cell reconstitution was confirmed 6-8 weeks after bone marrow transfer via flow cytometry using differential CD45.1/CD45.2 haplotype markers and corresponding to cells derived from donor or recipient hematopoietic progenitors, respectively, or by concomitant lethal irradiation of control mice that were not reconstituted with bone marrow cells.

### Tamoxifen treatment

Conditional *Fth^fl/fl^* deletion (Δ) in bone marrow chimeras generated with either *R26^CreERT2^Fth*^Δ*/*Δ^ recipient and/or donor mice was induced at 6-10 weeks post-transplantation by oral gavage of tamoxifen (Sigma-Aldrich, Cat. #T5648; 50 mg/kg body weight in 100µL Corn Oil (Sigma, Cat. #C8267)/5% EtOH; 3x every other day). Conditional deletion of the *Fth^fl/fl^* allele in *R26^CreERT2^Fth^fl/fl^* mice was achieved via oral gavage with tamoxifen, as described above with the following change in dosage: 225mg/kg in 100 μL corn oil /5% ethanol; 3x every second day. Body weight and temperature (Rodent Thermometer BIO-TK8851, Bioseb, France) were monitored from daily to once a week.

### Parabiosis

Parabiosis was performed essentially as described (Kamran *et al*, 2013). Briefly, age and weight-matched male mice were cohoused at least 2 weeks before surgery to ensure harmonious cohabitation. Mice were injected with Meloxicam (2mg/kg, subcutaneous, s.c.) 30 min. prior to surgery. Mice were anaesthetized (intraperitoneal, *i*.*p*.) using ketamine (75 mg/kg) and xylazine (15mg/kg) (∼140µL/mouse, 1:1 vol/vol in sterile 0.9% saline) and placed on a heating pad. The corresponding lateral body parts were shaved and disinfected with Betadine® solution. A longitudinal skin incision was made starting at 0.5 cm above the elbow to 0.5 cm below the knee joint, and the subcutaneous fascia was bluntly dissected to create about 0.5 cm of free skin. The corresponding elbow and knee joints were sutured together using a silk suture (3-0 Mersilk #W212) and the corresponding dorsal and ventral skin were attached using continuous sutures (5-0 Vicryl). Mice were resuscitated with 0.9% saline solution (1mL, s.c.) and placed on a heating pad (30min. -2h) until recovery from anesthesia. Following recovery, each parabiotic pair was placed in a clean cage and provided with free access to food and water by placing hydrogel or food pellets on the bottom of the cage. Mice were injected with analgesics buprenorphine (0.1mg/kg s.c.) every 12 hours for 48 hours and Meloxicam (1mg/kg s.c.) every 24 hours for 48 hours. Mice were monitored for signs of pain, distress and weight loss for 6-8 weeks until at least 80% of the combined original weight was recovered. Tamoxifen food was given for 10 days after which it was replaced by normal food. The survival and weight loss of the parabiotic mouse pairs were monitored for 100 days.

### Adoptive cell transfer

Adoptive transfer of bone marrow-derived monocytes (BMDMo) was performed essentially as described (Wagner *et al*, 2014). Briefly, BMDMo cells were generated from both *Fth*-competent (*tdT^R26^*) and *Fth*-deleted (*tdT^R26^Fth^R2fl/fl^*). To this end, bone marrow was freshly isolated from the tibia and femurs of *tdT^R26^*and *tdT^R26^Fth^R2fl/fl^* mice and placed in culture for 7 days in Ultra low-adherence T75 cell culture flasks (Corning cat. #734-4139) with BMDMo differentiation culture medium (RPMI, 10%FCS, 1%Pen/Strep, supplemented with 10% L929 culture supernatant containing M-CSF1). Concomitantly, *Fth^fl/fl^* and *Fth^R26fl/fl^* mice were treated with tamoxifen as described above to induce the deletion of *Fth*. Differentiated BMDMo were collected from cell culture flasks, washed in RPMI (10ml; 300g, 5min., 4°C) and resuspended in RPMI (final concentration: 20-30×10^6^ cells/ml). On days 4, 8, 12 and 15 post-tamoxifen treatment, 100µL of BMDMo cell suspension was injected retroorbitally in each mouse (2-3×10^6^ cells/mouse/injection).

### qRT-PCR

RNA was isolated from organs using RNeasy Mini Kit (QIAGEN). Briefly, mice were euthanized and transcardially perfused with ice cold PBS. Organs were collected into Eppendorf tubes and snap frozen in liquid nitrogen. RNA was isolated and processed according to the manufacturer instructions. cDNA was transcribed from total RNA with transcriptor first strand cDNA synthesis kit (Roche). Quantitative real-time PCR (qRT-PCR) was performed using 1μg cDNA and Syber Green Master Mix (Applied Biosystems, Foster City, CA, USA) in duplicate on a ABI QuantStudio - 384 Real-Time PCR System (Applied Biosystems) under the following conditions: 95°C/10 min, 40 cycles/95°C/15 s, annealing at 60°C/30 s and elongation 72°C/30 s. Primers were designed using Primer Blast (Ye *et al*, 2012). Primer sequences are available in ***Table 2***:

### Cardiovascular Function

Cardiovascular function was measured using pressure–volume conductance catheter technique (Pacher *et al*, 2008). Briefly, 7 days post-tamoxifen treatment as described above, *Fth^fl/fl^, R26^CreERT2^* and *R26^CreERT2^Fth*^Δ*/*Δ^ mice were anesthetized with isoflurane, tracheotomized and artificially ventilated. Temperature was recorded continuously and kept stable. The apex of the left ventricle was punctured with a 27 G needle using the open chest approach and a pressure-volume conductance catheter (FTS-1912B-8018; Scisense, London, Canada) was inserted in the left ventricle. Mice were stabilized for 3-10 min. Baseline values, values with varying preload caused by inferior vena cava clamps using a blunt forceps and aortic pressures, were recorded with the Scisense pressure-volume control unit FV896B and analyzed using the Labscribe2 (Labscribe, iWorx Systems, USA) software. The machine was calibrated with internal and cuvette calibration, as described (Pacher *et al*., 2008). Bone marrow chimeras were monitored following *Fth* deletion and analyzed when body temperature dropped below 32°C.

### *In Vivo* Luciferase assay

Following deletion of *Fth* via Tamoxifen treatment as described above bone marrow chimeric mice on *OKD48^Luc^* genetic background were monitored daily for luciferase activity. Mice were anesthetized using intraperitoneal Ketamine/Xylasine injection. The abdomen was shaved, and mice received an intravenous injection of luciferin (2mg/mouse in 100µL PBS). Luciferase signal was acquired in a Hamamatsu Aequoria using an electron multiplying CCD (EMCCD) camera with highest sensitivity (255) and maximum gain (5) for 10sec, 30sec, 60sec, 120sec and 240sec. Quantification was performed using Fiji software (ImageJ).

### Iron quantification in organs and plasma

Non-heme iron concentration in heart and livers of bone marrow chimeras was assessed as previously described (Martins *et al*, 2016). Briefly, liver or heart were homogenized 1:5 in PBS and 100µl (equivalent to 20mg tissue) of homogenate or 20µl plasma were hydrolyzed (65°C; overnight) by adding 50µl of 26% HCl, 1,8M trichloroacetic acid. The hydrolyzed samples were then clarified by centrifugation (3.000xg; RT). Clarified samples (60µl) were transferred to a 96-well plate and mixed with 160µl of 3.8M sodium acetate, 575µM bathophenanthroline disulfonic acid and 2.5mM ascorbic acid. Samples were incubated (5min.; RT) and absorbance was measured at λ_540nm_ and iron concentration was calculated using [Fe] = (((A_S_ - A_b_)x V x MW))/((e x l x t)) , where *A_s_* is the sample absorbance, *A_b_* is the blank absorbance, *V* is the reaction volume (0.22), *MW* is the molecular weight of iron (56 g·mol^-1^), *e* is the millimolar absorptivity of bathophenanthroline disulfonic acid (22.14 mM^-1^·cm^-1^), *l* is the path length (0.6 cm) and *t* is the weight of tissue used. In addition, measured concentrations were confirmed using standard iron serial dilutions.

### Western Blot

Proteins were extracted, electrophoresed, and transferred essentially as described (Blankenhaus *et al*., 2019). Briefly, organs were collected from mice following euthanasia and perfusion with ice cold PBS (20mL) and snap frozen in liquid nitrogen. For protein extraction tissue was homogenized in RIPA buffer using a tissue douncer kit, sonicated and centrifuged. Supernatant was collected and total protein was quantified using Bradford assay. Anti-mouse-FTH1 (clone D1D4; 1:1000; Cell Signaling cat. #4393), anti-V5-HRP (1:5000; Invitrogen cat. #46-0708), Anti-GAPDH (1:5000; SICGEN cat. # AB0049-200) and anti-β-actin (1:5000; Sigma cat. #A5441) were detected using peroxidase conjugated secondary antibodies (HRP-conjugated anti-rabbit IgG – SantaCruz Biotechnology sc-2030; HRP-conjugated anti-mouse IgG – SantaCruz Biotechnology sc-2005; HRP-conjugated anti-goat IgG – Thermo Fisher PA1-28664; 1:5000; 1 hour; RT) and developed with ECL western blotting substrate (ThermoFisher Scientific). Detection of proteins of the ETC by Western blot was performed using OxPhos Rodent WB Antibody Cocktail (ThermoFisher cat. #45-8099; 1:1000; 4°C overnight), followed by washing and incubation with secondary HRP-conjugated anti-mouse IgG antibody (1:2000, RT; Cell signalling cat. #7076P2). WB quantification was performed using FiJi (ImageJ) or ImageLab Software (BioRad).

### Serology

Bone marrow chimeras were euthanized using CO_2_ inhalation and blood was collected by cardiac puncture and placed in heparin tubes. Blood samples were sent to DNAtech (Clinical and veterinary analysis laboratory, Lisbon) for analysis of serologic parameters including ALT, AST, Urea, CPK, Troponin I and LDH as well as Transferrin and Transferrin saturation.

### Histology

Organs were harvested, fixed in 10% formalin, embedded in paraffin, sectioned into 3 μm-thick sections and stained with Hematoxylin and Eosin (H&E). Whole sections were analyzed and images acquired with a Leica DMLB2 microscope (Leica) and NanoZoomer-SQ Digital slide scanner (Hamamatsu). For WAT adipocytes area and BAT lipid droplets measurements, H&E-stained paraffin-embedded sections (3μm sections) were scanned into digital images (NanoZoomer-SQ Digital slide scanner - Hamamatsu). The average WAT adipocyte size in adipose tissue sections (expressed as the mean cross-sectional area per cell (μm2)) was determined using Fiji software, as described elsewhere (Blankenhaus *et al*., 2019). Briefly, a slide scanned picture was captured at 2.5x magnification. An average of 1500 adipocytes were measured per sample. The following macro was applied: run (“Set Scale…”, “distance=560 known=250 pixel=1 unit=um global”); run (“Duplicate…”, “ “); run (“Subtract Background…”, “rolling=50 light separate sliding”); run (“Despeckle”); run (“8-bit”); setAutoThreshold(“Mean dark”); //run (“Threshold…”); //setThreshold(250, 255); setOption(“BlackBackground”, false); run (“Convert to Mask”); run (“Make Binary”); run (“Dilate”); run (“Close-”); run (“Invert”); run (“Analyze Particles…”, “size=330-15000 circularity=0.50-1.00 display exclude clear summarize add”). The average BAT lipid droplet size (expressed as the mean cross-sectional area per lipid droplet (μm2)) were quantified in 3 non-overlapping 40x magnification fields for each mouse. The following macro was applied: run (“Set Scale…”, “distance=452 known=100 pixel=1 unit=um global”); run (“Duplicate…”, “ “); run (“Subtract Background…”, “rolling=50 light separate sliding”); run (“Despeckle”); run (“8-bit”); setAutoThreshold(“Mean dark”); //run (“Threshold…”); //setThreshold(245, 255); setOption(“BlackBackground”, false); run (“Convert to Mask”); run (“Make Binary”); run (“Dilate”); run (“Close-”); run (“Invert”); run 25 (“Watershed”); run (“Analyze Particles…”, “size=1-5000 circularity=0.4-1.00 display exclude clear summarize add”).

### Flow cytometry

Reconstitution of bone marrow chimeras was tested by staining peripheral blood leukocytes with CD45.1-FITC (BioLegend cat. #110705), CD45.2-APC (BioLegend cat. #109813), to determine the relative contribution of donor bone marrow towards engraftment. To quantify the numbers of tissue leukocytes, animals were transcardially perfused with 10mL cold PBS. Briefly, liver, kidney, lung and heart were cut into small pieces, digested in 10 mL digestion medium (HBSS supplemented with 1mg/mL Collagenase D (Sigma-Aldrich cat. #11088866001) and 10µg/mL DNase I (Sigma-Aldrich cat. #10104159001)), shaking at 220rpm, 37°C for 45 min. The digested solution was passed through a cell strainer (100µm) and washed with 5 mL of RPMI. Cells were pelleted by centrifugation (300g; 4°C; 10min) and re-suspended in 5 mL ACK red blood cell (RBC) lysis buffer. After 5 min at RT RBC lysis was stopped by adding 5 mL of FACS buffer (1x PBS 3% FCS) and cells were passed through a 40µm cell strainer. Cells were again centrifuged (300g; 4°C; 10min) and finally re-suspended in 3mL (liver and kidney) or 1mL (heart and lung) FACS buffer. 300µL of cell suspension were used for the staining of leukocytes with the following antibodies: Cx3CR1-APC (Biolegend, cat. #149007), CD11b-FITC (BD, cat. #553310), Ly6C-PerCPCy5.5 (Biolegend, cat. #128012), Ly6G-PE (BD, cat. #551461), F4/80-PE-Cy7 (Biolegend, cat. #123114), CD45-APC-e780 (Invitrogen, cat. #47045182), CD3-biotin (BioLegend, cat. #100243), CD11b-BV785 (BioLegend, cat. #101243), CD11c-BV605 (BioLegend, cat. #11733), CD19-biotin (BD, cat. #553784), CD31-BV605 (BioLegend, cat. #102427), CD49b-biotin (BioLegend, cat. #103521), CD90.1-Thy1.1-PE (BioLegend, cat. #202524) F4/80-PE-Cy5 (BioLegend, cat. #123112) Ly6C-PE-Cy7 (eBioscience, cat. #25-5932-8), Ly6G-AF700 (Invitrogen, cat. #56-9668-8), and SAV-PE-Fire-700 (BioLegend, cat. #405174). In addition, Fc-block (in-house anti-CD16/32) was used to minimize unspecific Ig binding and LIVE/DEAD^TM^ Fixable viability dye (ThermoFisher cat. #L34957), and LIVE/DEADTM Fixable Yellow viability dye (ThermoFisher cat. # L34959) were used to assess cell viability. Data acquisition was performed using either CyanADP (Beckman Coulter), BD LSRFortessa X-20 (BD Biosciences) or Aurora (Cytek) flow cytometers. Samples were analyzed using FlowJo software.

### Cell sorting

Monocyte/macrophages were sorted from the liver, WAT and heart of bone marrow chimeras, as indicated in *Fig. S9C*. Briefly, *Fth* deletion was induced via tamoxifen treatment as described above, and 19 days later, bone marrow chimeras were euthanized, and organs were collected. Liver, and heart were cut into small pieces and incubated with digestion buffer (liver: 4ml; heart: 1ml; 0.2mg/mL Liberase (Roche cat. #5401127001) and 100µg/mL DNAse I (Sigma-Aldrich cat. #10104159001). After digestion, cell suspensions passed through a 40µm cell strainer. Liver and heart cells were then pelleted (350g, 5min., 4°C) and resuspended in 40% Percoll (GE healthcare cat. #10607095) in RPMI (liver: 15ml, heart: 6ml). A density gradient was made by overlaying the cell suspension onto 80% Percoll in RPMI (liver: 5ml, heart: 2ml), and centrifuged (700g no brake/lowest acceleration, 20min. RT). Liver and heart cells were collected from the density interface and washed once with FACS buffer (10-15ml; 1x PBS 3%FCS 2mM EDTA) and centrifuged (350g, 5min. 4°C). plate for antibody staining. WAT was cut into small pieces and placed in 3mL WAT digestion buffer (4mg/mL Collagenase IV, 10mM CaCl_2_, 0.5% BSA, in PBS without Ca^2+^/Mg^2+^). After digestion, WAT cell suspensions were passed through a 100µm cell strainer and centrifuged (500g, 10min., 4°C). Liver and heart cell pellets were resuspended in 5mL FACS buffer, and WAT cell pellets were resuspended in 0.25-1mL FACS buffer. Cells were counted using trypan blue and 1∼2×10^6^ cells were placed on a 96-well. Cells were centrifuged (700g, 2min. 4°C), resuspended in 150µL of ACK buffer for red blood cell lysis and incubated (5min, RT). Lysis was stopped by adding 50µL FACS buffer and cells were washed (700g, 2min. 4°C). Cells were resuspended in 50µL FACS buffer and antibody staining was performed using: CD11b-BV421 (BioLegend, cat. #101235), Ly6G-AF700 (Invitrogen, cat. #56-9668-82), CD45.2-APC (BioLegend cat. #109813), TCRb-FITC (In-house) and CD19-FITC (In-house). In addition, Fc-block (in-house anti-CD16/32) was used to minimize unspecific Ig binding and Zombie aqua viability dye (BioLegend cat. #423101) was used to assess cell viability. Cell sorting and data acquisition was performed using BD FACSAria II (BD Biosciences) cell sorter. 500 cells/sample were sorted directly into 0.2mL Eppendorf tubes containing 2.5µL Buffer RLT plus (Qiagen, cat. #1053393). Flow cytometry data were analyzed using FlowJo software.

### Bulk RNA sequencing

RNA was extracted, cleaned (RNeasy MinElute Cleanup Kit, Qiagen) and its quality assessed using an Agilent Bioanalyzer 2100 (Agilent Technologies) together with an RNA 6000 pico kit (Agilent Technologies). Full-length cDNAs and sequencing libraries were generated according to the SMART-Seq2 protocol, as previously described (Ramos *et al*, 2022). Library preparation including cDNA ‘tagmentation’, PCR-mediated adaptor addition and amplification of the adapted libraries was done following the Nextera library preparation protocol (llumina Tagment DNA Enzyme and Buffer, Illumina #20034211; KAPA HiFi HotStart ReadyMix, Roche #07958935001; Nextera XT Index Kit v2 Set A, Illumina #15052163; Nextera XT Index Kit v2 Set D, Illumina #15052166), as previously described (Ramos *et al*., 2022). Libraries were sequenced (NextSeq500 sequencing; Illumina) using 75 SE high throughput kit. Sequence information was extracted in FastQ format, using Illumina’s bcl2fastq v.2.19.1.403, producing on average approximately 38×10^6^ (liver), 32×10^6^ (WAT) and 43×10^6^ (heart) reads per sample. Library preparation and sequencing were optimized and performed at the Gulbenkian Institute for Molecular Medicine Genomics Unit.

FastQ reads were aligned against the mouse reference genome GRCm39 using the GENCODE vM27 annotation to extract splice junction information (STAR; v.2.5.2a) (Dobin *et al*, 2013). Read summarization was performed by assigning uniquely mapped reads to genomic features using *FeatureCounts* (v.1.5.0-p1)(Liao *et al*, 2014). Gene expression tables were imported into the R programming language and environment (v.4.1.0) to perform differential gene expression and functional enrichment analyses, as well as data visualization.

Differential gene expression was performed using the DESeq2 R package (v.1.32) (Love *et al*, 2014). Gene expression was modeled by genotype for each organ, which included the following factors: *tdT^LysM^*⇨*Fth^fl/fl^*, *tdT^LysM^Fth^LysM^*^Δ*/*Δ^⇨*Fth^fl/fl^* or *tdT^LysM^Fth^LysM^*^Δ*/*Δ^⇨*Fth^R26^*^Δ*/*Δ^ bone marrow chimeras. Genes not expressed or with fewer than 10 counts across the samples were removed, leaving 22,529 (liver), 23,327 (WAT) and 29,821 (heart) genes for downstream differential gene expression analysis. We subsequently ran the function *DESeq* to estimate the size factors (by *estimateSizeFactors*), dispersion (by *estimateDispersions*) and fit a binomial GLM fitting for βi coefficient and Wald statistics (by *nbinomWaldTest*). Pairwise comparisons tested with the function *results* (alpha = 0.05), were: 1) *tdT^LysM^*⇨*Fth^fl/fl^*, *vs*. *tdT^LysM^Fth^LysM^*^Δ*/*Δ^⇨*Fth^fl/f^l*; 2) *tdT^LysM^*⇨*Fth^fl/f^l* vs. *tdT^LysM^Fth^LysM^*^Δ*/*Δ^⇨*Fth^R26^*^Δ*/*Δ^ and 3) *tdT^LysM^Fth^LysM^*^Δ*/*Δ^⇨*Fth^fl/fl^* vs. *tdT^LysM^Fth^LysM^*^Δ*/*Δ^⇨*Fth^R26^*^Δ*/*Δ^. In addition, the log_2_ fold change for each pairwise comparison was shrunken with the function *lfcShrink* using the algorithm *ashr* (v.2.2– 47)(Stephens, 2016). Differentially expressed genes were considered for genes with an adjusted p value<0.05 and an absolute log_2_ fold change>0. Normalized gene expression counts were obtained with the function *counts* using the option normalized = TRUE. Regularized log transformed gene expression counts were obtained with *rlog*, using the option blind = TRUE. Ensembl gene ids were converted into gene symbols from Ensembl (v.107) by using the mouse reference (GRCm39) database with biomaRt R package (v.2.48.2)(Durinck *et al*, 2005; Durinck *et al*, 2009). All scatterplots, including volcano plots, were done with the ggplot2 R package (v.3.3.5)(Wickham, 2016). Functional enrichment analysis was performed with the gprofiler2 R package (v.0.2.1)(Kolberg *et al*, 2020). Enrichment was performed using the function *gost* based on the list of up- or down-regulated genes (genes with an adjusted p value<0.05 and a log_2_ fold-change>0 or <0), between each pairwise comparison (independently), against annotated genes (domain_scope = “annotated”) of the organism *Mus musculus* (organism = “mmusculus”). Gene lists were sorted according to adjusted p value (ordered_query = TRUE) to generate GSEA (Gene Set Enrichment Analysis) style p values. Only statistically significant (user_threshold = 0.05) enriched functions are returned (significant = TRUE) after multiple testing corrections with the default method g:SCS (correction_method = “analytical”). The gprofiler2 queries were run against all the default functional databases for mouse which include: Gene Ontology (GO:MF, GO:BP, GO:CC), KEGG (KEGG), Reactome (REAC), TRANSFAC (TF), miRTarBase (MIRNA), Human phenotype ontology (HP), WikiPathways (WP), and CORUM (CORUM). For future reference, gprofiler2 was performed using database versions Ensembl 107, Ensembl genomes 54 (database updated on 12/07/2022). For STRING database network analysis, genes contained within enriched gene sets associated related to mitochondrial proteins, mitochondrial ribosome, electron transport chain, respirasome were merged and uploaded to the STRING database (v11.5) (Shannon *et al*, 2003) and queried for known protein-protein interactions (organism: Mus musculus; interaction score >0.4). The resulting network was imported into Cytoscape (v3.9.0) (Shannon *et al*., 2003) for network layout design.

### Thermal imaging

BAT and tail temperatures were measured in mice that were allowed to move freely in a cage, using an infrared camera, (FLIR E96 Compact-Infrared-Thermal-Imaging-Camera; FLIR Systems). At least 2 days prior, mice were anesthetized (1-2% Isoflurane) and the interscapular area was shaved. Acquired images were analyzed using FLIR Tools software (v6.4) and individual BAT and tail temperatures were taken as the maximum temperature measured at the interscapular area or tail base, respectively.

### Metabolic phenotyping

Promethion Core (Sable Systems, USA) was used to measure indirect calorimetry. *Fth* deletion was induced via tamoxifen administration as described above and 7 (early onset) or 20 (late onset) days post-tamoxifen treatment, bone marrow chimeras were placed in metabolic cages for metabolic phenotyping. Mice were kept on a 14/10 h light/dark cycle with controlled temperature and humidity. Recording continued for the following 5-6 days. The system consists of a standard GM-500 cage with a food hopper and a water bottle connected to load cells (2 mg precision) with 1 Hz rate data collection. Additionally, the cage contains a red house enrichment. Ambulatory activity was monitored at 1 Hz rate using an XY beam break array (1 cm spacing). Oxygen, carbon dioxide and water vapor were measured using a CGF unit (Sable Systems). This multiplexed system operated in pull-mode. Air flow was measured and controlled by the CGF (Sable Systems) with a set flow rate of 2LJL/min. Oxygen consumption and carbon dioxide production were reported in milliliters per minute (mL/min). Energy expenditure was calculated using the Weir equation and Respiratory Exchange Ratio (RER) was calculated as the ratio of VCO_2_/VO_2_. Raw data was processed using Macro Interpreter v2.41(Sable Systems), as described (Ramos *et al*., 2022).

### Fluorescence microscopy

Bone marrow chimeras were treated with tamoxifen to induce *Fth* deletion, as described above. 21 days post tamoxifen treatment, bone marrow chimeras were euthanized and transcardially perfused with 20mL cold PBS, followed by prefusion with 10mL 4%PFA (Alfa Aesar cat. #043368-9M) in PBS. Livers were collected and fixed in 4%PFA in PBS overnight in the dark. Organs were then washed in PBS overnight and embedded in 4% low-melting point agarose (Invitrogen cat. #15517-014). Tissue sections (100µm) were obtained using a Leica Vibratome VT 1000 S (Leica Biosystems) and placed in Eppendorf tubes with PBDO permeabilization solution (1% Bovine Serum Albumin; 1% DMSO; 0.6 % Triton X-100 in PBS; overnight; 4°C). Tissue slices were then incubated with primary Abs (overnight; 4°C): anti-FTH (1:100, Cell Signalling cat. #4393), Anti-V5 (1:200, Abcam cat. # AB9137) and Anti-CD68 (1:200, BioLegend cat. #123102). Tissue slices were further washed with PBDO (overnight; 4°C) and were then incubated (1:500; overnight; 4°C) with secondary antibodies: Cy5-conjugated donkey anti-goat IgG antibody (Jackson IR; cat. #705-175-147), Alexa 568 conjugated anti-rabbit (ThermoFisher, cat. #A-11011) and Alexa 568 conjugated anti-rat (ThermoFisher, cat. #A-11077). Tissue slices were washed once more with PBDO (overnight, 4°C) and mounted with Mowiol mounting medium with 10ug/mL of DAPI (Merck, cat. #81381-50G). Images were then acquired on a Leica SP5 confocal-based on a Leica DM6000 inverted microscope (458, 476, 488, 514, 561 and 633 nm lasers).

### Transmission electron microscopy

Animals were euthanized and perfused with fixating media (2% formaldehyde (EMS), 2.5% glutaraldehyde (Polysciences) in 0.1M Phosphate Buffer (PB); pH 7.4). Organs were dissected and immersed in the same primary fixative (1h; RT). Further processing was achieved using a PELCO BioWave Microwave Processor at 23 °C, restricted by a PELCO SteadyTemp Pro. Samples were additionally fixed in the primary fixative using a time-sequence of 7 × 2 min with ON and OFF sequential cycles of 0 and 100 W irradiating power in vacuum and rinsed with PB before post-fixation in 1% (v/v) osmium tetroxide (®EMS) with 1% (w/v) potassium ferrocyanide (®Sigma Aldrich) in PB for 8 × 2 min also with ON and OFF sequential cycles of 100 W in vacuum. Subsequently, samples were washed with PB and dH_2_O twice and immersed in 1% (w/v) tannic acid (®EMS) followed by *en-bloc* staining with 0.5% (w/v) uranyl acetate. Both steps were made using a time-sequence of 7 × 2 min with ON and OFF sequential cycles of 0 and 150 W irradiating power in vacuum. Between the steps, samples were rinsed with dH_2_O. Dehydration was done in a graded ethanol series of 30%, 50%, 75%, 90% and 100%, for 40 s at 150 W each. EPON resin (®EMS), 25%, 50%, 75% and 100% was infiltrated, for 3 min at 250 W in vacuum each step, and cured Over-Night, at 60 °C. Sections of 70 nm were obtained on a Leica UC7 and mounted on palladium-copper grids coated with 1% (w/v) formvar (®Agar Scientific) in chloroform (®VWR). Sections were stained with 1% (w/v) uranyl acetate and Reynolds lead citrate for 5 min each and imaged on an FEI Tecnai G2 Spirit BioTWIN Transmission Electron Microscope operating at 120keV.

### Mitochondria quantification

Livers, WAT and heart were harvested and snap frozen as described above. A piece of the tissue was cut and placed into lysis buffer (100 mM NaCl, 10 mM EDTA, 0.5% SDS and 20 mM Tris-HCl, pH 7.4) and homogenized in a Qiagen TissueLyser II using tungsten carbide beads. Upon homogenization, an equal volume of Phenol:Chloroform:Isoamyl Alcohol (25:24:1, v/v) was added. DNA, present in the aqueous phase, was precipitated using 1vol. isopropanol and 0.3M sodium acetate for 3h at −20 °C. The isolated DNA was used to perform the quantification of mitochondrial DNA (mtDNA) as compared to nuclear DNA (nDNA) using a qRT-PCR -based method, similar to what was previously described (Blankenhaus *et al*., 2019; Quiros *et al*, 2017). Briefly, qRT-PCR was performed using 20 ng of DNA and SYBR Green Master Mix (Applied Biosystems, Foster City, CA, USA), in duplicate on a ABI QuantStudio - 384 Real-Time PCR System (Applied Biosystems), under the following conditions: 50 °C/2min and 95°C/5.min (Hold stage), 45 cycles/95 °C/10 s, annealing at 60 °C/30 s, and elongation 72 °C/20 s, followed by melting curve: 95 °C for 15 s, 60 °C for 1 min, and gradual increase in temperature up to 95 °C. Primers for NADH-ubiquinone oxidoreductase chain 1 encoded by the mitochondrial gene MT-Nd1 (Nd1) and for the nuclear encoded hexokinase 2 gene (Hk2) (Blankenhaus *et al*., 2019; Quiros *et al*., 2017) are available in *Table 1*. Mitochondria number per cell was calculated by the ratio of mRNA expression of the single copy mitochondrial gene *Nd1* and the single copy nuclear gene *Hk2*.

**Table 1:** Mouse strains.

|  |  |  |
| --- | --- | --- |
| *Fth<sup>fl/fl</sup>* (C57BL/6) | Prof. Lukas Kuhn (ETH, Switzerland) | N/A |
| *R26<sup>CreERT2</sup>* | The Jackson Laboratory | Strain: 008463 |
| *Cx3cr1<sup>Cre</sup>* | The Jackson Laboratory | Strain: 025524 |
| *LysM<sup>Cre</sup>* | The Jackson Laboratory | Strain: 004781 |
| *CD2<sup>Cre</sup>* | The Jackson Laboratory | Strain: 027406 |
| *OKD48<sup>Luc</sup>* | (Oikawa *et al.*, 2012) | Prof. Iwawaki T. (RIKEN 2-1, Wako, Japan) |
| *OKD48<sup>Luc</sup> Fth<sup>fl/fl</sup>* | (Blankenhaus *et al.*, 2019) | N/A |
| *OKD48<sup>Luc</sup> R26<sup>CreERT2</sup>* | (Blankenhaus *et al.*, 2019) | N/A |
| *OKD48<sup>Luc</sup> R26<sup>CreERT2</sup> Fth<sup>Δ/Δ</sup>* | (Blankenhaus *et al.*, 2019) | N/A |
| *R26<sup>CreERT2</sup> Fth<sup>Δ/Δ</sup>* | This manuscript | N/A |
| *LysM<sup>Cre</sup> Fth<sup>Δ/Δ</sup>* | This manuscript | N/A |
| *CD2<sup>Cre</sup> Fth<sup>Δ/Δ</sup>* | This manuscript | N/A |
| *Cx3CR1<sup>Cre</sup> Fth<sup>Δ/Δ</sup>* | This manuscript | N/A |
| C57BL/6 Ly5.1 | The Jackson Laboratory | Strain: 002014 |
| *Ccr2<sup>-/-</sup>* | The Jackson Laboratory | Strain: 004999 |
| B6.129S *Tfrc<sup>fl/fl</sup>* | The Jackson Laboratory | Strain: 028363 |
| C57BL/6 *Slc40a1<sup>fl/fl</sup>* | (Wu *et al.*, 2023) | N/A |
| C57BL/6 *Tfam<sup>fl/fl</sup>* | (Larsson *et al.*, 1998) | MGI: 1860962 |
| C57BL/6 *Uqcqr<sup>fl/fl</sup>* | (Weinberg *et al.*, 2019) | Referred to as *CIII<sup>fl/fl</sup>*; MGI: 6358198 |
| *LysM<sup>Cre</sup> Tfrc<sup>Δ/Δ</sup>* | This manuscript | N/A |
| *LysM<sup>Cre</sup> Slc40a1<sup>fl/fl</sup>* | This manuscript | N/A |
| *LysM<sup>Cre</sup> Tfam<sup>fl/fl</sup>* | Prof. David Sancho, CNIC, Spain | N/A |
| *LysM<sup>Cre</sup> Uqcqr<sup>fl/fl</sup>* | Prof. David Sancho, CNIC, Spain | N/A |
| *R26<sup>tdTomato</sup>* | The Jackson Laboratory | Strain: 007909 |
| *R26<sup>tdTomato/CreERT2</sup>* | This manuscript | N/A |
| *R26<sup>tdTomato/CreERT2</sup> Fth<sup>Δ/Δ</sup>* | This manuscript | N/A |
| *LysM<sup>Cre</sup> R26<sup>tdTomato</sup>* | This manuscript | N/A |
| *LysM<sup>Cre</sup> R26<sup>tdTomato</sup> Fth<sup>Δ/Δ</sup>* | This manuscript | N/A |
| *Fth<sup>V5/V5</sup>* | This manuscript | N/A |
| *R26<sup>PhAM</sup>* | The Jackson Laboratory | Referred to as *PhAM<sup>fl/fl</sup>*; Strain: 018385 |
| *R26<sup>PhAM/CreERT2</sup>* | This manuscript | N/A |
| *R26<sup>PhAM/CreERT2</sup> Fth<sup>Δ/Δ</sup>* | This manuscript | N/A |
| *LysM<sup>Cre</sup> R26<sup>PhAM</sup>* | This manuscript | N/A |
| *LysM<sup>Cre</sup> R26<sup>PhAM</sup> Fth<sup>Δ/Δ</sup>* | This manuscript | N/A |

**Table 2:**
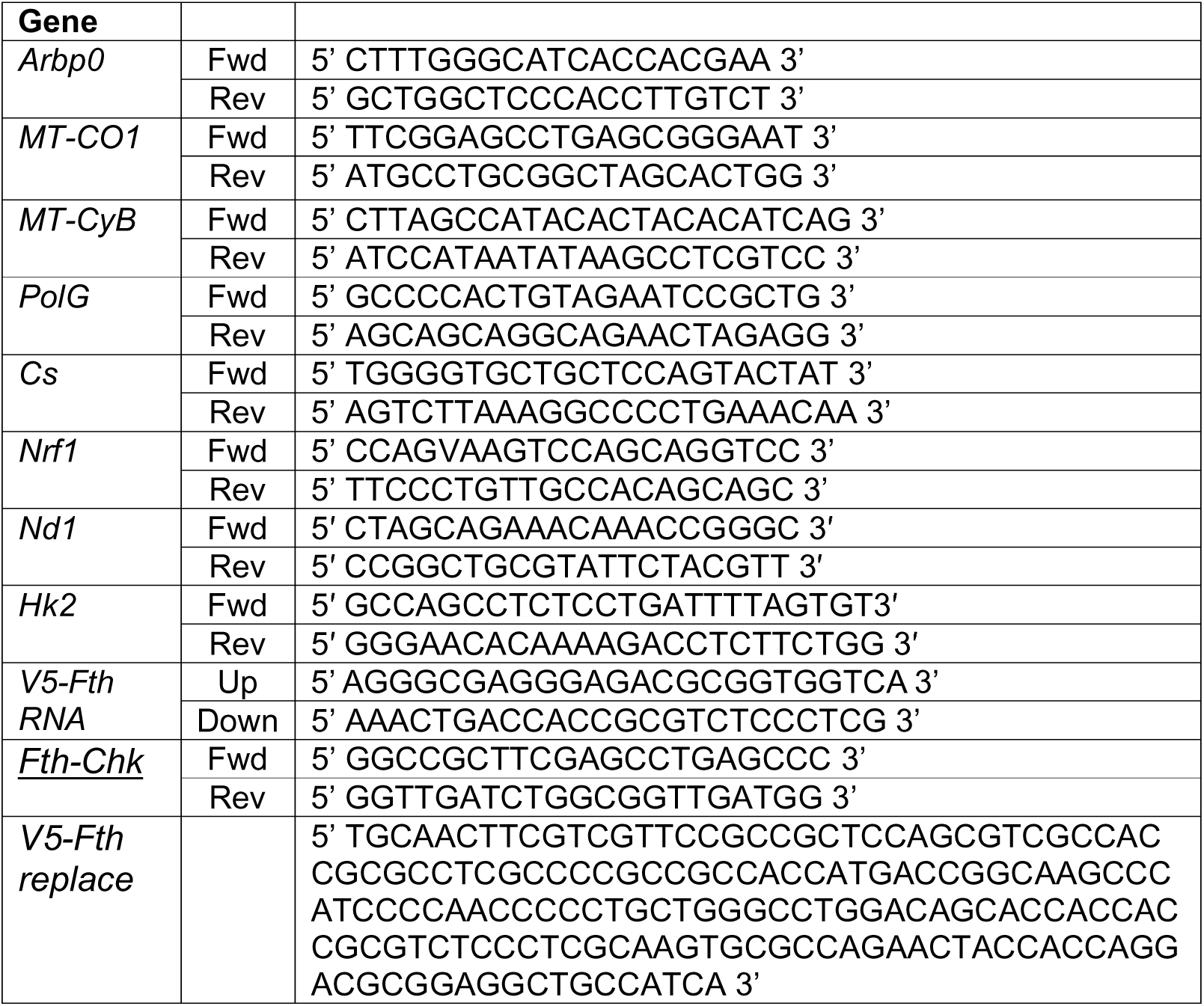
Primer sequences.

### Statistical analysis

Statistical analysis was conducted using GraphPad Prism v8.4.2 software. All data are displayed as means ± standard deviation of the mean (SD) unless otherwise noted. Statistical comparison between two groups was performed using either a student’s T test or Mann–Whitney U test. Groups of three or more were analyzed by one-way analysis of variance (ANOVA) or the Kruskal–Wallis test, using Tukey’s range test or Dunn’s test for multiple comparison correction, respectively. Survival was assessed using a log-rank (Mantel–Cox) test. Statistical parameters for each experiment can be found within the corresponding figure legends.

## Notes

### Competing Interest Statement

The authors have declared no competing interest.

